# The influence of HIV sense and antisense transcripts on stochastic HIV transcription and reactivation

**DOI:** 10.1101/2025.05.07.652610

**Authors:** Kamil Więcek, Janusz Wiśniewski, Heng-Chang Chen

## Abstract

In this study, we established dozens of single provirus-infected cellular clones offering various transcriptional phenotypes of HIV. We proposed that stochastic fluctuations in HIV transcription can appear at, at least, two levels: (1) the chromosomal landscape and (2) the in situ HIV integration site. In the former case, proviruses integrating at different genomic locations demonstrated a variety of transcriptional bursting and can be classified in noise space constructed based on the parameters associated with the coefficient of variation and using a mathematical model fitting a curve of exponential decay. In the latter case, stochastic HIV transcription can be unveiled through its phenotypic bifurcation and tended to be a pure epigenetic phenomenon: the identical provirus demonstrated fluctuations in its transcription with an elevated frequency. We observed similar expression patterns between sense and antisense RNA transcripts. Notably, both HIV long terminal repeats reacted to drug stimulation and may reveal distinct behaviors. Overall, our data suggest that HIV antisense transcripts could be involved in the stochastic nature of HIV transcription.

## 1 Introduction

Noise in gene expression can be generated at multiple levels, ranging from transcription and translation to chromatin remodeling and its associated pathway, as well as gene-specific factors [1–7]. Studying such noise thus aids in a better understanding of how genes are regulated and their biological functions.

Human immunodeficiency virus (HIV) possesses two promoters, the 5’ and 3’ long terminal repeat (5’LTR and 3’LTR) promoters, and its transcription is an inherently stochastic process, generating strong fluctuations in HIV gene products [8–10]. Such stochastic viral gene transcription [11–17] and translation [18–20] can persist during the administration of antiretroviral therapy (ART), rendering HIV reservoirs most likely transcriptionally heterogeneous.

At the chromosome level, the heterogeneity of HIV transcription has been observed from proviruses that integrate at different genomic locations associated with various epigenetic features, demonstrating inconsistency in HIV transcription [21–26]. At the virus level, HIV transcription has a long and rich history of investigations and is well known to be tightly regulated by the 5’LTR promoter [27–29], with which HIV hijacks host cellular transcription factors to control its gene expression [30]. One well-studied mechanism is the impact of RNA polymerase II (RNAPII) elongation on HIV transcription [27–29]; another is the effect of chromatin remodeling and nucleosome positioning [31,32]. Either way, evidence explaining how some viruses remain active and others turn into latency, reflecting the stochastic phenotype of HIV transcription, remains lacking.

Although it is currently unknown whether the HIV Tat positive-feedback loop is the only intrinsic molecular mechanism that governs the provirus transcriptional state (i.e., 5’LTR in the ON versus the OFF state), at present, the two-state promoter model (ON versus OFF) remains the most popular model for representing HIV transcriptional noise, an episodic process of gene expression [10,33] and explaining the molecular mechanism behind the bursty phenotype of HIV transcription. This two-state model relies on a specific regulatory structure to determine the thresholds that allow a biological system to either respond to or disregard an input signal. The Tat positive-feedback circuit (the presence and absence of Tat) is sufficient to induce a burst of HIV 5’LTR promoter activity that leads to fluctuations in HIV transcription [34]; however, this model fails to generate a clear threshold response with deterministic bistability [35], implying that either the two-state model for the HIV 5’LTR promoter is atypical with a transient threshold [35,36], or an additional unknown determinant is involved in this circuit. Indeed, in physiological HIV infection conditions, latent HIV infections can be established at the early stage of infection [37–39], indicating that perhaps an additional molecular determinant that functions as an antagonist to the HIV Tat transactivator is most likely concomitantly present when the Tat positive-feedback circuit is initiated.

A recent study (preprint) suggested that, although the expression of HIV antisense transcripts (ASTs) could be influenced by Tat-dependent sense HIV transcription based on *in vitro* cellular models, the antisense RNA synthesis and the expression remain detectable in cells infected with Tat-deficient viruses. This finding, which suggests that most likely the expression of HIV ASTs is Tat-independent [40], aligns with the previous findings reported by Bentley et al. (2004) [41]. A profound mechanism behind the regulation between HIV ASTs and Tat, still requires further investigation. Antisense RNAs are a type of non-coding RNA and were discovered by Singer et al. in 1963 [42]. Different endogenous antisense RNAs have been identified in mammalian cells [43]. In 1988, the existence of HIV ASTs and the encoded protein was first postulated using computer analyses [44] and confirmed more than ten years later [45–48]. To date, HIV is well established as being able to perform antisense transcription from the 3’LTR of its proviral genome [49,50]. HIV ASTs most likely encode at least one major gene product, named HIV antisense protein (ASP), 189 amino acids in length [51]. Like many other antisense RNAs discovered in animals and plants, HIV antisense RNAs repress the expression of their sense RNAs [52–54] and most likely contribute to maintaining proviruses in latency [54].

Although, at present, very little is known about the involvement of HIV ASTs in its pathogenesis, one recent study (preprint) provided evidence that the expression of HIV ASTs is associated with the efficiency of HIV integration [40]. It was observed that the expression of HIV ASTs showed an increase, while the interaction between HIV integrase and the host-factor lens epithelium-derived growth factor (LEDGF)/p75 was impaired [40]. This finding suggests that the expression of HIV ASTs might be tightly connected to HIV DNA integration, a critical step for HIV persistence infection. Furthermore, the expression of HIV ASP is observed to be restricted to the group M HIV type 1 [55], which is responsible for the human pandemic. These findings imply that HIV ASTs are most likely a functional and pathogenic mechanism left behind after a long period of evolutionary selection and might be critical in fulfilling its pathogenesis. Li et al. [56] showed that the ectopic expression of ASTs in CD4+ T cells from people living with HIV on ART blocks latency reversal in response to pharmacologic and T-cell receptor stimulation, suggesting the functional role of HIV ASTs in enforcing transcriptional silencing. Understanding how HIV ASTs interact with their sense RNA transcription, subsequently governing the fluctuation of HIV transcription may be beneficial to gaining better insights into HIV entry and relief from its latency.

Given that the amount of HIV ASTs in bulk infected cells is much lower than that of the sense RNA transcripts (from 1/100 to 1/2500 compared with sense RNAs) [48,53], quantitatively investigating HIV ASTs remains challenging. For this reason, in this study, we attempted to establish dozens of cellular models harboring a single provirus demonstrating various transcriptional phenotypes (**Fig. 1A** and **2A**, n = 33), allowing for better characterization of the interplay between sense and antisense transcripts at the single-virus level. Such cellular models were named “single provirus-integrated cellular clones” (hereinafter, sinpro clones). We first utilized parameters measured via flow cytometry to define noise spaces [10], enabling the classification of sinpro clones harboring proviruses that demonstrated different transcriptional phenotypes. We further performed strand-specific reverse transcription (RT) followed by either quantitative PCR (qPCR) or digital PCR (dPCR) to track the dynamics of sense and antisense RNA transcription independently and provided the potential threshold levels of sense and antisense RNAs that determine stochastic fluctuations in HIV transcription across sinpro clones. We proposed mathematical models to fit curves of exponential decay, enabling better discrimination of distinct intensities of transcription and transcriptional noise in proviruses across sinpro clones. Finally, both HIV LTRs between sinpro clones in different groups reacted to drug (phorbol 12-myristate 13-acetate, PMA plus ionomycin or ingenol-3-angelate, PEP005) stimulation.

**Fig. 1.**
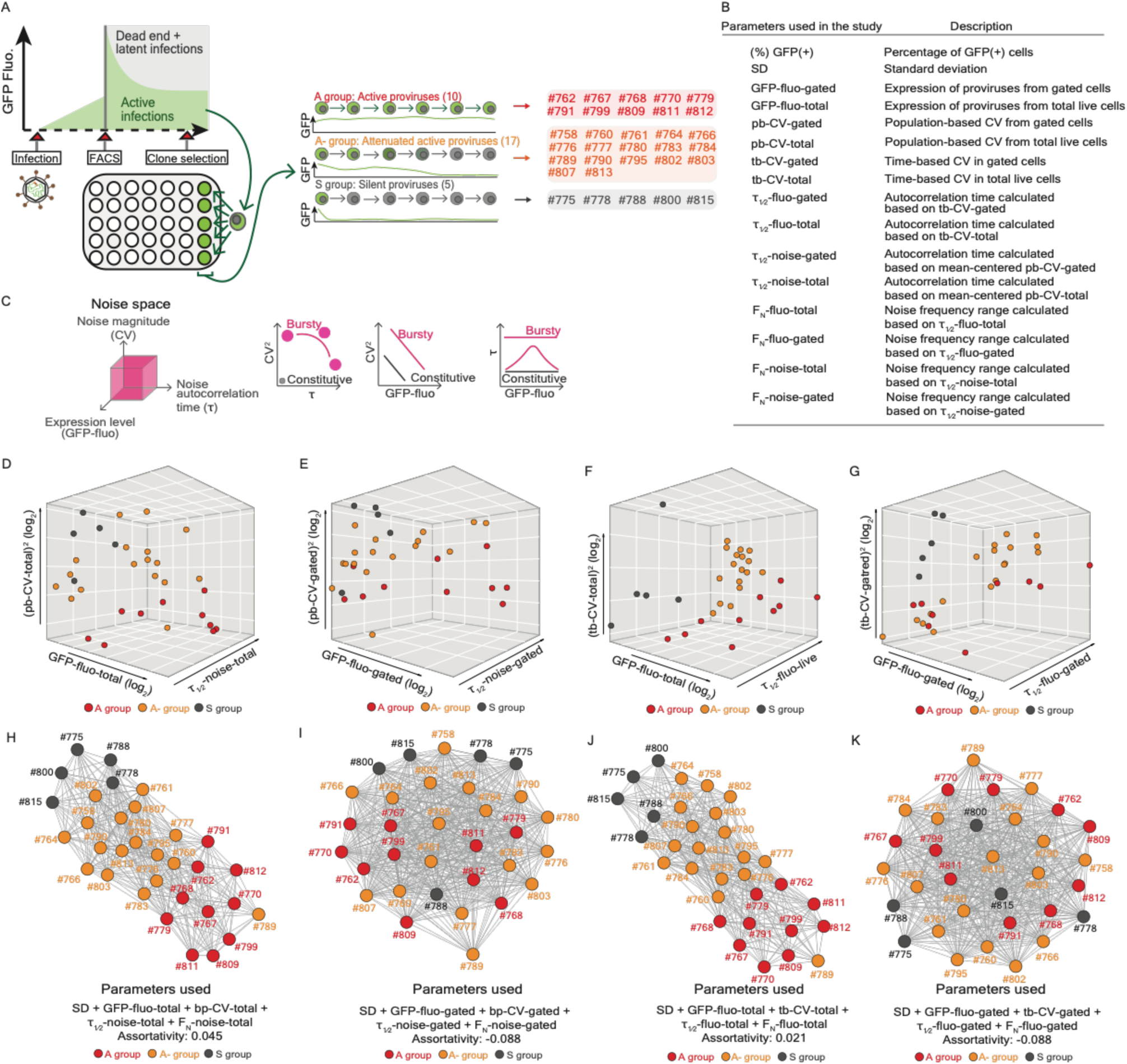
Characteristics of sinpro clones in the A, A-, and S groups. (**A**) Schematic representation of the experimental strategy of the establishment of single provirus-integrated cellular clones (sinpro clones). A total of 33 clones, in which proviruses demonstrated different transcriptional phenotypes were assigned to four different groups, including the active group (the A group, 10 clones), the attenuated active group (the A-group, 17 clones), the silent group (the S group, 5 clones) and the fluctuation group (the F group, 1 clone; see Fig. 2A). (**B**) Table listing parameters used in this work. Detailed descriptions, resources, and calculations of parameters are provided where appropriate in the main text and in the **3. Theory/calculation**. (**C**) Schematic representation of the rationale of noise space previously reported by Dar et al. (2012). Three-dimensional (3D) noise space consists of noise magnitude represented by coefficient of variation (CV), the mean GFP expression, and noise autocorrelation time. Two-dimensional projections on the right-hand side of 3D noise space illustrate different behaviors between bursty (marked in magenta) and constitutive (marked in grey) gene expression. (**D**-**G**) Noise space demonstrating a spatial distribution of sinpro clones in the A, A-, and S groups. Space is independently constructed using four different sets of parameters, including (**D**) population-based (pb)-CV-total, GFP-fluo-total, and τ_1/2_-noise-total; (**E**) pb-CV-gated, GFP-fluo-gated, and τ_1/2_-noise-gated; (**F**) time-based (tb)-CV-total, GFP-fluo-total, and τ_1/2_-fluo-total; and (**G**) tb-CV-gated, GFP-fluo-gated, and τ_1/2_-fluo-gated. Spots marked in red, orange, and grey represent sinpro clones in the A, A-, and S groups, respectively. (**H**-**K**) Graph networks illustrating the topological distribution of sinpro clones in the A, A-, and S groups in the network property. Once again, network properties are structured using four different sets of parameters mentioned above and correspond to respective noise space. Feature parameters used to construct corresponding networks and assortativity coefficients are placed beneath graph networks. Spots marked in red, orange, and grey represent sinpro clones in the A, A-, and S groups, respectively. See also **Fig. S1** and **Tables S1** and **S2**.

**Fig. 2.**
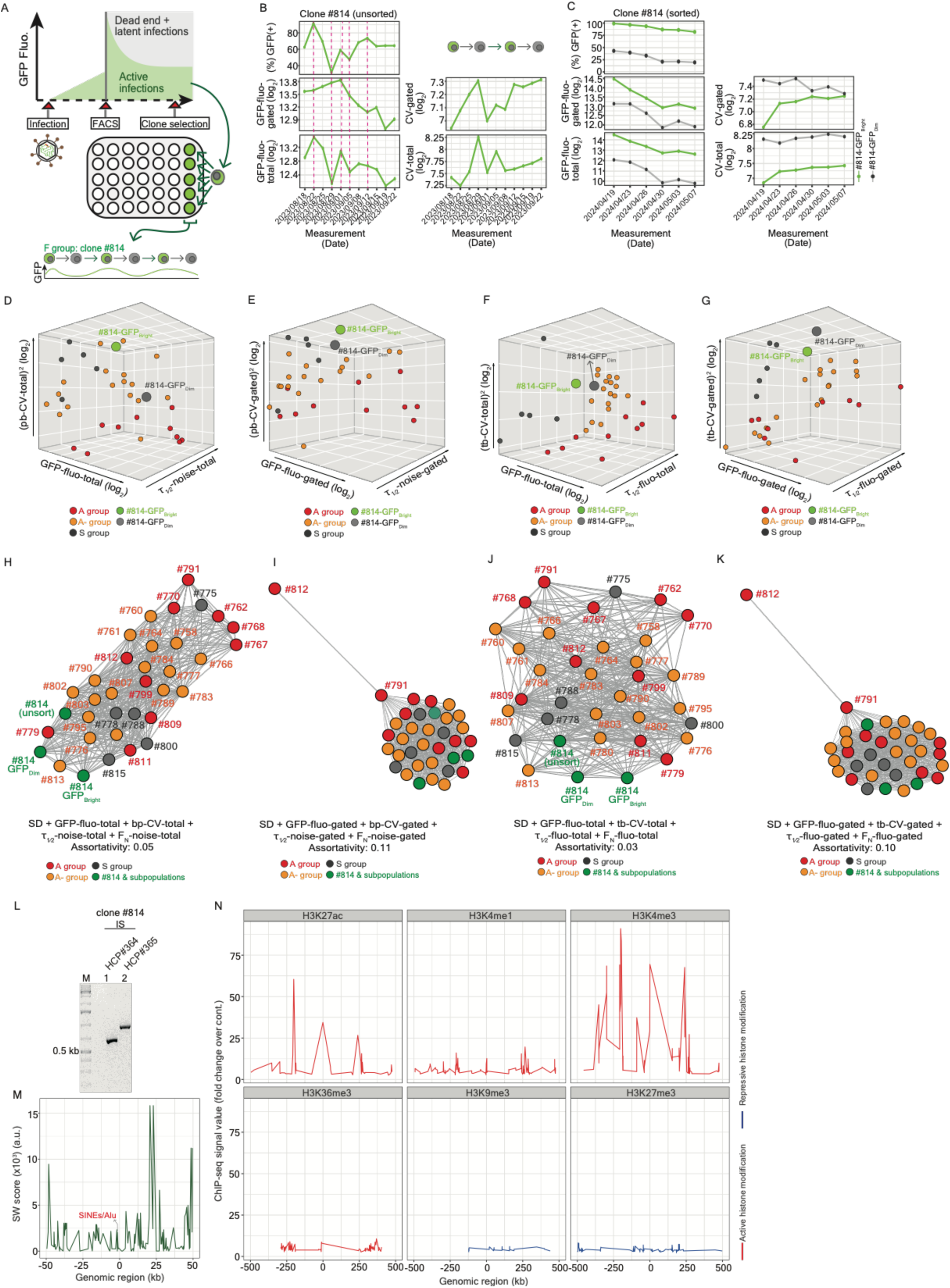
Characteristics of sinpro clone #814. (**A**) Schematic representation of the experimental strategy of the establishment of clone #814, in which the turnover of the state of HIV transcription showed elevated frequency. (**B**) Line plots illustrating the dynamics of measures of parameters, including (%) GFP(+), GFP-fluo-gated, GFP-fluo-total, CV-gated, and CV-total recorded by flow cytometry with unsorted clone #814 at 11 subsequent time points. (**C**) Line plots illustrating the dynamics of measures of parameters, including (%) GFP(+), GFP-fluo-gated, GFP-fluo-total, CV-gated, and CV-total recorded by flow cytometry with clone #814-sorted GFP_Bright_ and GFP_Dim_ subpopulations at six subsequent time points. (**D**-**G**) Noise space demonstrating a spatial distribution of sinpro clones in the A, A-, and S groups as well as clone #814-sorted GFP_Bright_ and GFP_Dim_ subpopulations. Space is independently constructed using four different sets of parameters, including (**D**) population-based (pb)-CV-total, GFP-fluo-total, and τ_1/2_-noise-total; (**E**) pb-CV-gated, GFP-fluo-gated, and τ_1/2_-noise-gated; (**F**) time-based (tb)-CV-total, GFP-fluo-total, and τ_1/2_-fluo-total; and (**G**) tb-CV-gated, GFP-fluo-gated, and τ_1/2_-fluo-gated. Spots marked in red, orange, dark grey, light green, and light grey represent sinpro clones in the A, A-, S groups as well as clone #814-sorted GFP_Bright_ and GFP_Dim_ subpopulations, respectively. (**H**-**K**) Graphic networks illustrating the topological distribution of sinpro clones in the A, A-, and S groups, clone #814 as well as its two subpopulations in the network property. Once again, network properties are structured using four different sets of parameters mentioned above and correspond to the respective noise space. Feature parameters used to construct corresponding networks and assortativity coefficients are placed beneath graph networks. Spots marked in red, orange, and grey represent sinpro clones in the A, A-, and S groups, respectively. Spots marked in green include clone #814 and its two subpopulations. (**L**) Expected results from PCR amplification confirming the provirus integration site identified using LIS-seq. 2.0% (w/v) agarose gel displaying a PCR product obtained from PCR amplification using a pair of primers flanking the junction between the HIV integration site and the host genome (lane 1, HCP#364: 560 bp; lane 2, HCP#365: 650 bp). (**M**) Line plot illustrating the signal profile of human repetitive regions. Smith Waterman (SW) alignment score representing the peak intensity is plotted on the y-axis in the genomic region of 50 kb upstream and downstream from the provirus integration site (chr13:110,913,078) (x-axis) in clone #814. (**N**) Line plot illustrating ChIP-seq signal profiles in the genomic region of 500 kb upstream and downstream from the provirus integration site (chr13:110,913,078) (x-axis) in clone #814. ChIP-seq signal value (fold change over control) represents the intensity of signal peaks shown on the y-axis. Red lines, including H3K27ac, H3K4me1, H3K4me3, and HeK36me3, represent active histone marks, whereas blue lines, including H3K9me3 and H3K27me3, represent repressive histone marks. See also **Fig. S2** and **S3** and **Tables S1**-**S3**.

## 2 Material and methods

### 2.1 Transfection and viral infection

For preparation of viral stocks, 2 million HEK293TN cells in 10-cm dishes were transfected with 3 µg pCMVΔR8.2, 2 µg pVSV-G, and 6 µg barcoded pHCC1 [23,57]. After 16 hr, the medium was replaced. The supernatant was collected 48 and 72 hr after transfection. Transfection efficiency was validated on the basis of the percentage of GFP(+) cells determined through FACS analysis. To determine the viral stock titer, five-fold serial dilutions of virus stocks were titrated in quadruplicate in 100 µL culture medium in 96-well culture plates. After 48 hr, the percentage of GFP(+) cells was measured through FACS analysis. 1 million Jurkat cells in 6-well plates were infected with barcoded viral inocula at an MOI of approximately 0.03. The medium was replaced with 3 mL fresh RPMI 24 hr after infection. 48 hr after infection, the efficiency of infection was monitored by FACS analysis.

### 2.2 Establishment of single provirus-integrated cellular clone (sinpro clone) models

Sinpro clones established from a pool of infected Jurkat cells with HIV-based vector, pHCC1 [23,57]. Infected cells were FACS-sorted using BD FACSAria™ (BD Biosciences) four days post-infection. The population of GFP(+) cells was diluted down to one cell per well in 96-well plates to obtain clones (**Fig. 1A** and **2A**). After a period of about two weeks of growth, we screened the plate for GFP expression from viable cells (n = 97) by measuring the percentage of GFP(+) cells using BD FACSCanto™ II (BD Biosciences). Starting from 10 96-well plates, eventually, 33 clones with expression of GFP were obtained. To further characterize the transcriptional phenotype of each clone, parameters, including the percentage of GFP(+) cells [(%) GFP(+)], GFP expression (GFP-fluo-gated and GFP-fluo-total), standard deviation (SD), the coefficient of variation (CV-gated and CV-total) (**Fig. 1B**) were recorded using BD FACSCanto™ II (BD Biosciences) twice every week for five weeks (**Fig. S1A**, a total of 11 subsequent time points). GFP expression and the coefficient of variation were independently measured from either GFP(+)-gated cells or a population of total live cells.

### 2.3 Cell culture

The human Jurkat T cell line (obtained from the cell collection of the Center for Genomic Regulation, Barcelona) was grown at 37 °C under a 95% air and 5% CO2 atmosphere, in RPMI 1640 medium (Gibco, Cat. No. 42401-018) supplemented with 10% fetal bovine serum (FBS; Gibco, Cat. No. 10270-106), 1% penicillin-streptomycin (Gibco, Cat. No. 10378016), and 1% GlutaMAX (100x) (Gibco, Cat. No. 35050-061). Jurkat cells were passaged every 2 d with a 1:5 dilution. HEK293TN cells were grown under the same conditions in Dulbecco’s modified Eagle’s medium (DMEM; Gibco, Cat. No. 41965-039). Sinpro clones were cultured in the same conditions as the human Jurkat T cell line. Cells were tested for mycoplasma yearly.

### 2.4 Flow cytometry measurements

Flow cytometry experiments were conducted using the BD FACSCanto™ II (BD Biosciences) machine and standard protocols. FACS-sorting of GFP_Bright_ and GFP_Dim_ subpopulations of clone #814 was conducted using the BD FACSAria™ (BD Biosciences) machine. Single cells were filtered based on FSC-H and FSC-W filters. Live cells were filtered using a fluorescent intercalator 7-aminoactinomycin D (7-AAD; BioLegend, Cat. No. 420404) for staining the dead cells.

### 2.5 RNA extraction and DNase I treatment

Total RNA was extracted using the AllPrep DNA/RNA Mini Kit (50) (Qiagen, Cat. No. 80204) following the manufacturer’s instructions. Subsequently, total RNA was subjected to DNase I treatment (New England Biolabs, M0303S) according to the producer’s recommendations.

### 2.6 Strand-specific reverse transcription (RT)

Total RNA concentration after DNaseI treatment was measured using a Qubit 4 Fluorometer with QubitTM RNA High Sensitivity (HS) kit (Thermo Fisher, Cat. No. Q32852). Subsequently, RNA samples were reverse transcribed using an NG dART RT-PCR kit (EURx, Cat. No. e0802-02) according to the producer’s instructions. RT reactions were performed using 100 ng RNA with respective RT primers: sRNAs, primer HCP#266; asRNAs, primer HCP#264; RPL27A (internal control gene), primer HCP#270. Primers HCP#264 and HCP#270 were added to the dPCR mixture. The RT reaction was incubated at 50 °C for 1 hr and inactivated at 85 °C for 5 min. Primer sequences are available at **Table S5**.

### 2.7 Quantitative polymerase chain reaction reaction (qPCR)

qPCR was conducted using a CFX ConnectTM (Bio-Rad) instrument and PowerTrackTM SYBR Green Master Mix (Thermo Fisher, Cat. No. A46109) according to the producer’s instructions. 2 µl cDNA was added in each reaction containing respective primers in the final concentration of 0.3 µM (sRNAs, HCP#265 and HCP#266; asRNAs, HCP#263 and HCP#264; RPL27A, HCP#269 and HCP#270). The reaction mix was predenaturated at 95°C for 10 min and ran for 30 cycles under following conditions: 95 °C for 10 min, 58 °C for 30 s, and 72 °C for 30 s, finished by melting curve analysis. Primer sequences are available at **Table S5**.

### 2.8 Digital PCR (dPCR)

The dPCR procedure was performed using QIAcuity Digital PCR System (Qiagen, Cat. No. 911020 QIAcuity One, 5plex instrument), QIAcuity 26-k, 24-well Nanoplates (Qiagen, Cat. No. 250001) and Probe PCR Kit (Qiagen, Cat. No. 250102) according to the manufacturer’s recommendations. In brief, primer-probe mixes targeting both sense [HCP#265, HCP#266 and HCP#272 (5’-end fluorescence: FAM; 3’-end quencher: BHQ1)] and antisense [HCP#263, HCP#264 and HCP#271 (5’-end fluorescence: HEX; 3’-end quencher: BHQ1)] transcripts were added in a total 10 µl mixture containing non-diluted cDNA. The reaction mix was predenaturated at 95 °C for 10 min and ran for 30 cycles under presented conditions: 95 °C for 10 min, 58 °C for 30 s, and 72 °C for 30 s. Primer and probe sequences are available at **Table S5**.

### 2.9 Integration Site mapping

Schematic representation of the experimental workflow is shown in **Fig. S2M**. Genomic DNA extraction was carried out using the AllPrep DNA/RNA Mini Kit (50) (Qiagen, Cat. No. 80204) following the manufacturer’s instructions and digested using 2 µL 5,000 U/mL HpyCH4III (New England Biolabs, R0618S) and 2 µL 10 U/µL BplI (Thermo Scientific, Cat. No. ER1011) in Tango buffer complemented with 1× SAM in a 50 µL final volume at 37 °C for 3 hr. The digestion was terminated by incubating samples for 20 min at 65 ℃. Digested DNA was used for *in vitro* transcription (IVT) using HiScribe® T7 High Yield RNA Synthesis Kit (NEB, Cat. No. E2040S). The reaction was performed according to the producer’s recommendations with the addition of DTT at a final concentration of 5 mM and incubated at 37 ℃ for 16 hr. IVT products were digested with RNase-free DNase I (NEB, Cat. No. M0303S). DNase I-treated IVT products were diluted at room temperature in a final volume of 500 µl RNase-free water, containing 1 µl 20 mg/mL glycogen (Thermo Scientific™, Cat. No. R0561), and 50 µl 3M sodium acetate (Sigma, Cat. No. 71196-100ml), followed by adding 500 µl isopropanol (CHEMPUR, Cat. No. 117515002) to the mixture. The reaction was incubated at room temperature for 20 min. RNA was precipitated by centrifugation at 12,000 x g for 15 min, followed by two times washing using 70% ice-cold ethanol.

Purified IVT products were first verified using RT-PCR with primers HCP#237* and HCP#238* (**Table S5**) (**Fig. S2N**). Subsequently, poly(A) tailing was subjected to purified IVT products using *E. coli* Poly(A) Polymerase (NEB, Cat. No. M0276) at 37 ℃ for 30 min, followed by reverse transcription (NG dART RT-PCR kit, EURx, Cat. No. E0802-02) using a string of polythymine in the length of 33 bp, followed by two additional random nucleotides with options guanine, adenine, or cytosine at the 3’ end of the primer sequence (HCP#273, **Table S5**) at 48 ℃ for 1 hr. The obtained cDNA was used in two independent PCR reactions: one aims to identify the viral integration site (primers HCP#279 and HCP#290, **Table S5**), and another targets the HIV 5’LTR (HCP#237* and HCP#238*, **Table S5**), serves as a PCR positive control. Both reactions were performed using Phusion™ High-Fidelity DNA Polymerases (2 U/µL) (Thermo Scientific, Cat. No. F534S) according to the protocol recommended by the producer. 5 µl non-diluted cDNA was used for the reaction in a total 20 µl reaction mixture. The PCRs ran in the following cycling conditions: 30 cycles, initial denaturation at 98 ℃ for 10 min, denaturation at 98 ℃ for 10 s, annealing at 59 ℃ for 30 s, extension at 72 ℃ for 30 s, final extension at 72 ℃ for 5 min. The PCR products from the first round of PCR amplification were purified using MinElute PCR Purification Kit (Qiagen, Cat. No. 28004), followed by the second round of PCR with tailored indexing primers (HCP#304 and HCP#305, **Table S5**). The same reaction condition and cycling parameters as the first round of PCR amplification were used. The final amplification products were purified, once again, using MinElute PCR Purification Kit (Qiagen, Cat. No. 28004) and visualized on 1.5% (wt/vol) agarose gel (**Fig. S2O**). The identified integration site of the provirus in clone #814 was verified using PCR amplification with two pairs of the primers, HCP#364 and HCP#238*, as well as HCP#365 and HCP#238*, which yield a specific 560 bp and 650 bp PCR product, respectively (**Fig. 2L**). Primer sequences are available at **Table S5**.

### 2.10 Illumina high-throughput sequencing of HIV integration sites (IS)

Integration site mapping amplicons were pooled at a final concentration of 2 nM for high-throughput sequencing. Libraries were sequenced as 50 bp single reads on a NovaSeq 6000 sequencer (Illumina) with the NovaSeq 6000 SP Reagent Kit v1.5 (100 cycles) (Illumina, Cat. No. 20028401).

### 2.11 Drug reactivation assay

1 million established sinpro clonal cells (post screening) were seeded per well in 6-well plates for the performance of drug reactivation. 81 nM phorbol 12-myristate 13-acetate (PMA) (Sigma-Aldrich, Cat. No. P8139) with 1.134 µM ionomycin (ThermoFisher, Cat. No. I24222) or 200 nM ingenol 3-angelate (PEP005) (Sigma-Aldrich, Cat. No. SML1318) were applied to provirus reactivation. Dimethyl sulfoxide (DMSO) (NEB, Cat. No. B0515A) was used as a negative control at a final concentration of 0.1% (v/v). After 24 hr incubation, cells were collected for cytometric measurements. Total RNA was isolated from cells treated with PMA plus ionomycin, PEP005, and DMSO, respectively, and used for RT-qPCR analysis.

## 3 Theory/calculation

### 3.1 The framework of noise space and computation of its components

The concept of noise space was borrowed from the study of Dar et al. [10]. It consists of three parameters: coefficient of variation (CV), GFP expression and the autocorrelation time (τ_1/2_). In this work, we divided parameters related to noise space into four sets: (1) population-based (pb)-CV-total, GFP-fluo-total, and τ_1/2_-noise-total, (2) pb-CV-gated, GFP-fluo-gated, and τ_1/2_-noise-gated, (3) time-based (tb)-CV-total, GFP-fluo-total, and τ_1/2_-fluo-total, and (4) tb-CV-gated, GFP-fluo-gated, and τ_1/2_-fluo-gated (**Fig. 1B**). To summarize, noise space was visualized in GFP expression measured in either population of total live cells or GFP(+)- gated cells; each scenario was further coupled with either pb-CV and τ_1/2_-noise (**Fig. 1D** and **1E**; **Fig. 2D** and **2E**) or tb-CV and τ_1/2_-fluo (**Fig. 1F** and **1G**; **Fig. 2F** and **2G**). Code used for the visualization of noise space is described in the following section. Computation of the parameters CV and τ_1/2_ is described below.

#### 3.1.1 Coefficient of variation (CV)

CV values for both gated and all live cells for each clone and each measurement time point were recorded directly from the flow cytometry results. The values were converted from percentage to ratio and their respective means were taken as pb-CV-gated and pb-CV-total. tb-CV-gated and tb-CV-total were measured based on the formula *cv_tb_* = *δ_tb_* /*m_tb_*; that is, the standard deviation *δ_tb_* of GFP fluorescence normalized by the GFP fluorescence mean *m_tb_*.

#### 3.1.2 Noise autocorrelation time (τ_1/2_)

In this study, the calculation of τ_1/2_ was computed based on mean transcription levels (namely τ_1/2_-fluo-total and τ_1/2_-fluo-gated) or mean-centered transcriptional noise (namely τ_1/2_-noise-total and τ_1/2_-noise-gated). First, data for GFP-fluo-gated, GFP-fluo-total, CV-gated, and CV-total were linearly interpolated to a regular spacing of one day. To calculate τ_1/2_-fluo-total and τ_1/2_-fluo-gated, autocorrelation function available in Python 3.11 statsmodels 0.14.4 (https://www.statsmodels.org/stable/index.html), the function acf(), was immediately applied for all possible time lags in days to the regularized GFP-fluo values. τ_1/2_-fluo-total and τ_1/2_-fluo-gated were recorded as the earliest time lags at which the autocorrelation was lower than 0.5.

The calculation of τ_1/2_-noise-total and τ_1/2_-noise-gated followed the concept described in Austin et al. [33], where τ_1/2_ was calculated based on transcriptional noise of individual clones against the noise averaged from clones in the same group [10,33]. Given such a limitation of our experimental instrument with which we cannot record CV for individual cells in a population of cells, using the same concept proposed by Austin et al. [33], we reformulated the calculation: regularized CV values were first mean-centered, followed by the application of autocorrelation functions available in Python 3.11 statsmodels 0.14.4 (<u>https://</u> www.statsmodels.org/stable/index.html), the function acf(). The first time lag, at which the autocorrelation function value dropped below 0.5 was recorded as τ_1/2_. The analytical pipeline is available at GitHub (https://github.com/HCAngelC/HIV_sinpro_clones_project).

#### 3.1.3 Computing the noise frequency range (F_N_)

The calculation of F_N_ was based on the definition *F_N_* = 1/*τ*_1/2_, published by Austin et al. [33]. In this study, we took the values of τ_1/2_-fluo-total, τ_1/2_-fluo-gated, τ_1/2_-noise-total, τ_1/2_-noise-gated to compute F_N_, respectively (**Figure S1O**).

### 3.2 Data analysis

#### 3.2.1 Plotting 2D and 3D projection of the parameters under the framework of noise space

##### 3.2.1.1 2D projection

The correlation between every pair of three parameters, including CV, GFP expression and τ_1/2_, required for the construction of noise space was plotted using the R package “ggplot” [58] (**Fig. S1I**-**S1N**; **Fig. S2F** and **S2G**). Four different sets of the parameters, including (1) pb-CV-total, GFP-fluo-total, and τ_1/2_-noise-total, (2) pb-CV-gated, GFP-fluo-gated, and τ_1/2_-noise-gated, (3) tb-CV-total, GFP-fluo-total, and τ_1/2_-fluo-total, and (4) tb-CV-gated, GFP-fluo-gated, and τ_1/2_-fluo-gated, were plotted independently.

##### 3.2.1.2 3D projection

In noise space, three coordinate axes were given as follows: (1) x*-*axis was referred to as GFP expression in a logarithmic scale (log_2_), (2) y*-*axis was referred to as τ_1/2_, and (3) z-axis was referred to as the square of CV in a logarithmic scale (log_2_) (**Fig. 1D-1G** and **Fig. 2D-2G**). Noise space was visualized using the function scatter3D() implemented in the R package “plot3D” (version 1.4.1) (https://cran.r-project.org/web/packages/plot3D/index.html), separated by the four sets of the parameters (**Fig. 1D-1G**; **Fig. 2D-2G**).

#### 3.2.2 The performance of principal component assay (PCA) on parameters measured by flow cytometry across sinpro clones

Four parameters, including CV, SD, the mean GFP expression and τ_1/2_ were included for the performance of PCA (**Fig. S1E**-**S1H**; **Fig. S2H**-**S2K**) using the function PCA() imbedded in the R package “FactoMineR” (version 2.11) [59] and visualized using the function fviz_pca_biplot() embedded in the R package “factoextra” (version 1.0.7) [60]. PCA was independently performed, aligning with the criteria assigning four different sets of parameters in this study.

#### 3.2.3 The network-based analysis on parameters measured by flow cytometry across sinpro clones

Five parameters, including CV, SD, the mean GFP expression, τ_1/2_, and F_N_ were utilized for the calculation of correlation coefficients between two adjacent vertices in a network property using the function rcorr() embedded in the R package “Hmisc” (Version 5.2-2) (https://CRAN.R-project.org/package=Hmis). Subsequently, correlation matrices were transformed into data frames containing four columns. The first two columns served as ID columns, designating two adjacent enriched signatures irrespective of the direction. The remaining columns included the correlation coefficient and the associated *p*-value. To filter out weak or spurious connections, correlation coefficients with a *p*-value > 0.05 are excluded, thereby generating a correlation matrix. We further utilized this correlation matrix as the edge list to depict simple graphs (**Fig. 1H-1K**; **Fig. 2H-2K**; **Fig. 4D**). In this list, the correlation coefficients representing the edge were calculated between two adjacent vertices that represent individual sinpro clones. The node list consisted of a series ID for individual sinpro clones, information about groups (**Fig. 1H-1K**; **Fig. 2H-2K**) or models (**Fig. 4D**) assigned to respective clones. Once again, we structured individual network properties for four different sets of parameters, respectively. The network structure was then established using the function graph_from_data_frame() from the R package ‘‘igraph’’ (Version 2.1.4) (https://igraph.org) [61] with the following arguments: d for the edge list, vertices for the node list, and directed = FALSE to account for undirected edges in the network.

#### 3.2.4 Mapping HIV integration site (IS) in sinpro clones

The code for mapping HIV integration sites was written in Python 3.12 using pandas 2.2.3 [62], Biopython 1.84 [63] and pysam 0.22.1 [64] packages. The source code is available at https://github.com/wis-janusz/LISseq, while the latest release (Version 1.2.2) is available at https://github.com/wis-janusz/LISseq/releases.

50 bp single-read raw reads were filtered as follows: first, the 5’ LTR sequence GGAGTGAATTAGCCCTTCCA was removed from the 5’ end of reads. Any reads that do not contain the 5’LTR sequence with 1 mismatch allowed were discarded. Next, the poly(A) sequence in length of minimal 5 bp was removed from the 3’ ends of the reads. Finally, reads that contain at least 1 ambiguous base or position with the phred quality score (q-score) lower than 20 were discarded.

The filtered reads were mapped to the human reference genome GRCh38 no-alt loci analysis set (https://ftp.ncbi.nlm.nih.gov/genomes/all/GCA/000/001/405/GCA_000001405.15_GRCh38/seqs_for_alignment_pipelines.ucsc_ids/GCA_000001405.15_GRCh38_no_alt_analysis_set.fna.bowtie_index.tar.gz) using bowtie2 version 2.5.4 [65] with default parameters. Alignments with quality lower than 20, as reported by bowtie2, were discarded. Filtered alignments were grouped by chromosome number and position. Loci with sequencing depth, i.e., the number of reads aligned to the same chromosome and position, of less than 1000 were discarded. The reported integration site indicated the single locus with the highest sequencing depth in this study.

#### 3.2.5 Visualization of the landscape of (epi)genomic features surrounding the provirus integration site in sinpro clones

The landscape of repeating elements in the genomic region of 50,000 base pairs upstream and downstream of the provirus integration site (chr13:110,913,078) in clone #814 was plotted using R with default functions (**Fig. 2M**). Smith Waterman (SW) alignment score was applied to represent the peak intensity. Human, Homo sapiens, (hg38) DNA sequences for interspersed repeats and low complexity DNA sequences data were downloaded from RepeatMasker (https://www.repeatmasker.org/species/hg.html).

ChIP-seq reads, including signals from H3K27ac, H3K4me1, H3K4me3, H3K36me3, H3K9me3 and H3K27me3 were downloaded from the ENCODE Project Consortium [66] (https://ftp.ncbi.nlm.nih.gov/genomes/all/GCA/000/001/405/GCA_000001405.15_GRCh38/seqs_for_alignment_pipelines.ucsc_ids/GCA_000001405.15_GRCh38_no_alt_analysis_set.fna.bowtie_index.tar.gz) and plotted in the genomic region of 500,000 base pairs upstream and downstream of the provirus integration site (chr13:110,913,078) in clone #814 using R with default functions (**Fig. 2N**). ChIP-seq signal value (fold change over control) was applied to represent the intensity of signal peaks. The landscape of chosen histone modifications is also visualized using the Integrative Genomics Viewer (IGV) (IGV version 2.19.1 for Linux) [67] (**Fig. S2P**). The characteristics of the provirus integration sites retrieved in sinpro clones #760 (the A- group), #761 (the A- group), #762 (the A group), #770 (the A group), #776 (the A- group), #778 (the S group), #788 (the S group), #800 (the S group), #802 (the A- group), #803 (the A- group), #811 (the A group), #812 (the A group), and #813 (the A- group) using lentiviral integration site sequencing (LIS-seq) is summarized in **Fig. S3A**. Distances (in logarithmic scale of base pairs) to the closest histone marks of the provirus integration sites mentioned above were computed and plotted in **Fig. S3B**-**S3G**.

#### 3.2.6 Obtaining the best-fitted curve of exponential decay model

##### 3.2.6.1 Reciprocal curve model

The formula embedded in the reciprocal curve model was taken from the equation previously reported by Singh et al. [9] to systematically quantify variability in GFP expression driven by the HIV 5’LTR across different HIV integration sites in the human genome. The equation is written as follows:

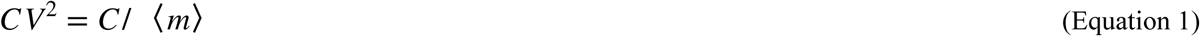

where *C* is the proportionality factor and 〈*m*〉 is the average number of GFP molecules per cell (〈 〉 denotes the average) [9,68]. The performance of curve fitting on 32 sinpro clones in A, A-, and S groups was conducted using tailored functions implemented in Python 3.11 SciPy 1.15.1 [69] (available at GitHub https://github.com/HCAngelC/HIV_sinpro_clones_project). The curve was visualized in **Fig. 4B**.

##### 3.2.6.2 Exponential decay model

The performance of curve-fitting of exponential decay was based on the equation written as follows:

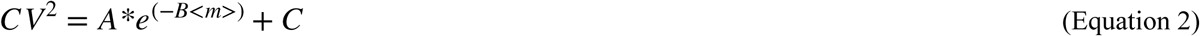

where *CV* is the coefficient of variation, *m* is the mean GFP fluorescence, *e* is the Euler’s number and *A*, *B*, and *C* are fitted constants (**Fig. 4F**). Five distinct models of the exponential decay model across 33 sinpro clones were determined based on Equation 2 using tailored functions implemented in Python 3.11 SciPy 1.15.1 [69] (available at GitHub https://github.com/HCAngelC/HIV_sinpro_clones_project). The curve was visualized in **Fig. 4C**. Values of the constants A, B, C in each model were shown in **Fig. 4F**.

To further probe the influence of sense- and antisense RNAs on stochastic variation driven by the HIV 5’LTR and the 3’LTR, independently at the transcription level, the parameter value of 〈*m*〉 was replaced by the values measured from either sRNAs (**Fig. 4H** and **4J**) or asRNAs (**Fig. 4I** and **4K**) using strand-specific qPCR and dPCR. To determine a set of initial constants A, B, and C to fit the model, the constant C was estimated as its value has to be smaller than the minimum value of *CV*^2^and be greater than zero. The constants A and B were determined based on a linear model transformed from Equation 2. The mathematical operation was demonstrated as follows:

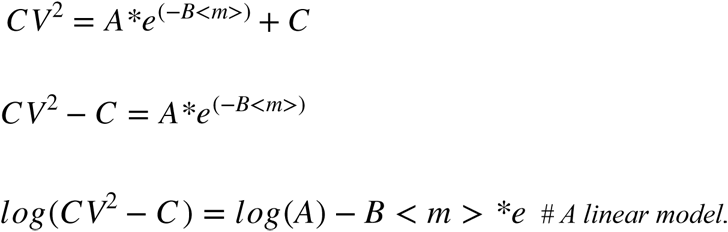

Step by step operation code is available at Git Hub (https://github.com/HCAngelC/HIV_sinpro_clones_project). In brief, models were tested using the function nls() implemented in the R package “stats” (version 4.3.3) (https://stat.ethz.ch/R-manual/R-devel/library/stats/html/stats-package.html). Estimated values of each constant in models were recorded using the function summary() implemented in the R. Of note, in the circumstance that the default Gauss-Newton algorithm in the function nls() cannot support the estimated values for the initiation of curve fitting, a feasible alternative of Golub-Pereyra algorithm for Partial Least Squares model with the argument alg = "plinear" built-in in the function nls() was applied. The plot of fitted curves (**Fig. 4**) were visualized using R with default functions.

### 3.3 Data and code availability

#### 3.3.1 Data availability

The raw sequencing data generated during this study are available from Sequence Read Archive (SRA) (PRJNA1244909).

FACS measurements (measured in 2023) on 33 sinpro clones at 11 subsequent time points (**Fig. 1A**, **2A,** and **2B**; **Fig. S1A**; **Fig. 4B** and **4C**) are available at **Supplementary Information**, **Supplementary Table S1**.

Measures of four sets of parameters used to construct noise space (**Fig. 1D-1G**; **Fig. 2D-2G**) are available at **Supplementary Information, Supplementary Table S2.**

FACS measurements (measured in 2024) on FACS-sorted clone #814 GFP_Bright_ and GFP_Dim_ subpopulations at six subsequent time points (**Fig. 2C**) are available at **Supplementary Information**, **Supplementary Table S3**.

FACS measurements (measured in 2024) coupled with sRNAs and asRNAs measured by strand-specific qPCR and dPCR from nine sinpro clones at six subsequent time points (**Fig. 3**; **Fig. 4G-4I**; **Fig. S5C**; **Fig. S6**; **Fig. S7**) are available at **Supplementary Information**, **Supplementary Table S4**.

**Fig. 3.**
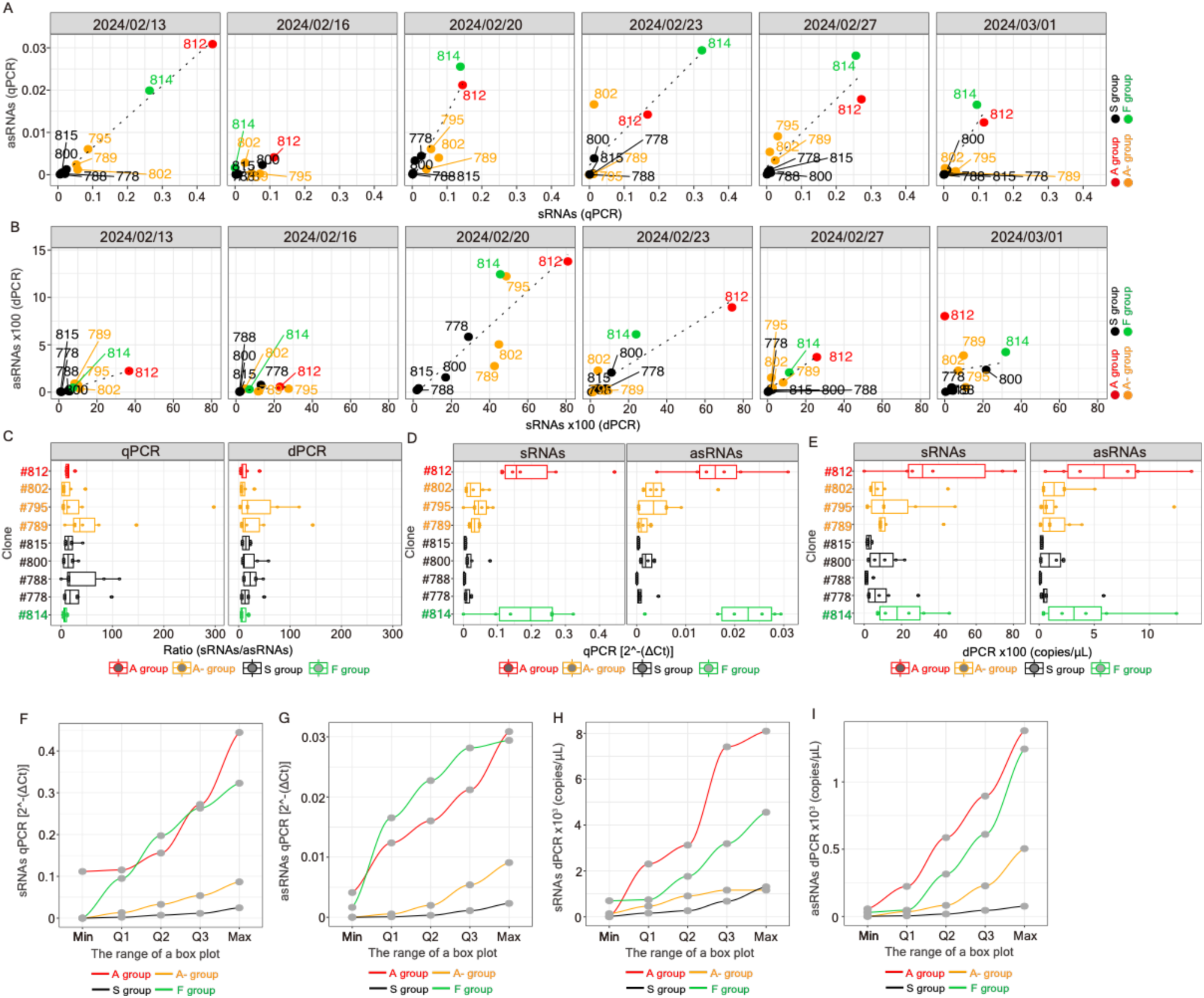
Similar trends of consecutive expression between sRNAs and asRNAs. (**A**, **B**) Scatter plots representing the correlation between sense and antisense transcripts measured via strand-specific RT, followed by qPCR (**A**) or dPCR (**B**) at six subsequent time points. Facets separate the measurements at different time points. Spots marked in red, orange, black, and green represent sinpro clones in the A (#812), A- (#789, #795, and #802), S (#778, #788, #800, #815), and F (#814) groups. (**C**) Boxplot representing the ratio between sense and antisense transcripts (sRNAs versus asRNAs) across nine sinpro clones. Facets separate the performance of measurements using qPCR or dPCR. Boxes marked in red, orange, black, and green represent sinpro clones in the A (#812), A- (#789, #795, and #802), S (#778, #788, #800, #815), and F (#814) groups. (**D**, **E**) Boxplots representing the abundance of sense and antisense transcripts measured via qPCR (**D**) or dPCR (**E**) across nine sinpro clones. Facets separate the measures of sRNAs versus asRNAs. Boxes marked in red, orange, black, and green represent sinpro clones in the A (#812), A- (#789, #795, and #802), S (#778, #788, #800, #815), and F (#814) groups. (**F**, **G**, **H**, **I**) Line plots representing the transcription threshold between sense and antisense transcripts measured via qPCR (**F**, sRNAs; **G**, asRNAs) or dPCR (**H**, sRNAs; **I**, asRNAs) across nine sinpro clones. The abundance of transcripts from clones in each group is compared in the range from the minimum (Min.), lower quartile (Q1), median (Q2), upper quartile (Q3), to the maximum (Max.) values (x-axis). Lines in red, orange, black, and green represent sinpro clones in the A (#812), A- (#789, #795, and #802), S (#778, #788, #800, #815), and F (#814) groups, respectively. See also **Fig. S4**-**S6** and **Table S4**.

Sequences of primers and probes are available at **Supplementary Information**, **Supplementary Table S5**.

Human, Homo sapiens, (hg38) DNA sequences for interspersed repeats and low complexity DNA sequences data were downloaded from RepeatMasker (https://www.repeatmasker.org/species/hg.html).

ChIP-seq data from Jurkat clone E6-1 and CD4+, αβ T cells for H3K27ac (ENCFF338LAI, ENCFF701MUZ), H3K4me1 (ENCFF002WRW, ENCFF704DNO, ENCFF068SDD, ENCFF545FKD), H3K4me3 (ENCFF860LFO, ENCFF017AFO, ENCFF923RAA, ENCFF067KWB, ENCFF280EJY, ENCFF855AXA), H3K36me3 (ENCFF536HDR, ENCFF275ZSW, ENCFF215CDE), H3K9me3 (ENCFF010VZL, ENCFF154QXP, ENCFF353AHL, ENCFF201NFN) and H3K27me3 (ENCFF720HUE, ENCFF514ZIY, ENCFF153LTQ, ENCFF669USD) were downloaded from the ENCODE Project Consortium^84^ (https://ftp.ncbi.nlm.nih.gov/genomes/all/GCA/000/001/405/GCA_000001405.15_GRCh38/seqs_for_alignment_pipelines.ucsc_ids/GCA_000001405.15_GRCh38_no_alt_analysis_set.fna.bowtie_index.tar.gz). Experiments in ENCFF 338 LAI, ENCFF701MUZ, ENCFF002WRW, ENCFF068SDD, ENCFF 280 EJY, ENCFF855AXA, ENCFF215CDE, ENCFF010VZL, ENCFF154QXP, ENCFF720HUE, and ENCFF514ZIY were conducted using Homo sapiens CD4+, αβ T cells treated with phorbol 13-acetate 12-myristate and ionomycin; experiments in ENCFF704DNO, ENCFF545FKD, ENCFF923RAA, ENCFF067KWB, ENCFF536HDR, ENCFF275ZSW, ENCFF353AHL, ENCFF201NFN, ENCFF153LTQ, and ENCFF669USD were conducted using Homo sapiens CD4+, αβ T cells without stimulation; experiments in ENCFF860LFO, and ENCFF017AFO were conducted using Jurkat clone E6-1.

#### 3.3.2 Code availability

Analytical pipelines and scripts generated in this work are available at GitHub (https://github.com/HCAngelC/HIV_sinpro_clones_project). The script for mapping provirus integration sites is available at GitHub (https://github.com/wis-janusz/LISseq). Any additional information required to reanalyze the data reported in this paper is available from the lead contact upon request. This paper reports the original code. The open-source packages used in this study, which have not been assigned DOIs, are listed as follows: (1) Python 3.11 statsmodels 0.14.4—Seabold, Skipper, and Josef Perktold. “statsmodels: Econometric and statistical modeling with python.” Proceedings of the 9th Python in Science Conference. 2010 (<u>https://</u> www.statsmodels.org/stable/index.html); (2) RepeatMasker—Smit, AFA, Hubley, R & Green, P. *RepeatMasker Open-4.0*. 2013-2015 (http://www.repeatmasker.org); (3) R package “Hmisc” (Version 5.2-2)—Harrell Jr., F., & Dupont, Ch. (2019). Hmisc: Harrell Miscellaneous (https://CRAN.R-project.org/package=Hmis); (4) R package “plot3D” (Version 1.4.1)—Soetaert, K. (2024). plot3D: Plotting Multi-Dimensional Data (https://cran.r-project.org/web/packages/plot3D/index.html); (5) R package “stats” (Version 4.3.3)—R Core Team (2024). _R: A Language and Environment for Statistical Computing_. R Foundation for Statistical Computing, Vienna, Austria (https://stat.ethz.ch/R-manual/R-devel/library/stats/html/stats-package.html).

### 3.4 Statistics

All statistical tests were performed using Python or R (version 4.3.3) with default options and specific details are provided in the main text and figure legends where applicable.

## 4 Results

### 4.1 Establishment of sinpro clones and parameter setting

The selection procedure for sinpro clones is described in detail in the **Material and methods**. After a screening of 10 96-well plates, 33 sinpro clones were obtained (**Fig. 1A** and **2A**). We performed a flow cytometry analysis to record and calculate a total of 16 parameters (**Fig. 1B**) at 11 subsequent time points (**Fig. S1A**). Descriptions of each parameter can be found in **Fig. 1B** and the **Material and methods**. Of note, given that the viability of each clone differs (some clones were robust in culture for a long period of time; some were not), not all sinpro clones could be included in every experiment planned in this work.

Based on the sequential measurement of the percentage of GFP-positive cells [(%) GFP(+)], these 33 clones can be categorized into four groups: clones in which proviruses remained transcriptionally active over 11 subsequent time points, were assigned to the active group (the A group, 10 clones); clones in which proviruses demonstrated a decrease in transcriptional activity over time, were assigned to the attenuated active group (the A- group, 17 clones); clones in which the transcription of proviruses showed a sharp decrease and remained transcriptionally silent, were assigned to the silent group (the S group, 5 clones) (**Fig. 1A**; **Fig. S1A** and **S1B**); and one clone (clone #814) in which the turnover of the state of HIV transcription showed elevated frequency was assignd to the F group (**Fig. 2A**). Given such a unique phenotype of provirus transcription in clone #814, more details will be discussed separately in the following content.

With respect to clones in the A, A-, and S groups, a tendency for a positive correlation was observed between (%) GFP(+) and the mean GFP fluorescence measured from GFP(+)-gated cells (GFP-fluo-gated; R^2^ = 0.193; cor = 0.939) (**Fig. S1C**) or from a population of total live cells (GFP-fluo-total; R^2^ = 0.724; cor = 0.85) (**Fig. S1D**). Of note, in addition to the mean coefficient of variation (CV) measured via flow cytometry at independent time points [namely population-based (pb)-CV], we calculated CV for a time series across 11 time points [namely time-based (tb)-CV; detailed in the **Material and methods]** for comparison. Overall, a tendency for a negative correlation was observed between pb-CV, separated by the measurement from GFP(+)-gated cells (pb-CV-gated) or a population of total live cells (pb-CV-total), and GFP-fluo-gated and GFP-fluo-total, respectively (**Fig. S1I**). A similar observation of a modestly negative correlation was observed when tb-CV was applied (**Fig. S1J**). This observation indicated that the transcription of proviruses in the A, A-, and S groups aligned with that in previous findings [9,58]: a gene that exhibits a higher level of transcription has a lower degree of CV.

To better characterize sinpro clones across different groups, we constructed noise space (**Fig. 1C-1G**) using FACS-relevant parameters, divided into four sets (**Fig. 1B**). The definition of noise space was borrowed from the study of Dar et al. [10] with modifications. In principle, noise space consists of three parameters: CV, the mean GFP expression, and the autocorrelation time (τ_1/2_). An analysis of the τ_1/2_ axis is critical given that it facilitates a direct comparison of data containing a wide variety of expression levels by removing the reciprocal dependence of noise magnitude on expression level [70]. In this work, τ_1/2_ was independently calculated based on mean transcription levels (namely τ_1/2_-fluo-total and τ_1/2_-fluo-gated) or mean-centered transcriptional noise (namely τ_1/2_-noise-total and τ_1/2_-noise-gated). The rationales and calculations are described in the **Material and methods**. Overall, four sets of parameters were applied to construct noise space: (1) pb-CV-total, GFP-fluo-total, and τ_1/2_-noise-total (**Fig. 1D**), (2) pb-CV-gated, GFP-fluo-gated, and τ_1/2_-noise-gated (**Fig. 1E**), (3) tb-CV-total, GFP-fluo-total, and τ_1/2_-fluo-total (**Fig. 1F**), and (4) tb-CV-gated, GFP-fluo-gated, and τ_1/2_-fluo-gated (**Fig. 1G**).

### 4.2 Visualization of transcriptional characteristics of sinpro clones in noise space

We first examined whether the mentioned four parameter sets were sufficient to classify sinpro clones across different (A, A-, and S) groups in two-dimensional (2D) projections of the principal component analysis (PCA) (**Fig. S1E**-**S1H**). Overall, only a modest classification of clones across different groups in different parameter sets was observed (**Fig. S1E**-**S1H**). A separation was demonstrated as the parameters related to the population of live cells were implemented (**Fig. S1E** and **S1G**) compared with PCA conducted using the parameters related to GFP(+)-gated cells (**Fig. S1F** and **S1H**). In each set, a positive correlation between the mean GFP expression (GFP–fluo-total and GFP-fluo-gated) and a standard deviation (SD) was shown; a lack of correlation between the mean GFP expression and τ_1/2_, as well as CV was present (**Fig. S1E**-**S1H**). Given that a linear correlation is expected between CV and the mean GFP expression as 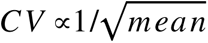, a lack of correlation between these parameters may suggest either an atypical phenotype of stochastic HIV gene expression [35] in sinpro clones or the need for additional parameters to characterize its fluctuations in transcription.

A clear separation of clones across the three groups (A, A-, and S) was shown in noise space when a population of total live cells was applied irrespective of τ_1/2_ (**Fig. 1D**, τ_1/2_-noise-total; **Fig. 1F**, τ_1/2_-fluo-total). The separation between clones among the A, A-, and S groups was less effective in noise space constructed based on the parameters related to the GFP(+)-gated cells (**Fig. 1E** and **1G**). Using network-based analysis, we verified that the parameters related to a population of total live cells (**Fig. 1H** and **1J**) enabled the structural topology of an assortative network (assortativity = 0.045, **Fig. 1H**; assortativity = 0.021, **Fig. 1J**), whereas networks structured based on the parameters related to the GFP(+)-gated cells (assortativity = −0.088, **Fig. 1I**; assortativity = −0.088, **Fig. 1K**) failed to separate clones in which the proviruses possessed different transcriptional phenotypes. The same observation was seen in the correlation between every pair of the three parameters used for defining noise space assayed in a 2D projection (**Fig. S1I**-**S1N**). We stress that transcriptional bursts varied across clones within the same group while comparing the noise frequency range (F_N_) [33] (**Fig. 1B**) calculated for the four parameter sets (**Fig. S1O**: F_N_-fluo-total, F_N_-fluo-gated, F_N_-noise-total, and F_N_-noise-gated). Altogether, these observations suggest that the transcriptional characteristics in sinpro clones could be better defined based on the parameters related to a population of total live cells, rather than a focus on GFP(+)-sorted cells only. In the following analysis, the focus will be on these parameter sets.

### 4.3 Characteristics of the unique model, clone #814, unveiling stochastic HIV transcription with elevated frequency

We obtained one clone, clone #814, in which the turnover of the state of HIV transcription demonstrated an elevated frequency (**Fig. 2B**), as visualized in the measurement of (%) GFP(+) cells over time (**Fig. 2B**). Such a stochastic phenotype of HIV transcription reappeared as GFP-fluo-gated and GFP-fluo-total were measured at 11 subsequent time points (**Fig. 2B**). We also observed an inconsistency between the measures of (%) GFP(+) and GFP-fluo-gated: even though a small subset of (%) GFP(+) cells was present in the clonal population of clone #814, the highest level of proviral transcription was still detected (**Fig. 2B**, highlighted by a red dotted line), whereas similar curves were observed between the measures of (%) GFP(+) and GFP-fluo-total (**Fig. 2B**), implying that, in clone #814, GFP fluorescence measured from a population og total live cells could better represent the heterogeneity of the clonal population shown by (%) GFP(+) cells. Remarkably, given that the site of HIV integration is identical regardless of the state of provirus transcription in clone #814, such a phenotype of stochastic HIV transcription should be a pure epigenetic phenomenon. The integration site of clone #814 will be discussed in the following content. Such stochastic fluctuations measured via (%) GFP(+) remained as additional independent measurements in a time series were conducted in 2025.

We wondered whether such a fluctuation in HIV transcription could be inheritable in both subpopulations of GFP_Bright_ and GFP_Dim_ of clone #814. To test this, we FACS-sorted a clonal population of clone #814 based on its GFP expression into two subpopulations, #814-GFP_Bright_ and #814-GFP_Dim_ (**Fig. S2A**), and, once again, recorded the FACS-related parameters (**Fig. 1B**) across six time points (**Fig. 2C**). Although similar dynamics between GFP-fluo-gated and GFP-fluo-total were observed (**Fig. 2C**), GFP-fluo-total showed a higher intensity in the #814-GFP_Bright_ subpopulation compared with that measured in the #814-GFP_Dim_ subpopulation (**Fig. S2C**), whereas GFP-fluo-gated was indistinguishable between these two subpopulations (**Fig. S2B**). In addition, the #814-GFP_Dim_ subpopulation demonstrated a significant increase in the measures of CV-gated and CV-total compared with those measured in the #814-GFP_Bright_ subpopulation (**Fig. 2C**; **Fig. S2D** and **S2E**). When examining the correlation between CV and GFP-fluo in both gated (**Fig. S2F**) and total live cells (**Fig. S2G**) at six subsequent time points, we observed that even though clonal cells in the #814-GFP_Dim_ subpopulation were more bursty than those in the #814-GFP_Bright_ subpopulation, all measures showed a negative regression, suggesting that the regulation of transcriptional bursting is likely inherited in both subpopulations. An undistinguishable F_N_ between these two subpopulations further verified this assumption (**Fig. S2L**).

When we assayed the spatial correlation between all sinpro clones, including #814-GFP_Bright_ and #814-GFP_Dim_ in noise spaces (**Fig. 2D-2G**), we observed that both subpopulations were in the spatial vicinity of each other, indicating a continuation of frequent turnover of transcriptional bursting in both subpopulations. The same observation was observed in the network properties structured based on parameters related to a population of total live cells: unsorted clone #814, #814-GFP_Bright_, and #814-GFP_Dim_ were in the vicinity of each other (**Fig. 2H** and **2J**). Once again, networks structured based on parameters related to the GFP(+)- gated cells failed to clearly discriminate between sinpro clones across different groups (**Fig. 2I** and **2K**). We noticed that after the addition of clone #814 and its subpopulations, clones from the A and A- groups were indistinguishable (**Fig. 2H** and **2J**) compared with those from our previous observation, shown in **Fig. 1H** and **1J**, suggesting the presence of additional subsets of clones between the A and A- groups. Additional mathematical approaches for the better discrimination of sinpro clones with additional subsets and clone #814 as well as its subpopulations based on provirus transcriptional bursting will be discussed in **Fig. 4**. While applying 2D projections of PCA, the distribution of #814-GFP_Bright_ and #814-GFP_Dim_ was also irrelevant to the transcriptional pattern of other sinpro clones (**Fig. S2H**-**S2K**). Altogether, these findings suggest that the mechanism that most likely governs the transcription of the provirus in clone #814 differed from that of other sinpro clones and that transcriptional bursting in clone #814 could be inheritable at a minimal level.

**Fig. 4.**
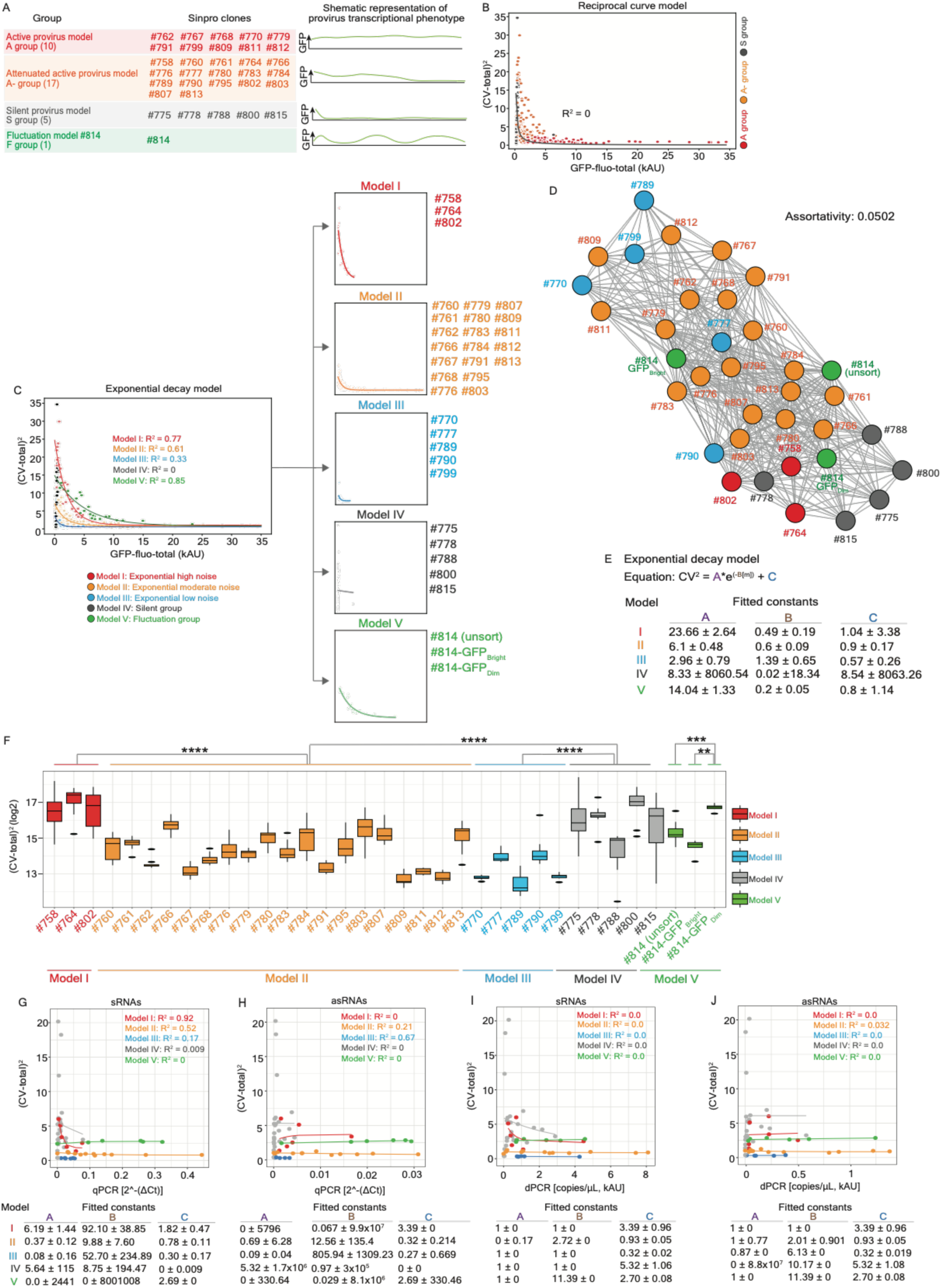
Curve fitting of exponential decay depicting the transcription phenotypes of sinpro clones. (**A**) Schematic summary of distinct transcriptional phenotypes across 33 sinpro clones assigned to four different groups based on the measurement of (%) GFP(+) cells. (**B**) Plot of the mean GFP expression (measured via GFP-fluo-total) versus transcription noise (measured via CV-total) across sinpro clones in the A, A-, and S groups. Solid black line represents the fitting curve (R^2^ = 0) based on the reciprocal curve model (Equation 1) mentioned in the main text. GFP-fluo-total is plotted with the values divided by 1,000; CV-total is plotted with the values divided by 100. (**C**) Plot of the mean GFP expression (measured via GFP-fluo-total) versus transcription noise (measured via CV-total) across sinpro clones in all groups. Solid lines represent fitting curves based on the exponential decay model (Equation 2) described in the main text, including (1) model I marked in red (the exponential high noise group, R^2^ = 0.77), (2) model II marked in orange (the exponential moderate noise group, R^2^ = 0.61), (3) model III marked in azure (the exponential low noise group, R^2^ = 0.33), (4) model IV marked in grey (the silent group, R^2^ = 0), and (5) model V marked in green (the fluctuation group, R^2^ = 0.85). Facets separate individual curves fitting in each model; sinpro clones assigned to respective groups are labeled on the right-hand side of facets. Of note, measures from #814-GFP_Bright_ and #814-GFP_Dim_ subpopulations are included in the analysis and assigned to the same model (model V) as unsorted clone #814. GFP-fluo-total is plotted with the values divided by 1,000; CV-total is plotted with the values divided by 100. (**D**) Graph networks (assortativity coefficient = 0.0502) illustrating the topological distribution of sinpro clones across five models. Spots in red, orange, azure, grey, and green represent clones in models I, II, III, IV, and V, respectively. (**E**) The formula of Equation 2 used for the simulation of curve fitting. The A, B, and C constants corresponding to individual models are summarized in the table beneath the formula. (**F**) Boxplot representing transcription noise (measured via CV-total) across 33 sinpro clones and FACS-sorted #814 GFP_Bright_ and GFP_Dim_ subpopulations. Clones assigned to models I, II, III, IV, and V are written in red, orange, azure, grey, and green, respectively. The y-axis is in logarithmic scale. \*\**P* < 0.01, \*\*\**P* < 0.001, \*\*\*\**P* < 0.0001. (**G**, **H**, **I**, **J**) Plot of the abundance of RNA transcripts, separated between sRNAs and asRNA versus transcription noise (measured via CV-total) across nine sinpro clones used in Fig. 3. The abundance of sense and antisense transcripts are measured using strand-specific RT, followed by either qPCR (**G**, sRNAs; **H**, asRNAs) or dPCR (**I**, sRNAs; **J**, asRNAs). Solid lines represent fitting curves based on the exponential decay model (Equation 2) described in the main text. Solid lines and spots corresponding to models I, II, III, IV, and V are marked in red, orange, azure, grey, and green, respectively. The A, B, and C constants corresponding to individual models are summarized in the table beneath each plot. See also **Fig. S7** and **Table S1**.

### 4.4 The provirus in clone #814 integrated in repetitive regions associated with active histone modifications

Given that we hypothesize that the fluctuation in HIV transcription in clone #814 could be a pure epigenetic phenomenon, we wondered whether any (epi)genomic features were involved. To identify the provirus integration site in clone #814, we developed a high-throughput method based on the T7 promoter-driven *in vitro* transcription, namely lentiviral integration site sequencing (LIS-seq) (**Fig. S2M** and **S2N**; detailed in the **Material and methods**). The specific PCR product was obtained before Illumina high-throughput sequencing (**Fig. S2O**). We obtained 403,596 raw reads after sequencing, and 252,571 reads remained after quality control and read filtering using our tailored pipeline (see the **Material and methods**), of which a total of 221,470 were mapped to the locus at 110,913,078 base pairs (bp) within the region of the protein-coding gene ankyrin repeat domain 10 (*ankrd10*) at the chromosome 13 (chr13) (**Fig. S2P**). The identified integration was confirmed via PCR amplification using a pair of primers: one annealed to the local genomic content surrounding the integration site and another annealed to the start of the HIV 5’LTR (**Fig. 2L**). We first assayed genomic features within the 10 kilobase pair (kb) region upstream and downstream from the site (chr13:110,913,078) of provirus integration. The first peak of repetitive sequences classified in the family short interspersed nuclear elements (SINEs) and Alu elements was present ∼1.6 kb upstream from the integration site (**Fig. 2M**). Furthermore, we observed intense ChIP-seq signals (fold change over control) of the acetylation of lysine 27 on histone H3 (H3K27ac) and trimethylation of lysine 4 on histone H3 (H3K4me3) within this 10 kb region—the first noticeable peak of H3K27ac was present at the position 661 bp upstream from the integration site, and the first peak of H3K4me3 was present at the position 108 bp upstream from the integration site (**Fig. 2N**). Other ChIP-seq signals, including monomethylation of lysine 4 on histone H3 (H3K4me1), trimethylation of lysine 36 on histone H3 (H3K36me3), trimethylation of lysine 9 on histone H3 (H3K9me3), and trimethylation of lysine 27 on histone H3 (H3K27me3), appeared at a basal level (**Fig. 2N**). The landscape of assayed ChIP-seq signals in the 20 kb genomic region surrounding the integration site is visualized in **Fig. S2P**. For comparison reasons, we further identified provirus integration sites in additional 13 sinpro clones assigned to the A (#762, #770, #811, and #812), A- (#760, #761, #776, #802, #803, and #813), and S groups (#778, #788, and #800) (**Fig. S3A**) and computed the distance to the closest histone marks of the provirus integration sites (**Fig. S3B**-**S3G**). We observed a tendency that proviruses in clones assigned to the A and A- groups are in the vicinity to active histone marks (**Fig. S3B**-**S3D**) except H3K36me3 (**Fig. S3E**), whereas proviruses in clones assigned to the S group are in the vicinity to repressive histone marks (**Fig. S3F** and **S3G**). This observation aligned with our previous findings [23,25,26], indicating the feasibility of using sinpro clones and LIS-seq to dissect the interplay between stochastic HIV transcription and the local genomic context associated with epigenetic features. A lack of statistical significance that appears for certain histone marks might be due to the small number of clones that were examined here. The provirus in clone #814 preferentially integrated closer to H3K27ac (**Fig. S3B**), H3K4me1 (**Fig. S3C**), and H3K4me3 (**Fig. S3D**) compared to those retrieved in the A, A-, and S groups. In contrast, the integration site of the provirus in clone #814 was relatively distal to the histone mark H3K36me3 compared to those retrieved in clones assigned to the A, A-, and S groups (**Fig. S3E**). The integration site of the provirus in clone #814 also appeared closer to repressive histone marks, H3K9me3 (**Fig. S3F**) and H3K27me3 (**Fig. S3G**), compared with those identified in clones assigned to the A, A-, and S groups. Altogether, the provirus integration site in clone #814 was in the vicinity of the SINEs’ and Alu elements’ repetitive regions associated with H3K27ac and H3K4me3 histone modifications.

### 4.5 The involvement of HIV ASTs in characterizing distinct transcriptional phenotypes across sinpro clones

Although, at present, the fundamental mechanism behind the role of HIV ASTs and its consequence is not fully understood, accumulating evidence has shown the potential interference of HIV ASTs with its sense RNA transcription [48,52–59,71]. Here, we further hypothesize that HIV ASTs may also be involved in the regulation of stochastic transcription of the proviruses through the competition between sense and antisense transcripts. In order to independently measure sense and antisense transcription, we applied the method of strand-specific reverse transcription (RT), followed by PCR amplification, which was previously published by Landry et al. [49], with modifications (**Fig. S4**; see the **Material and methods**). We first verified the expression of antisense RNAs in an HIV-based vector [23,57] in the sinpro clones; then, we observed the PCR products resulting from RT in the antisense direction (**Fig. S4B**, lanes 10 and 11). The size of the antisense transcripts was no longer detectable starting from the middle of the gene encoding GFP in the antisense direction (**Fig. S4B**, lanes 7 and 8). To increase the specificity of this assay, we redesigned the RT primers, allowing for independent amplification of the sense (HCP#266) and antisense (HCP#264) transcripts (**Fig. S4C**), optimized the RT-PCR condition (**Fig. S4D**), and confirmed the specificity of the PCR product (**Fig. S4E**). In parallel, we implemented a strand-specific RT assay, followed by digital PCR (dPCR), enabling an absolute quantitative measurement of RNA molecules at near-single-molecule resolution (**Fig. S4F**). The dose-dependent (10X and 100X) detection of sense and antisense transcripts was achieved across 12 sinpro clones at three time points (**Fig. S4G**). Altogether, these observations indicate the feasibility of using our methods to independently measure sense and antisense transcripts.

In a follow-up study, we applied strand-specific RT, followed by qPCR and dPCR, on nine clones, including clone #812 in the A group, three clones (#789, #795, and #802) in the A- group, four clones (#778, #788, #800, and #815) in the S group, and clone #814, to observe the sense and antisense RNA transcription at six subsequent time points (**Fig. 3A** and **3B**; **Fig. S5**) and to match them to corresponding FACS measurements recorded at the same time (**Fig. S6**).

Given that the RNAs were isolated from the total cell population, here, we only took into account the parameters measured from a population of total live cells. We observed a positive correlation for GFP-fluo-total versus either sense or antisense RNAs measured via qPCR (**Fig. S6A** and **S6B**) and dPCR (**Fig. S6C** and **S6D**), respectively (dotted lines of linear regression shown in each panel). Moreover, a positive correlation between sense and antisense RNAs was observed over time (**Fig. 3A** and **3B**). A similar trend for sense and antisense RNA transcription is presented in **Fig. S5**. Although the magnitude of transcriptional bursting varied across individual clones, based on our methods, the abundance of sense transcripts was about 15 to 20 times (qPCR: mean, 28.32 and median, 12.51; dPCR: mean, 20.83 and median, 9.69) higher than that of antisense RNAs in the sinpro clones. Altogether, these findings suggest that fluctuations in HIV transcription in the sinpro clones most likely resulted from the transcriptional competition between sense and antisense transcripts, rather than alternative transcription between each other.

Intriguingly, while assaying the ratio between sense and antisense transcripts for clones in different groups, no difference was observed (**Fig. 3C**). However, as we independently assayed the abundance of sense and antisense transcripts, an obvious deterministic threshold level was obtained (**Fig. 3D**, qPCR and **3E**, dPCR). Using both methods, a superior abundance of sense and antisense transcripts was observed in clones #812 and #814, followed by clones in the A- and S groups.

We further sought the threshold for determining HIV transcription in sinpro clones. We computed the minimum (Min.), lower quartile (Q1), median (Q2), upper quartile (Q3), and maximum (Max.) values, representing the ensemble of provirus transcripts in clones in each group, and observed a clear separation between clone #812 (the A group), clone #814 (the F group), and other clones (the A- and S groups) (**Fig. 3F-3I**). As the Min. measured via qPCR was over 0.1 (arbitrary unit, a.u.; **Fig. 3F**, sRNAs) and 0.05 (a.u.; **Fig. 3G**, asRNAs), the clone tended to be in the A group. Using dPCR, a clear separation between the A group and other clones was observed at Q1, with values over 2 (a.u.; **Fig. 3H**, sRNAs) and 0.25 (a.u.; **Fig. 3I**, asRNAs). In all circumstances, the separation of clones between the A- and the S groups faltered: a visible separation lay at Q3 and Max. (**Fig. 3F-3I**). The abundance of sense and antisense transcripts measured in clone #814 either revolved around the threshold generated based on clone #812 using qPCR (**Fig. 3F** and **3G**) or lay between the thresholds between the A and A- groups (**Fig. 3H** and **3I**) using dPCR. Altogether, our results suggest that the yield of sense and antisense transcripts, rather than their ratio, determined the stochastic phenotype of provirus transcription in sinpro clones.

### 4.6 Exponential decay model demonstrated a better fit of stochastic HIV transcription in sinpro clones

To strengthen the classification of these clones with an improving systematic quantification of stochastic variations across sinpro clones, we further implemented the mathematical model previously proposed by Singh et al. [9] to fit their transcriptional phenotypes. As demonstrated in their study, in most of the clones infected by minimal lentiviral vector noise aligned with the equation *CV*^2^ = *C* /〈*m*〉 (Equation 1), where *C* is the proportionality factor and 〈*m*〉 is the average number of GFP molecules per cell (〈〉denotes the average) [9,68] with the proportionality factor in the range between 15,000 and 65,000 [9]. In this study, we call this equation the “reciprocal curve model” (**Fig. 4B**) in order to distinguish it from our proposed model described below. While plotting the correlation between *CV*^2^ and GFP fluorescence across 32 clones in the A, A-, and S groups (**Fig. 4A**), a lack of dependence between these two variables was observed (**Fig. 4B**, R^2^ = 0). A fragile correlation between these two parameters across eight clones in different groups was also observed as measurements were performed in a time series (**Fig. S7A**-**S7F**).

We wondered about the cause of this observation. It might have resulted from the relatively few input data in our experimental setting compared with that in the previous studies that investigated stochastic variation in HIV gene expression [9,33] and in other organisms [68]. In an attempt to seek models that may better fit HIV transcription in sinpro clones, we formulated an equation of exponential decay, namely the “exponential decay model” (**Fig. 4C**)

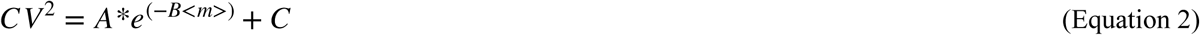

Where *CV* and *m* are identical to the parameters present in Equation 1; *e* is Euler’s number; and *A*, *B*, and *C* are fitted constants (**Fig. 4E**). Intriguingly, we observed a slight redistribution of clones across the A, A-, and S groups, whereas clone #814 remained independent from other clones (**Fig. 4C**). Based on the curve fitting using the exponential decay model, five new models were identified: (1) model I—clones in which a relatively high degree of transcriptional bursting (as represented by *CV*^2^) was detected (R^2^ = 0.77), (2) model II—clones in which moderate transcriptional bursting was detected (R^2^ = 0.61), (3) model III—clones in which relatively low transcriptional bursting was detected (R^2^ = 0.33), (4) model IV—the identical clones previously assigned to the S group (R^2^ = 0), and (5) model V—clone #814 (R^2^ = 0.58) (**Fig. 4C**). Here, we note that no convincing model can be constructed for the clones in model IV because of the wide variety of transcription noise and the extremely low level of GFP expression, resulting in the appearance of a vertical line aligning all data points (**Fig. 4C**). In addition, as we FACS-sorted the GFP_Bright_ and GFP_Dim_ subpopulations from clone #814, both subpopulations fell into the curve of model V (**Fig. 4C**). This observation somehow support our previous conclusion that the mechanism governing transcriptional bursting in clone #814 could be inheritable at a minimal level. Individual clones assigned to the corresponding models are listed in **Fig. 4C**.

Noticeably, here, better discrimination across clones in the A and A- groups was observed using the exponential decay model. Clones #758, #764, and #802, in which a high degree of noise was measured (model I), were clustered in the vicinity of those in model IV in the network property due to a high degree of noise appearing in clones in both models (**Fig. 4D**), whereas the wide distribution of clones in model III may reflect a relative weakness of curve fitting (R^2^ = 0.33) given such the low degree of measured noise (**Fig. 4C** and **4D**). Although the exponential model representing clone #814 remained isolated from others (**Fig. 4C**), the subpopulation of #814-GFP_Dim_, in which higher noise was measured over time, tended to be close to the clones in models I and IV (**Fig. 4D** and **4F**), whereas the unsorted #814 and the #814-GFP_Bright_ subpopulations, in which relatively low levels of noise were measured (**Fig. 4F**), were embedded in the clones in model II (**Fig. 4D**). The A, B, and C constants are summarized in **Fig. 4E**. Nevertheless, these findings suggest that the exponential decay model enabled a better classification of sinpro clones, in which the proviruses demonstrated distinct levels of transcriptional bursting. In addition, this result, once again, highlighted that the mechanism governing stochastic HIV transcription in clone #814 may differ from that regulating provirus transcription in the clones in the other models.

To verify whether HIV sense and antisense transcription can also be depicted via the exponential decay model constructed based on GFP expression, we applied measures of sense and antisense RNAs from nine clones, shown in **Fig. 3**, to fit the models. The curve of exponential decay recapitulated sense RNA transcription in model I with a better fit when transcription was measured using qPCR (**Fig. 4G**, R^2^ = 0.92) than when measured using dPCR (**Fig. 4I**, R^2^ = 0.0). The robustness of curve fitting faltered in the rest of the models as 〈*m*〉was replaced with sRNAs: models II (**Fig. 4G**, qPCR, R^2^ = 0.52; **Fig. 4I**, dPCR, R^2^ = 0.0), III (**Fig. 4G**, qPCR, R^2^ = 0.17; **Fig. 4I**, dPCR, R^2^ = 0.0), IV (**Fig. 4G**, qPCR, R^2^ = 0.009; **Fig. 4I**, dPCR, R^2^ = 0.0), and V (**Fig. 4G**, qPCR, R^2^ = 0.0; **Fig. 4I**, dPCR, R^2^ = 0.0). Antisense RNA transcription measured via qPCR modestly fit the curves in models II (**Fig. 4H**, R^2^ = 0.21) and III (**Fig. 4H**, R^2^ = 0.67). Similar to the observation in sRNAs, a poor overall fit was observed when antisense transcription was measured using dPCR (**Fig. 4J**). These findings imply that either different promoter activities occur between the HIV 5’LTR and 3’LTR or perhaps additional parameters are required to better describe the transcriptional phenotype of asRNAs. Of note, we also verified the effectiveness and robustness of our models using GFP expression measured via flow cytometry in the same nine clones: model I remained robust regarding its fit to the provirus transcription in clone #802 based on the GFP expression at six subsequent time points (**Fig. S7G**, R^2^ = 0.99), whereas semi-robustness of models II (R^2^ = 0.43) and V (R^2^ = 0.35) was observed (**Fig. S7G**). A poor fit was observed in models III (**Fig. S7G**, R^2^ = 0.0) and IV (**Fig. S7G**, R^2^ = 0.04). We assumed that this phenomenon was due to either the few clones included here or the few time points used for the measurements. Clones in which proviruses demonstrated a high degree of noise and a low level of transcription (clones #778, #788, #800, and #815), once again, failed to formulate a curve of exponential decay (**Fig. S7G**). Altogether, the proposed exponential decay model with the parameter measured at the protein level (GFP expression) enabled better recapitulation of the sinpro clones, in which the proviruses demonstrated distinct transcriptional phenotypes. The models were fragile at the transcription level, especially the test fitting asRNAs, which might be due to the weaker promoter activity of the HIV 3’LTR and a higher background noise level in a measurement at the RNA level compared with that at the protein level.

### 4.7 Both HIV LTRs were responsive to drug stimulation

As a final step, we wondered whether the HIV 5’ and the 3’LTRs that drive sense and antisense RNA transcription, respectively, behave in the same manner while clones are subjected to different activators and drugs. To test this, we continuously treated the sinpro clones used in the previous experiments, as well as #814-GFP_Bright_ and #814-GFP_Dim_ subpopulations, with phorbol 12-myristate 13-acetate (PMA) plus ionomycin (hereinafter, “PMA/Ionomycin”), ingenol-3-angelate (PEP005), a PKC activator, or dimethyl sulfoxide (DMSO) as a negative control (**Fig. S8A**-**S8I**). The fact that the proviruses in these clones and in two #814 subpopulations remained functional and were responsive to PMA/Ionomycin and PEP005 was evidenced by an increase in the measures of (%) GFP(+), GFP-fluo-gate, and GFP-fluo-total (**Fig. S8A**-**S8I**). The reactivation shown in clone #814 and its two subpopulations suggests that the HIV 5’LTR remained transcriptionally accessible despite the presence of the integration site proximal to the genomic repetitive regions (**Fig. 2M**).

Intriguingly, we observed that proviruses in clones assigned to different groups and models showed different responses to different drugs (**Fig. S9**). In clones #812 (**Fig. S9A**) and #789 (**Fig. S9B**), provirus reactivation tended to elevate the mean GFP expression (GFP-fluo-total), showing a negative but linear relationship with transcription noise [represented by (pb-CV-total)^2^]. In contrast, in clones #795 (**Fig. S9C**) and #802 (**Fig. S9D**), provirus reactivation tended to synergetically affect the mean GFP expression and transcription noise, which showed a positive but linear relationship. Of note, although both pairs of clones— #812 and #789, and clones #795 and #802—were assigned to different groups or models, clones in the same pair demonstrated topological proximity to the network property (**Fig. 4D**). This may partially explain their similar response toward drugs. The reactivation pattern of the provirus in the S group was more diverse (**Fig. S9E**-**S9H**). We assumed that this phenomenon might be due to the relatively high intrinsic background noise (shown in the Mock 1 and 2 controls and cells treated with DMSO) that lasted until the proviruses were reactivated. At present, it remains challenging for us to tell whether the intense magnitude of noise shown in reactivated proviruses in the S group resulted as an effect of drugs or was due to the viruses themselves. Different reactivation patterns were observed between clone #814 and its subpopulations (**Fig. S9I**): unsorted clone #814 and #814-GFP_Dim_ showed similar reactivation patterns to those observed for clones #812 and #789 (**Fig. S9I**), whereas all data points from #814-GFP_Bright_ shifted to the lower right quadrant (**Fig. S9I**). In the former group, the presence of PMA/Ionomycin or PEP005 enhanced the mean GFP expression, and the presence of DMSO enhanced transcription noise (**Fig. S9I**). In the latter case, the enrichment of the mean GFP expression was comparable between drug reactivation and the DMSO control (**Fig. S9I**).

At the RNA level, the enrichment of both sense and antisense transcripts appeared in the presence of PMA/ Ionomycin or PEP005 across clones in all groups (**Fig. 5A-5H**). A more intense augmentation was shown in sRNAs compared with asRNAs, reflecting a general expectation that the HIV 5’LTR activity should be superior to that of 3’LTR. Intriguingly, we observed that in clone #814, the provirus preferentially responded to PEP005, especially the 5’LTR that drives sense RNA transcription (**Fig. 5H**); an extremely weak enrichment for both sense and antisense transcripts against the presence of PMA/Ionomycin (**Fig. 5H**). Notably, although the overall abundance of asRNAs was less than that of sRNAs, we observed that the comparable enrichment between both sense and antisense RNAs when the fold changes were normalized by measures from clonal cells treated with DMSO in the A, A-, and S groups (**Fig. 5I**), suggesting that the activity of two HIV LTRs from the proviruses in clones assigned to these three groups were inducible irrespective of the presence of PMA/Ionomycin or PEP005. Although a tendency that two HIV LTRs were more susceptible to PMA/Ionomycin than PEP005 was observed in proviruses in clone #812 (the A group) (**Fig. 5I**), a bias resulting from few data points here should also be taken into account. Notably, a discrepancy between regression lines formed by the enrichments of sense and antisense transcripts in clone #814 (the F group) indicated that the 5’LTR was more susceptible to PEP005, whereas the 3’LTR was more sensitive to PMA/Ionomycin (**Fig. 5I**). Nevertheless, these findings offered a proof-of-concept that both HIV LTRs are responsive to provirus reactivation in the presence of different drugs for provirus reactivation.

**Fig. 5.**
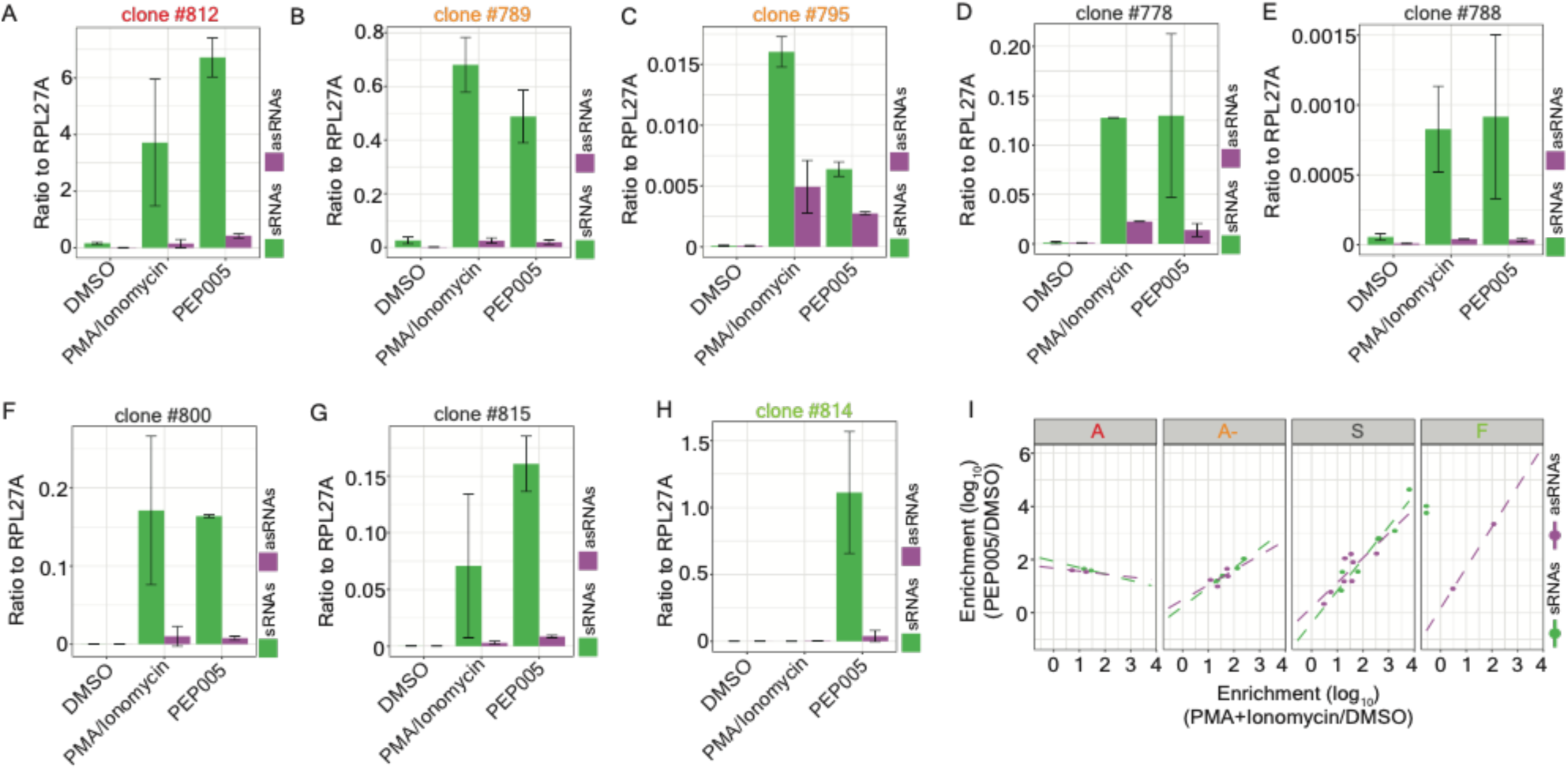
Both HIV LTRs were responsive to drug stimulation. (**A**-**H**) Bar charts representing the fold change (drugs versus DMSO) between sense and antisense RNA transcripts measured via strand-specific RT-qPCR (ratio to RPL27A) in sinpro clones treated with PMA/ Ionomycin or PEP005. Sipro clones used in this assay include clones (**A**) #812 (the A group), (**B**) #789 (the A- group), (**C**) #795 (the A- group), (**D**) #778 (the S group), (**E**) #788 (the S group), (**F**) #800 (the S group), (**G**) #815 (the S group), and (**H**) #814 (the F group). Results are the mean ± standard deviation (SD) of two independent experiments with duplicate samples. Bar in green and magenta represent the measures from sense and antisense transcripts, respectively. (**I**) Scatter plots representing the correlation between the enrichment of sense and antisense transcripts between sinpro clones treated with PMA/Ionomycin (x-axis) and those treated with PEP005 (y-axis). The enrichment is calculated using strand-specific RT-qPCR readouts from cells treated with drugs (PMA/Ionomycin or PEP005) normalized by readouts from cells treated with DMSO. Experiments were carried out from two independent biological replicates with duplicate samples. Both axes are in logarithmic scale. Dots and dotted lines marked in green and magenta represent measures from sense and antisense transcription, respectively.

## 5. Discussion

The stochastic expression of HIV proteins immediately after infection has been shown to be able to critically influence viral fate following active replication or post-integration latency [8,9,36,72]; however, the source of this noise is not fully understood. Mechanisms, including (1) stochastic fluctuations in the expression of the HIV Tat protein (Tat transcriptional pulses) [8,35,36,72], (2) the selection of HIV integration sites associated with epigenetic features [10,21–23], (3) RNAPII pausing [73], and (4) chromatin architecture and nucleosome occupancy [74], were reported to contribute to stochastic HIV transcription.

In this study, we proposed that stochastic fluctuations in HIV transcription can appear at, at least two levels: (1) the chromosomal landscape and (2) the in situ HIV integration site (**Fig. 6**) based on an experimental setting with 33 sinpro clones established in an immortalized Jurkat T cell line. The former case, represented by sinpro clones in the A, A-, and S groups (**Fig. 1A** and **Fig. S3**), resonated with previous findings, indicating that the level of provirus transcription differs between one provirus and another and can be influenced by the local genomic context surrounding HIV integration sites coupled with genomic features (e.g., chromatin accessibility) [21–23] and histone modifications [23,75]. Our results further set the stage for advancements in the classification of proviruses with varying levels of stochastic transcription in noise space (**Fig. 1D** and **1F**) or based on mathematical modeling (exponential decay models) (**Fig. 4C**).

**Figure 6.**
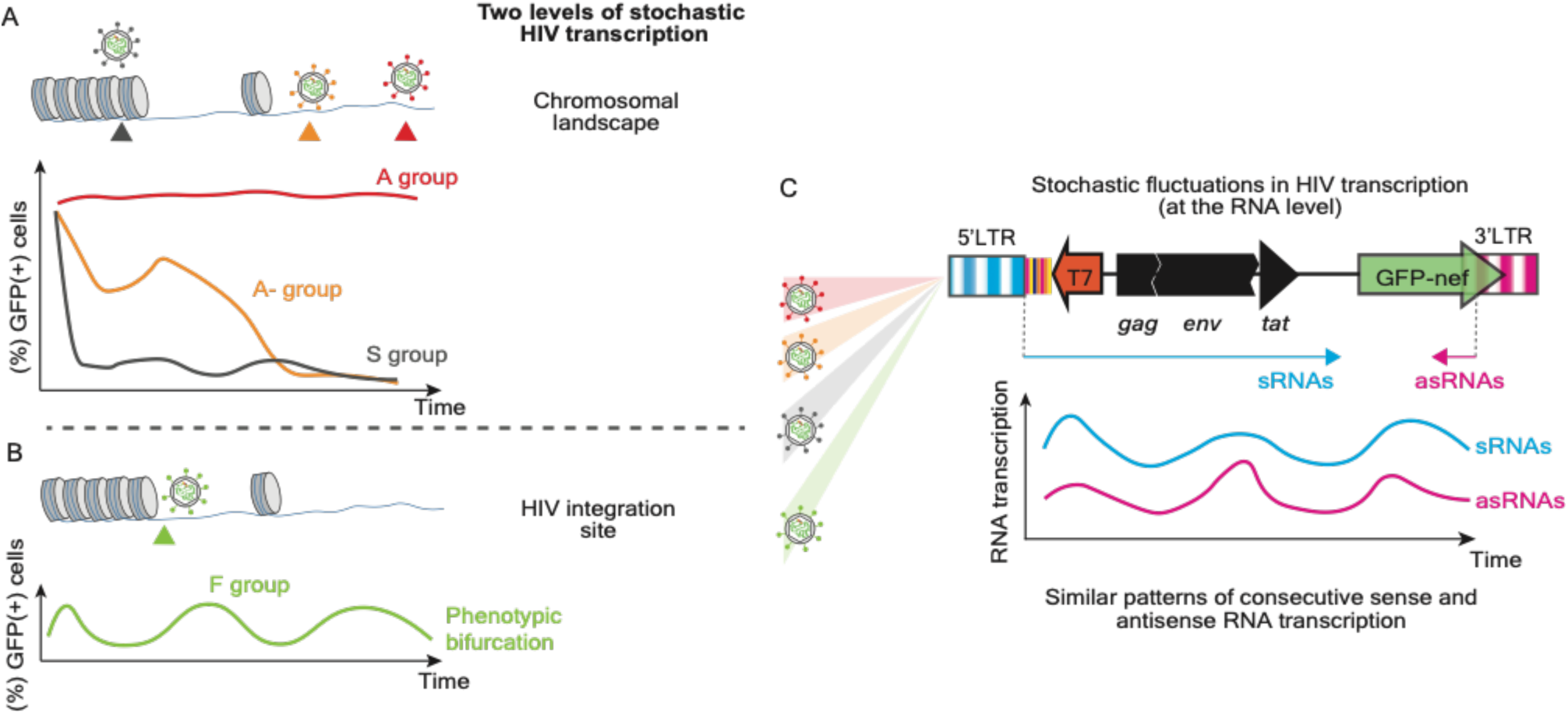
The involvement of HIV asRNAs in stochastic fluctuations in HIV transcription. We propose that fluctuations in HIV transcription can appear at, at least two levels: (1) the chromosomal landscape (**A**) and (2) the in situ HIV integration site (**B**). In the former case, proviruses integrating in different genomic locations (sinpro clones in the A, A-, and S groups) demonstrate different levels of HIV transcription coupled with a variety of transcriptional bursting (**A**). In the latter case represented by clone #814 (the F group), stochastic HIV transcription can be unveiled through its phenotypic bifurcation and tends to be a pure epigenetic phenomenon—the identical provirus demonstrates the turnover of HIV transcription with elevated frequency (**B**). We observed a minimum level of inheritance of such stochastic fluctuations (the provirus in the F group) and identified the provirus integration site, which is in the vicinity of the genomic repetitive regions associated with intense signals of H3K27ac and H3K4me3. In both cases, similar expression patterns between sense and antisense RNA transcripts in a time series were observed (**C**), suggesting that the independent yields of both transcripts determine the stochastic phenotype of provirus transcription, rather than their ratios.

In the latter case, represented by clone #814 (**Fig. 2A**), the turnover of HIV transcription (active versus silent) showed elevated frequency (**Fig. 2B**). In other words, stochastic fluctuations in HIV transcription in this circumstance are unveiled through its phenotypic bifurcation [measures of (%) GFP(+) cells] (**Figures S1A** and **2B**) and should be a pure epigenetic phenomenon, given that the provirus is identical regardless of the state of HIV transcription. We observed relatively intense signals of H3K27ac and H3K4me3 (**Fig. 2N**) within the 10 kb genomic region, suggesting that the HIV-1 5’LTR is most likely transcriptionally accessible even though the integration site was identified in the vicinity of SINEs and Alu elements (**Fig. 2M**). This conclusion can be supported by the provirus reactivation assay on clone #814 using PMA/Ionomycin or PEP005 (**Fig. S8I**). Heterochromatin is presently considered one of the genomic hubs where (deeply) latent proviruses persist via immune-mediated selection [76–80]. Future studies will be required to explore the correlation between clones harboring the same transcriptional phenotype as clone #814 and integration near repetitive genomic regions. Interestingly, unintegrated HIV DNA was proposed to adopt a repressive chromatin structure that competes with the transcription mechanism, leading to its silencing [81,82]; whether HIV antisense transcription can also be manifested around 2-LTR circles of unintegrated HIV DNA is presently unknown and will be studied in the future. It is also important to note that, although at present it remains unclear how frequent clones with phenotypic bifurcation are present in the HIV reservoir, the appearance of such clones most likely reflects the outcome of a frequent competition between sense and antisense transcription and/or a completion versus disruption of the Tat transactivation circuit. Whether they are susceptible to viral rebounds after ART is interrupted and viral blips observed in elite controllers and in HIV-infected individuals under prolonged ART will be important for further investigation.

Notably, the phenotype of clone #814 resembles a subset of clonal populations that generated a variegated expression phenotype, namely phenotypic bifurcation (PheB), which can exhibit two subpopulations, one with bright GFP fluorescence and another GFP fluorescence-free [8]. As we separated the GFP_Bright_ subpopulation of clone #814 from its GFP_Dim_ subpopulation, we observed a minimal level of inheritance of such stochastic fluctuations (**Fig. 2C**; **Fig. S2F** and **S2G**): #814-GFP_Bright_ and #814-GFP_Dim_ were indistinguishable in noise space (**Fig. 2D** and **2F**) and network properties (**Fig. 2H** and **2J**). The clear separation between these two subpopulations based on the measures of (%) GFP(+) cells may suggest unequal epigenetic memory between these two subpopulations. Nevertheless, cumulative studies have shown that long-term epigenetic inheritance mainly occurs in broad heterochromatin domains [83,84]; further investigations into these two subpopulations at higher order chromatin architectures will be required to gain more insights into the molecular mechanism underlying provirus transcription in such a clone.

In this study, we further hypothesized that HIV ASTs may also be involved in mechanisms that govern this stochastic phenotype of HIV transcription, perhaps subsequently promoting the establishment of its latency. Indeed, viruses have developed various strategies to establish a latent state to ensure their persistence in their hosts. For instance, herpes viruses encode latency proteins that are responsible for switching the viral cycle from a lytic to a latent state [85]. Another example is the detection of ASTs, namely CMV latency-associated transcripts in bone marrow aspirating from naturally infected, healthy seropositive donors infected by human cytomegalovirus [86]. Retroviruses, in contrast, are known to persist through transcriptional silencing and do not encode latency proteins. In fact, in other human retroviruses, like human T-cell leukemia virus type 1 (HTLV-1), one of the best characterized ASTs in a viral system, its antisense gene product, HTLV-1 basic leucine zipper factor (HBZ), has been proven to functionally promote the survival and proliferation of HTLV-1-infected cells, thereby influencing progression into adult T-cell leukemia/lymphoma or HTLV-1-associated myelopathy/tropical spastic paraparesis [87]. HIV ASTs are supposed to suppress sense RNA transcription; however, at present, the fundamental mechanism behind the competition between sense and antisense transcripts and its biological consequences require further characterization. A comprehensive understanding of the role of HIV ASTs involved in the regulation of HIV transcription may not only unveil an unprecedented mechanism used by HIV to establish a persistent infection but also reinforce concepts related to the fundamental mechanism of antisense RNA transcription.

To date, HIV ASTs have been demonstrated *in vitro* to be capable of promoting the initiation and maintenance of HIV latency by recruiting polycomb repressor complex 2 or chromatin modifying enzymes to the 5’LTR of the provirus, resulting in repression of the expression of their sense RNAs [52–54]. However, the expression of HIV ASTs *in vivo* has not been convincingly demonstrated, and only a few studies have examined its expression. Zapata et al. [53] reported low levels of ASTs in three people living with HIV on ART using real-time PCR in unstimulated CD4+ T cells. Mancarella et al. [71] detected ASTs in CD4+ T cells collected from both untreated and ART-treated patients after *ex vivo* stimulation with anti-CD3/CD28. Recently, a preprint reported by Capoferri et al. [88] provided sequence evidence for the expression of ASTs in unstimulated peripheral blood mononuclear cells collected from people living with HIV.

Presently, the fundamental mechanism behind the mechanistic interplay between HIV sense and antisense transcripts is unclear. In this study, we performed strand-specific RT followed by qPCR and dPCR to independently measure sRNAs and asRNAs in sinpro clones over time (**Fig. 3A** and **3B**; **Fig. S5**). We observed similar expression patterns (**Fig. S5**) and a positive correlation between the abundance of sRNAs and asRNAs (**Fig. 3A** and **3B**), indicating that, in sinpro clones, both the HIV 5’ and 3’LTR are most likely active, whereas their strengths differ. This phenomenon may partially align with the previous study, indicating a predominant usage of the 5’LTR promoter by HIV (so-called “promoter suppression” in this study) [89]; however, in our study, we observed an indistinguishable ratio between sRNAs and asRNAs across sinpro clones (**Fig. 3C**). We extrapolate that a time lag of the antisense transcription driven by the 3’LTR might be one of the possible causes that lead to a similar expression pattern between sense and antisense transcription in sinpro clones. Nevertheless, in our setting, based on the measurements conducted via qPCR and dPCR, more sense transcripts than antisense transcripts with clear threshold levels were detected (**Fig. 3D-3I**), which aligned with previous findings [48,53,90]. Intriguingly, Capoferri et al. [88] observed that in the subgenomic region of the HIV *env* gene, more HIV ASTs than their sense transcript counterparts can be targeted, highlighting that perhaps diverse transcript abundances between sRNAs and asRNAs are present throughout the HIV genome.

It is worth noting that whether the two-state promoter model (ON versus OFF) can completely capture stochastic fluctuations of HIV transcription at the transcription and the protein levels, especially antisense transcription, is presently unclear. It has been evidenced that perhaps the HIV Tat positive-feedback loop tends to be a monostable [8] or atypical two-state model with a transient threshold [35,36]. The robustness of our exponential decay models also faltered when sRNAs and asRNAs were applied (**Fig. 4G-4J**), implying the need for additional factors to understand the entire regulatory mechanism. Nevertheless, further investigations into the interaction between sRNAs and asRNAs of individual genes will be needed.

The shock-and-kill strategy attempts to reactivate latent proviruses back into active replication states and to simultaneously purge HIV infections via ART [91,92]. Despite remarkable efforts, expectations for this shock-and-kill strategy have not been met yet [93]. As previously proposed, such limited success may be due to the stochastic nature of latency [8,36,94–96]. The study from Li et al. [97] suggested that the limited efficacy of latency reversing agents may be due to concomitant induction of HIV ASTs, thwarting their effect on sense transcription. To compensate for the limitations of this strategy, alternative approaches are required. In the recent study reported by Li et al. [56], the ectopic expression of HIV ASTs in CD4+ T cells from people living with HIV under ART can block latency reversal in response to pharmacologic and T-cell receptor stimulation, underscoring the potential of HIV ASTs to be used in the block-and-lock functional cure strategy. Our results, as analyzed from another aspect, implied that both HIV LTRs were responsive to drug stimulation (**Fig. 5A-5H**) and may react with different strengths from proviruses in sinpro clones assigned to different groups at both the protein (**Fig. S9**) and RNA (**Fig. 5I**) levels. Indeed, it is known that two HIV LTRs possess different activities [89] and distinct binding sites of transcription factors [41]; however, how they respond to drug reactivation remains unclear. The distinct effectiveness of curve fitting in exponential decay models observed between sense and antisense RNAs also supports the assumption that different promoter activities between two HIV LTRs exist (**Fig. 4G-4J**). Distinct behaviors between 5’ and 3’LTRs of the provirus in clone #814 highlighted the opportunity that perhaps in unique scenarios, e.g., pure epigenetics-driven stochastic fluctuations, two HIV LTRs may respond to stimuli in different ways (**Fig. 5I**). In addition, from clinical perspectives, a discrepancy in promoter activity between two HIV LTRs stimulated by different latency reversing agents may suggest that the use of complementary drugs, enabling wide reactivation spectra of latent proviruses, is imperative. A more precise investigation into the reactivity and the affinity between two HIV LTRs will be critical in determining the threshold for either increasing the efficacy of latency reversing agents by minimizing the transcription level of HIV ASTs or permanently locking proviruses in a state of (deep) latency by increasing the transcription of ASTs. We stress, as a previous study indicated, that drugs with different functionalities influence provirus transcriptional bursting differently [96]: transcriptional activators, such as tumor necrosis factor, are prone to increase burst frequency [9,10], whereas chromatin remodelers, such as histone deacetylase inhibitors, methylation inhibitors, and bromodomain inhibitors tend to cause modest increases in burst size [96]. Nevertheless, elucidating the transcriptional behaviors of proviruses in the presence of different drugs may result in better designed antiretroviral drugs and strategies.

### The limitations of this study

This study was based on an experimental setting with 33 sinpro clones in total. Although we cannot rule out any bias introduced by these small-scale cellular models, we demonstrated a clear separation between sinpro clones in noise space and using exponential decay models. Since latent HIV infections can be established at the early stage of infection [37–39], we note that our strategy for screening of sinpro clones cannot cover the clones, in which proviruses have reached the state of latency before the screening was conducted. In addition, biological observations based on this cellular model, which was set up in an immortalized T cell line, may differ from studies conducted in primary CD4+ T cells, especially resting and unstimulated cells. A head-to-head fashion of two LTRs in HIV-based vector may not fully recapitulate the stochastic nature of wildtype HIV.

It is encouraging that a unique provirus transcriptional phenotype, shown in clone #814, can be identified. Such a clone may not be simply identified if a pool of infected cells is studied although only one clone harboring such a unique phenotype was obtained. We have confirmed the stability of the stochastic phenotype of provirus transcription in clone #814 based on two independent measurements in a time series: one was performed in 2023 (**Fig. 2B**) and another in 2025 (data not shown). In recent measurements, the phenotype reappeared, but faltered. At present, we cannot answer whether the flickering of such a phenotype will fade with time. And if so, does it result from multiple rounds of cell manipulation (freeze and thaw cells) or inheritance of unequal epigenetic memory in daughter cells, as previously discussed. More clones, in which proviruses demonstrate the same phenotype of stochastic transcription as that in clone #814 will be requisite to tackle this question. Finally, we cannot offer single-cell measurements for parameters, as previously performed [10,33]. Further investigations in large-scale experimental settings at the single transcripts resolutions in individual cells will be necessary to verify our observations on sinpro clones.

## Acknowledgments

HCC acknowledges funding from the Narodowe Centrum Nauki (Sonata Bis Grant UMO-2022/46/E/ NZ6/00022). We would like to thank Dr. Z. Debyser (KU Leuven) for his proofreading and critical feedback on the manuscript.

## Author contributions

Conceptualization, H.-C.C.; methodology, H.-C.C., K.W. and J.W.; software, H.-C.C., and J.W., formal analysis, H.-C.C., K.W. and J.W., investigation, H.-C.C., K.W., and J.W.; resources, H.-C.C. and J.W.; data curation, H.-C.C., and J.W.; writing of original draft manuscript, H.-C.C., K.W., and J.W.; writing, manuscript review and editing, H.-C.C., K.W., and J.W.; visualization, H.-C.C., and J.W.; supervision, H.-C.C., project administration, H.-C.C.; funding acquisition, H.-C.C.

## Declaration of generative AI and AI-assisted technologies

The authors declare no generative AI and AI-assisted technologies are used in this work.

## Supplemental information index

Document S1. Figures S1–S9 and Tables S1-S5

Table S1. Excel file containing additional data too large to fit in a PDF, related to Figures 1 and 4.

Table S4. Excel file containing additional data too large to fit in a PDF, related to Figures 3 and 4.

## References

1. Thattai, M., and van Oudenaarden, A. (2001). Intrinsic noise in gene regulatory networks. Proc Natl Acad Sci U S A 98, 8614–8619. 10.1073/pnas.151588598.

2. Ozbudak, E.M., Thattai, M., Kurtser, I., Grossman, A.D., and van Oudenaarden, A. (2002). Regulation of noise in the expression of a single gene. Nat Genet 31, 69–73. 10.1038/ng869.

3. Paulsson, J. (2004). Summing up the noise in gene networks. Nature 427, 415–418. 10.1038/nature02257.

4. Raser, J.M., and O’Shea, E.K. (2004). Control of stochasticity in eukaryotic gene expression. Science 304, 1811–1814. 10.1126/science.1098641.

5. Becskei, A., Kaufmann, B.B., and van Oudenaarden, A. (2005). Contributions of low molecule number and chromosomal positioning to stochastic gene expression. Nat Genet 37, 937–944. 10.1038/ng1616.

6. Colman-Lerner, A., Gordon, A., Serra, E., Chin, T., Resnekov, O., Endy, D., Pesce, C.G., and Brent, R. (2005). Regulated cell-to-cell variation in a cell-fate decision system. Nature 437, 699–706. 10.1038/nature03998.

7. Rosenfeld, N., Young, J.W., Alon, U., Swain, P.S., and Elowitz, M.B. (2005). Gene regulation at the single-cell level. Science 307, 1962–1965. 10.1126/science.1106914.

8. Weinberger, L.S., Burnett, J.C., Toettcher, J.E., Arkin, A.P., and Schaffer, D.V. (2005). Stochastic gene expression in a lentiviral positive-feedback loop: HIV-1 Tat fluctuations drive phenotypic diversity. Cell 122, 169–182. 10.1016/j.cell.2005.06.006.

9. Singh, A., Razooky, B., Cox, C.D., Simpson, M.L., and Weinberger, L.S. (2010). Transcriptional bursting from the HIV-1 promoter is a significant source of stochastic noise in HIV-1 gene expression. Biophys J 98, L32–L34. 10.1016/j.bpj.2010.03.001.

10. Dar, R.D., Razooky, B.S., Singh, A., Trimeloni, T.V., McCollum, J.M., Cox, C.D., Simpson, M.L., and Weinberger, L.S. (2012). Transcriptional burst frequency and burst size are equally modulated across the human genome. Proc. Natl. Acad. Sci. U. S. A. 109, 17454–17459. 10.1073/pnas.1213530109.

11. Dornadula, G., Zhang, H., VanUitert, B., Stern, J., Livornese, L., Jr, Ingerman, M.J., Witek, J., Kedanis, R.J., Natkin, J., DeSimone, J., et al. (1999). Residual HIV-1 RNA in blood plasma of patients taking suppressive highly active antiretroviral therapy. JAMA 282, 1627–1632. 10.1001/jama.282.17.1627.

12. Ishizaka, A., Sato, H., Nakamura, H., Koga, M., Kikuchi, T., Hosoya, N., Koibuchi, T., Nomoto, A., Kawana-Tachikawa, A., and Mizutani, T. (2016). Short Intracellular HIV-1 Transcripts as Biomarkers of Residual Immune Activation in Patients on Antiretroviral Therapy. J Virol 90, 5665–5676. 10.1128/JVI.03158-15.

13. Yukl, S.A., Kaiser, P., Kim, P., Telwatte, S., Joshi, S.K., Vu, M., Lampiris, H., and Wong, J.K. (2018). HIV latency in isolated patient CD4 T cells may be due to blocks in HIV transcriptional elongation, completion, and splicing. Sci Transl Med 10. 10.1126/scitranslmed.aap9927.

14. Fischer, M., Günthard, H.F., Opravil, M., Joos, B., Huber, W., Bisset, L.R., Ott, P., Böni, J., Weber, R., and Cone, R.W. (2000). Residual HIV-RNA levels persist for up to 2.5 years in peripheral blood mononuclear cells of patients on potent antiretroviral therapy. AIDS Res Hum Retroviruses 16, 1135–1140. 10.1089/088922200414974.

15. Günthard, H.F., Havlir, D.V., Fiscus, S., Zhang, Z.Q., Eron, J., Mellors, J., Gulick, R., Frost, S.D., Brown, A.J., Schleif, W., et al. (2001). Residual human immunodeficiency virus (HIV) Type 1 RNA and DNA in lymph nodes and HIV RNA in genital secretions and in cerebrospinal fluid after suppression of viremia for 2 years. J Infect Dis 183, 1318–1327. 10.1086/319864.

16. Halvas, E.K., Joseph, K.W., Brandt, L.D., Guo, S., Sobolewski, M.D., Jacobs, J.L., Tumiotto, C., Bui, J.K., Cyktor, J.C., Keele, B.F., et al. (2020). HIV-1 viremia not suppressible by antiretroviral therapy can originate from large T cell clones producing infectious virus. J Clin Invest 130, 5847–5857. 10.1172/JCI138099.

17. White, J.A., Wu, F., Yasin, S., Moskovljevic, M., Varriale, J., Dragoni, F., Camilo-Contreras, A., Duan, J., Zheng, M.Y., Tadzong, N.F., et al. (2023). Clonally expanded HIV-1 proviruses with 5’-leader defects can give rise to nonsuppressible residual viremia. J Clin Invest 133. 10.1172/JCI165245.

18. Ferdin, J., Goričar, K., Dolžan, V., Plemenitaš, A., Martin, J.N., Peterlin, B.M., Deeks, S.G., and Lenassi, M. (2018). Viral protein Nef is detected in plasma of half of HIV-infected adults with undetectable plasma HIV RNA. PLoS One 13, e0191613. 10.1371/journal.pone.0191613.

19. Passaes, C., Delagreverie, H.M., Avettand-Fenoel, V., David, A., Monceaux, V., Essat, A., Müller-Trutwin, M., Duffy, D., De Castro, N., Wittkop, L., et al. (2021). Ultrasensitive Detection of p24 in Plasma Samples from People with Primary and Chronic HIV-1 Infection. J Virol 95, e0001621. 10.1128/JVI.00016-21.

20. Wu, G., Zuck, P., Goh, S.L., Milush, J.M., Vohra, P., Wong, J.K., Somsouk, M., Yukl, S.A., Shacklett, B.L., Chomont, N., et al. (2021). Gag p24 Is a Marker of Human Immunodeficiency Virus Expression in Tissues and Correlates With Immune Response. J Infect Dis 224, 1593–1598. 10.1093/infdis/jiab121.

21. Jordan, A., Defechereux, P., and Verdin, E. (2001). The site of HIV-1 integration in the human genome determines basal transcriptional activity and response to Tat transactivation. EMBO J 20, 1726–1738. 10.1093/emboj/20.7.1726.

22. Jordan, A., Bisgrove, D., and Verdin, E. (2003). HIV reproducibly establishes a latent infection after acute infection of T cells in vitro. EMBO J 22, 1868–1877. 10.1093/emboj/cdg188.

23. Chen, H.-C., Martinez, J.P., Zorita, E., Meyerhans, A., and Filion, G.J. (2017). Position effects influence HIV latency reversal. Nat Struct Mol Biol 24, 47–54. 10.1038/nsmb.3328.

24. Battivelli, E., Dahabieh, M.S., Abdel-Mohsen, M., Svensson, J.P., Tojal Da Silva, I., Cohn, L.B., Gramatica, A., Deeks, S., Greene, W.C., Pillai, S.K., et al. (2018). Distinct chromatin functional states correlate with HIV latency reactivation in infected primary CD4 T cells. Elife 7. 10.7554/eLife.34655.

25. Miklík, D., Šenigl, F., and Hejnar, J. (2018). Proviruses with Long-Term Stable Expression Accumulate in Transcriptionally Active Chromatin Close to the Gene Regulatory Elements: Comparison of ASLV-, HIV- and MLV-Derived Vectors. Viruses 10. 10.3390/v10030116.

26. Vansant, G., Chen, H.-C., Zorita, E., Trejbalová, K., Miklík, D., Filion, G., and Debyser, Z. (2020). The chromatin landscape at the HIV-1 provirus integration site determines viral expression. Nucleic Acids Res 48, 7801–7817. 10.1093/nar/gkaa536.

27. Nabel, G., and Baltimore, D. (1987). An inducible transcription factor activates expression of human immunodeficiency virus in T cells. Nature 326, 711–713. 10.1038/326711a0.

28. Kinoshita, S., Su, L., Amano, M., Timmerman, L.A., Kaneshima, H., and Nolan, G.P. (1997). The T cell activation factor NF-ATc positively regulates HIV-1 replication and gene expression in T cells. Immunity 6, 235–244. 10.1016/s1074-7613(00)80326-x.

29. Perkins, N.D., Felzien, L.K., Betts, J.C., Leung, K., Beach, D.H., and Nabel, G.J. (1997). Regulation of NF-kappaB by cyclin-dependent kinases associated with the p300 coactivator. Science 275, 523–527. 10.1126/science.275.5299.523.

30. Ne, E., Palstra, R.-J., and Mahmoudi, T. (2018). Transcription: Insights From the HIV-1 Promoter. Int. Rev. Cell Mol. Biol. 335, 191–243. 10.1016/bs.ircmb.2017.07.011.

31. Verdin, E., Paras, P., Jr, and Van Lint, C. (1993). Chromatin disruption in the promoter of human immunodeficiency virus type 1 during transcriptional activation. EMBO J. 12, 3249–3259. 10.1002/j.1460-2075.1993.tb05994.x.

32. Mahmoudi, T. (2012). The BAF complex and HIV latency. Transcription 3, 171–176. 10.4161/trns.20541.

33. Austin, D.W., Allen, M.S., McCollum, J.M., Dar, R.D., Wilgus, J.R., Sayler, G.S., Samatova, N.F., Cox, C.D., and Simpson, M.L. (2006). Gene network shaping of inherent noise spectra. Nature 439, 608–611. 10.1038/nature04194.

34. Razooky, B.S., Pai, A., Aull, K., Rouzine, I.M., and Weinberger, L.S. (2015). A hardwired HIV latency program. Cell 160, 990–1001. 10.1016/j.cell.2015.02.009.

35. Aull, K.H., Tanner, E.J., Thomson, M., and Weinberger, L.S. (2017). Transient Thresholding: A Mechanism Enabling Noncooperative Transcriptional Circuitry to Form a Switch. Biophys. J. 112, 2428– 2438. 10.1016/j.bpj.2017.05.002.

36. Weinberger, L.S., Dar, R.D., and Simpson, M.L. (2008). Transient-mediated fate determination in a transcriptional circuit of HIV. Nat. Genet. 40, 466–470. 10.1038/ng.116.

37. Whitney, J.B., Hill, A.L., Sanisetty, S., Penaloza-MacMaster, P., Liu, J., Shetty, M., Parenteau, L., Cabral, C., Shields, J., Blackmore, S., et al. (2014). Rapid seeding of the viral reservoir prior to SIV viraemia in rhesus monkeys. Nature 512, 74–77. 10.1038/nature13594.

38. Colby, D.J., Trautmann, L., Pinyakorn, S., Leyre, L., Pagliuzza, A., Kroon, E., Rolland, M., Takata, H., Buranapraditkun, S., Intasan, J., et al. (2018). Rapid HIV RNA rebound after antiretroviral treatment interruption in persons durably suppressed in Fiebig I acute HIV infection. Nat Med 24, 923–926. 10.1038/s41591-018-0026-6.

39. Gantner, P., Buranapraditkun, S., Pagliuzza, A., Dufour, C., Pardons, M., Mitchell, J.L., Kroon, E., Sacdalan, C., Tulmethakaan, N., Pinyakorn, S., et al. (2023). HIV rapidly targets a diverse pool of CD4 T cells to establish productive and latent infections. Immunity 56, 653–668.e5. 10.1016/j.immuni.2023.01.030.

40. Tedbury, P.R., Mahboubi, D., Puray-Chavez, M., Shah, R., Ukah, O.B., Wahoski, C.C., Fadel, H.J., Poeschla, E.M., Gao, X., McFadden, W.M., et al. (2024). Disruption of LEDGF/p75-directed integration derepresses antisense transcription of the HIV-1 genome. bio Rxiv. 10.1101/2024.12.06.627169.

41. Bentley, K., Deacon, N., Sonza, S., Zeichner, S., and Churchill, M. (2004). Mutational analysis of the HIV-1 LTR as a promoter of negative sense transcription. Arch Virol 149, 2277–2294. 10.1007/s00705-004-0386-8.

42. Singer, M.F., Jones, O.W., and Nirenberg, M.W. (1963). The effect of secondary structure of the template activity of polyribonucleotides. Proc. Natl. Acad. Sci. U. S. A. 49, 392–399. 10.1073/pnas.49.3.392.

43. Røsok, Ø., and Sioud, M. (2004). Systematic identification of sense-antisense transcripts in mammalian cells. Nat. Biotechnol. 22, 104–108. 10.1038/nbt925.

44. Miller, D. (1988). HIV and social psychiatry. Br. Med. Bull. 44, 130–148. 10.1093/oxfordjournals.bmb.a072236.

45. Bukrinsky, M.I., and Etkin, A.F. (1990). Plus strand of the HIV provirus DNA is expressed at early stages of infection. AIDS Res Hum Retroviruses 6, 425–426. 10.1089/aid.1990.6.425.

46. Michael, N.L., Vahey, M.T., d’Arcy, L., Ehrenberg, P.K., Mosca, J.D., Rappaport, J., and Redfield, R.R. (1994). Negative-strand RNA transcripts are produced in human immunodeficiency virus type 1-infected cells and patients by a novel promoter downregulated by Tat. J Virol 68, 979–987. 10.1128/JVI.68.2.979-987.1994.

47. Tagieva, N.E., and Vaquero, C. (1997). Expression of naturally occurring antisense RNA inhibits human immunodeficiency virus type 1 heterologous strain replication. J. Gen. Virol. 78 ( *Pt 10*), 2503–2511. 10.1099/0022-1317-78-10-2503.

48. Kobayashi-Ishihara, M., Yamagishi, M., Hara, T., Matsuda, Y., Takahashi, R., Miyake, A., Nakano, K., Yamochi, T., Ishida, T., and Watanabe, T. (2012). HIV-1-encoded antisense RNA suppresses viral replication for a prolonged period. Retrovirology 9, 38. 10.1186/1742-4690-9-38.

49. Landry, S., Halin, M., Lefort, S., Audet, B., Vaquero, C., Mesnard, J.-M., and Barbeau, B. (2007). Detection, characterization and regulation of antisense transcripts in HIV-1. Retrovirology 4, 71. 10.1186/1742-4690-4-71.

50. Laverdure, S., Gross, A., Arpin-André, C., Clerc, I., Beaumelle, B., Barbeau, B., and Mesnard, J.-M. (2012). HIV-1 antisense transcription is preferentially activated in primary monocyte-derived cells. J Virol 86, 13785–13789. 10.1128/JVI.01723-12.

51. Barbeau, B., and Mesnard, J.-M. (2015). Does chronic infection in retroviruses have a sense? Trends Microbiol. 23, 367–375. 10.1016/j.tim.2015.01.009.

52. Saayman, S., Ackley, A., Turner, A.-M.W., Famiglietti, M., Bosque, A., Clemson, M., Planelles, V., and Morris, K.V. (2014). An HIV-encoded antisense long noncoding RNA epigenetically regulates viral transcription. Mol Ther 22, 1164–1175. 10.1038/mt.2014.29.

53. Zapata, J.C., Campilongo, F., Barclay, R.A., DeMarino, C., Iglesias-Ussel, M.D., Kashanchi, F., and Romerio, F. (2017). The Human Immunodeficiency Virus 1 ASP RNA promotes viral latency by recruiting the Polycomb Repressor Complex 2 and promoting nucleosome assembly. Virology 506, 34–44. 10.1016/j.virol.2017.03.002.

54. Kobayashi-Ishihara, M., Terahara, K., Martinez, J.P., Yamagishi, M., Iwabuchi, R., Brander, C., Ato, M., Watanabe, T., Meyerhans, A., and Tsunetsugu-Yokota, Y. (2018). HIV LTR-Driven Antisense RNA by Itself Has Regulatory Function and May Curtail Virus Reactivation From Latency. Front. Microbiol. 9, 1066. 10.3389/fmicb.2018.01066.

55. Cassan, E., Arigon-Chifolleau, A.-M., Mesnard, J.-M., Gross, A., and Gascuel, O. (2016). Concomitant emergence of the antisense protein gene of HIV-1 and of the pandemic. Proc Natl Acad Sci U S A 113, 11537–11542. 10.1073/pnas.1605739113.

56. Li, R., Daneshvar, K., Ji, X., Pleet, M., Igbinosun, G., Iqbal, M.S., Kashanchi, F., Mullen, A.C., and Romerio, F. (2025). Suppression of HIV-1 transcription and latency reversal via ectopic expression of the viral antisense transcript AST. Sci Adv 11, eadu8014. 10.1126/sciadv.adu8014.

57. Chen, H.-C., Zorita, E., and Filion, G.J. (2018). Using Barcoded HIV Ensembles (B-HIVE) for Single Provirus Transcriptomics. Curr Protoc Mol Biol 122, e56. 10.1002/cpmb.56.

58. The R Project for Statistical Computing https://www.R-project.org/.

59. Lê, S., Josse, J., and Husson, F. (2008). FactoMineR: AnRPackage for Multivariate Analysis. J. Stat. Softw. 25. 10.18637/jss.v025.i01.

60. Kassambara, A., and Mundt, F. (2016). Factoextra: Extract and visualize the results of multivariate data analyses. (The R Foundation). 10.32614/cran.package.factoextra.

61. Csárdi, G., Nepusz, T., Müller, K., Horvát, S., Traag, V., Zanini, F., and Noom, D. (2025). igraph for R: R interface of the igraph library for graph theory and network analysis (Zenodo) 10.5281/ZENODO.14736815.

62. The pandas development team (2024). pandas-dev/pandas: Pandas (Zenodo) 10.5281/ZENODO.3509134.

63. Cock, P.J.A., Antao, T., Chang, J.T., Chapman, B.A., Cox, C.J., Dalke, A., Friedberg, I., Hamelryck, T., Kauff, F., Wilczynski, B., et al. (2009). Biopython: freely available Python tools for computational molecular biology and bioinformatics. Bioinformatics 25, 1422–1423. 10.1093/bioinformatics/btp163.

64. Li, H., Handsaker, B., Wysoker, A., Fennell, T., Ruan, J., Homer, N., Marth, G., Abecasis, G., Durbin, R., and 1000 Genome Project Data Processing Subgroup (2009). The Sequence Alignment/Map format and SAMtools. Bioinformatics 25, 2078–2079. 10.1093/bioinformatics/btp352.

65. Langmead, B., and Salzberg, S.L. (2012). Fast gapped-read alignment with Bowtie 2. Nat Methods 9, 357–359. 10.1038/nmeth.1923.

66. ENCODE Project Consortium, Moore, J.E., Purcaro, M.J., Pratt, H.E., Epstein, C.B., Shoresh, N., Adrian, J., Kawli, T., Davis, C.A., Dobin, A., et al. (2020). Expanded encyclopaedias of DNA elements in the human and mouse genomes. Nature 583, 699–710. 10.1038/s41586-020-2493-4.

67. Robinson, J.T., Thorvaldsdottir, H., Turner, D., and Mesirov, J.P. (2023). igv.js: an embeddable JavaScript implementation of the Integrative Genomics Viewer (IGV). Bioinformatics 39. 10.1093/bioinformatics/btac830.

68. Bar-Even, A., Paulsson, J., Maheshri, N., Carmi, M., O’Shea, E., Pilpel, Y., and Barkai, N. (2006). Noise in protein expression scales with natural protein abundance. Nat Genet 38, 636–643. 10.1038/ng1807.

69. Virtanen, P., Gommers, R., Oliphant, T.E., Haberland, M., Reddy, T., Cournapeau, D., Burovski, E., Peterson, P., Weckesser, W., Bright, J., et al. (2020). SciPy 1.0: fundamental algorithms for scientific computing in Python. Nat Methods 17, 261–272. 10.1038/s41592-019-0686-2.

70. Cox, C.D., McCollum, J.M., Allen, M.S., Dar, R.D., and Simpson, M.L. (2008). Using noise to probe and characterize gene circuits. Proc Natl Acad Sci U S A 105, 10809–10814. 10.1073/pnas.0804829105.

71. Mancarella, A., Procopio, F.A., Achsel, T., De Crignis, E., Foley, B.T., Corradin, G., Bagni, C., Pantaleo, G., and Graziosi, C. (2019). Detection of antisense protein (ASP) RNA transcripts in individuals infected with human immunodeficiency virus type 1 (HIV-1). J Gen Virol 100, 863–876. 10.1099/jgv.0.001244.

72. Weinberger, L.S., and Shenk, T. (2007). An HIV feedback resistor: auto-regulatory circuit deactivator and noise buffer. PLoS Biol 5, e9. 10.1371/journal.pbio.0050009.

73. Tantale, K., Garcia-Oliver, E., Robert, M.-C., L’Hostis, A., Yang, Y., Tsanov, N., Topno, R., Gostan, T., Kozulic-Pirher, A., Basu-Shrivastava, M., et al. (2021). Stochastic pausing at latent HIV-1 promoters generates transcriptional bursting. Nat Commun 12, 4503. 10.1038/s41467-021-24462-5.

74. Dey, S.S., Foley, J.E., Limsirichai, P., Schaffer, D.V., and Arkin, A.P. (2015). Orthogonal control of expression mean and variance by epigenetic features at different genomic loci. Mol Syst Biol 11, 806. 10.15252/msb.20145704.

75. Wang, G.P., Ciuffi, A., Leipzig, J., Berry, C.C., and Bushman, F.D. (2007). HIV integration site selection: analysis by massively parallel pyrosequencing reveals association with epigenetic modifications. Genome Res 17, 1186–1194. 10.1101/gr.6286907.

76. Einkauf, K.B., Lee, G.Q., Gao, C., Sharaf, R., Sun, X., Hua, S., Chen, S.M., Jiang, C., Lian, X., Chowdhury, F.Z., et al. (2019). Intact HIV-1 proviruses accumulate at distinct chromosomal positions during prolonged antiretroviral therapy. J Clin Invest 129, 988–998. 10.1172/JCI124291.

77. Jiang, C., Lian, X., Gao, C., Sun, X., Einkauf, K.B., Chevalier, J.M., Chen, S.M.Y., Hua, S., Rhee, B., Chang, K., et al. (2020). Distinct viral reservoirs in individuals with spontaneous control of HIV-1. Nature 585, 261–267. 10.1038/s41586-020-2651-8.

78. Lian, X., Gao, C., Sun, X., Jiang, C., Einkauf, K.B., Seiger, K.W., Chevalier, J.M., Yuki, Y., Martin, M., Hoh, R., et al. (2021). Signatures of immune selection in intact and defective proviruses distinguish HIV-1 elite controllers. Sci Transl Med 13, eabl4097. 10.1126/scitranslmed.abl4097.

79. Einkauf, K.B., Osborn, M.R., Gao, C., Sun, W., Sun, X., Lian, X., Parsons, E.M., Gladkov, G.T., Seiger, K.W., Blackmer, J.E., et al. (2022). Parallel analysis of transcription, integration, and sequence of single HIV-1 proviruses. Cell 185, 266–282.e15. 10.1016/j.cell.2021.12.011.

80. Lian, X., Seiger, K.W., Parsons, E.M., Gao, C., Sun, W., Gladkov, G.T., Roseto, I.C., Einkauf, K.B., Osborn, M.R., Chevalier, J.M., et al. (2023). Progressive transformation of the HIV-1 reservoir cell profile over two decades of antiviral therapy. Cell Host Microbe 31, 83–96.e5. 10.1016/j.chom.2022.12.002.

81. Machida, S., Depierre, D., Chen, H.-C., Thenin-Houssier, S., Petitjean, G., Doyen, C.M., Takaku, M., Cuvier, O., and Benkirane, M. (2020). Exploring histone loading on HIV DNA reveals a dynamic nucleosome positioning between unintegrated and integrated viral genome. Proc Natl Acad Sci U S A 117, 6822–6830. 10.1073/pnas.1913754117.

82. Thenin-Houssier, S., Machida, S., Jahan, C., Bonnet-Madin, L., Abbou, S., Chen, H.-C., Tesfaye, R., Cuvier, O., and Benkirane, M. (2023). POLE3 is a repressor of unintegrated HIV-1 DNA required for efficient virus integration and escape from innate immune sensing. Sci Adv 9, eadh3642. 10.1126/sciadv.adh3642.

83. Rando, O.J., and Chang, H.Y. (2009). Genome-wide views of chromatin structure. Annu Rev Biochem 78, 245–271. 10.1146/annurev.biochem.78.071107.134639.

84. Kaufman, P.D., and Rando, O.J. (2010). Chromatin as a potential carrier of heritable information. Curr Opin Cell Biol 22, 284–290. 10.1016/j.ceb.2010.02.002.

85. Lekstrom-Himes, J.A., Wang, K., Pesnicak, L., Krause, P.R., and Straus, S.E. (1998). The comparative biology of latent herpes simplex virus type 1 and type 2 infections: latency-associated transcript promoter activity and expression in vitro and in infected mice. J Neurovirol 4, 27–37. 10.3109/13550289809113479.

86. Kondo, K., Xu, J., and Mocarski, E.S. (1996). Human cytomegalovirus latent gene expression in granulocyte-macrophage progenitors in culture and in seropositive individuals. Proc Natl Acad Sci U S A 93, 11137–11142. 10.1073/pnas.93.20.11137.

87. Bangham, C.R.M., Miura, M., Kulkarni, A., and Matsuoka, M. (2019). Regulation of Latency in the Human T Cell Leukemia Virus, HTLV-1. Annu Rev Virol 6, 365–385. 10.1146/annurev-virology-092818-015501.

88. Capoferri, A.A., Sklutuis, R., Famuyiwa, T.O., Pathak, S., Li, R., Rausch, J.W., Luke, B.T., Hoh, R., Deeks, S.G., Mellors, J.W., et al. (2024). *In vivo*detection of HIV-1 antisense transcripts in untreated and ART-treated individuals. bioRxiv. 10.1101/2024.12.06.627170.

89. Klaver, B., and Berkhout, B. (1994). Comparison of 5’ and 3’ long terminal repeat promoter function in human immunodeficiency virus. J Virol 68, 3830–3840. 10.1128/JVI.68.6.3830-3840.1994.

90. Lefebvre, G., Desfarges, S., Uyttebroeck, F., Muñoz, M., Beerenwinkel, N., Rougemont, J., Telenti, A., and Ciuffi, A. (2011). Analysis of HIV-1 expression level and sense of transcription by high-throughput sequencing of the infected cell. J Virol 85, 6205–6211. 10.1128/JVI.00252-11.

91. Archin, N.M., Liberty, A.L., Kashuba, A.D., Choudhary, S.K., Kuruc, J.D., Crooks, A.M., Parker, D.C., Anderson, E.M., Kearney, M.F., Strain, M.C., et al. (2012). Administration of vorinostat disrupts HIV-1 latency in patients on antiretroviral therapy. Nature 487, 482–485. 10.1038/nature11286.

92. Deeks, S.G. (2012). HIV: Shock and kill. Nature 487, 439–440. 10.1038/487439a.

93. Abner, E., and Jordan, A. (2019). HIV “shock and kill” therapy: In need of revision. Antiviral Res 166, 19–34. 10.1016/j.antiviral.2019.03.008.

94. Ho, Y.-C., Shan, L., Hosmane, N.N., Wang, J., Laskey, S.B., Rosenbloom, D.I.S., Lai, J., Blankson, J.N., Siliciano, J.D., and Siliciano, R.F. (2013). Replication-competent noninduced proviruses in the latent reservoir increase barrier to HIV-1 cure. Cell 155, 540–551. 10.1016/j.cell.2013.09.020.

95. Weinberger, A.D., and Weinberger, L.S. (2013). Stochastic fate selection in HIV-infected patients. Cell 155, 497–499. 10.1016/j.cell.2013.09.039.

96. Dar, R.D., Hosmane, N.N., Arkin, M.R., Siliciano, R.F., and Weinberger, L.S. (2014). Screening for noise in gene expression identifies drug synergies. Science 344, 1392–1396. 10.1126/science.1250220.

97. Li, R., Caico, I., Xu, Z., Iqbal, M.S., and Romerio, F. (2023). Epigenetic Regulation of HIV-1 Sense and Antisense Transcription in Response to Latency-Reversing Agents. Noncoding RNA 9. 10.3390/ncrna9010005.

