## Supplementary files for "The influence of HIV sense and antisense transcripts on stochastic HIV transcription and reactivation"

**This PDF file includes:**

Supplementary Figures S1 to S9
Supplementary Tables S1 to S5

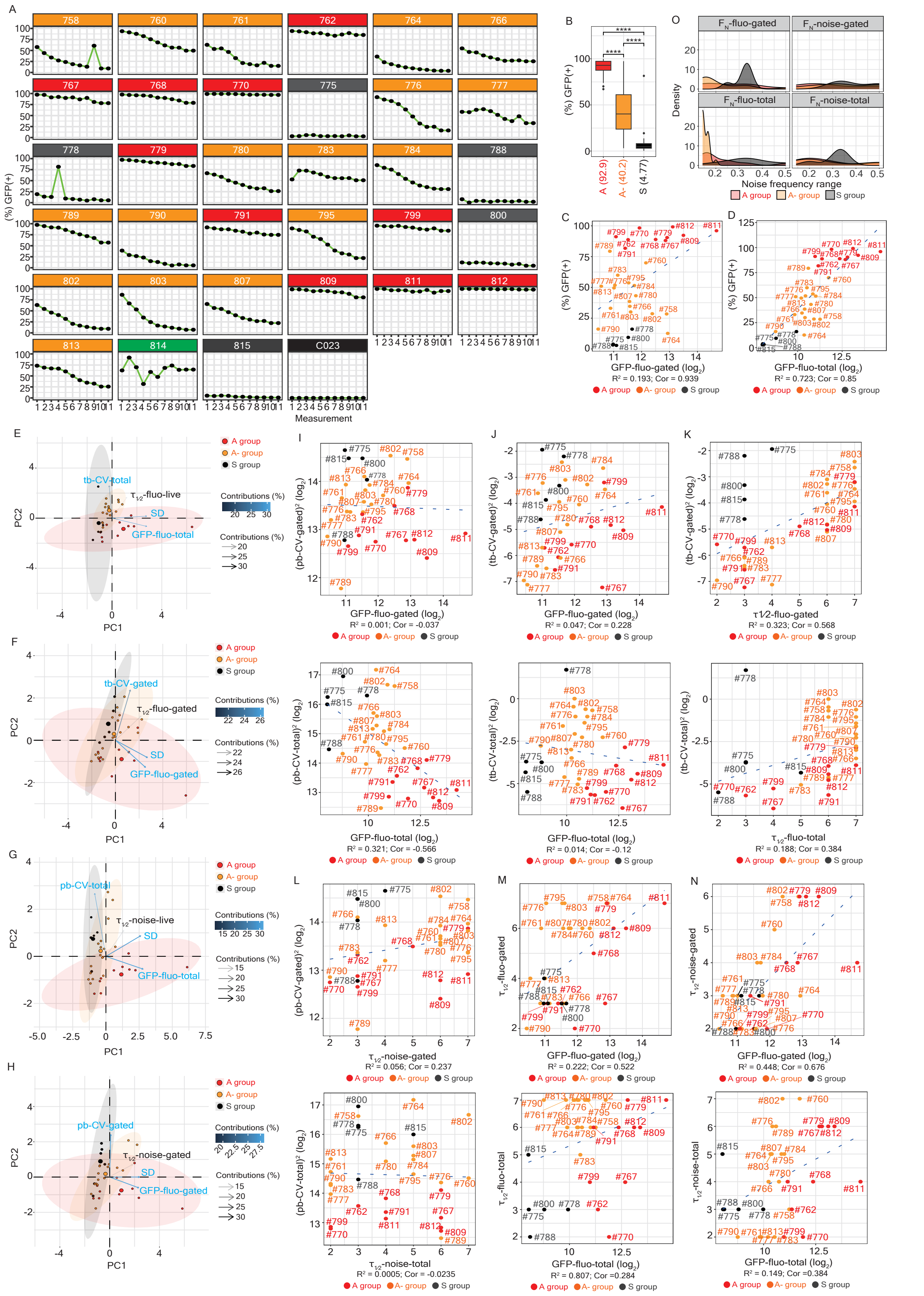

**Fig. S1. Characteristics of transcriptional phenotypes across sinpro clones in the A, A-, and S groups, related to Fig. 1.**
(**A**) Line plots representing the measure of (%) GFP(+) cells in 33 sinpro clones at 11 subsequent time points. Clone C023 is Jurkat T cells without HIV infections, as a negative control. (**B**) Boxplot representing the measures of (%) GFP(+) across clones in the A, A-, and S groups. Parentheses placed under the column

indicate the median of (%) GFP(+). ****$P$ < 0.0001. (**C**) Scatter plot representing the correlation between mean GFP expression (measured via GFP-fluo-gated) versus mean (%) GFP(+) across sinpro clones ($R^2$ = 0.193; correlation coefficient = 0.939). The y-axis is in logarithmic scale. Spots marked in red, orange, and grey represent sinpro clones in the A, A-, and S groups, respectively. (**D**) Scatter plot representing the correlation between mean GFP expression (measured via GFP-fluo-total) versus mean (%) GFP(+) across sinpro clones ($R^2$ = 0.723; correlation coefficient = 0.85). The y-axis is in logarithmic scale. Spots marked in red, orange, and grey represent sinpro clones in the A, A-, and S groups, respectively. (**E**, **F**, **G**, **H**) Principal component analysis (PCA) biplots illustrating the discrimination of sinpro clones in the A, A-, and S groups. Four different sets of parameters (i.e., variables) (detailed in the main text) are used to perform PCA, including (**E**) time-based (tb)-CV-total, GFP-fluo-total, and $\tau_{1/2}$-fluo-total, (**F**) tb-CV-gated, GFP-fluo-gated, and $\tau_{1/2}$-fluo-gated, (**G**) population-based (pb)-CV-total, GFP-fluo-total, and $\tau_{1/2}$-noise-total, and (**H**) pb-CV-gated, GFP-fluo-gated, and $\tau_{1/2}$-noise-gated. Arrows point the direction of variables. Color scale and color code shown in the arrows represent the contribution, a scaled version of the squared correlation between variables and component axes, of each variable. Individuals (sinpro clones) marked in red, orange, and grey represent sinpro clones in the A, A-, and S groups, respectively. Areas of separated individuals are enclosed in ellipses, following the same color codes as sinpro clones. (**I**) Scatter plot representing the correlation between mean GFP expression [measured via GFP-fluo-gated (top) or GFP-fluo-total (bottom)] versus population-based transcriptional noise [represented by squared pb-CV-gated (top) or squared pb-CV-total (bottom)] across sinpro clones. Both axes are in logarithmic scale. Spots marked in red, orange, and grey represent sinpro clones in the A, A-, and S groups, respectively. Top panel: $R^2$ = 0.001, correlation coefficient = -0.037; bottom panel: $R^2$ = 0.321; correlation coefficient = -0.566. (**J**) Scatter plot representing the correlation between mean GFP expression [measured via GFP-fluo-gated (top) or GFP-fluo-total (bottom)] versus time-based transcriptional noise [represented by squared tb-CVgated (top) or squared tb-CV-total (bottom)] across sinpro clones. Both axes are in logarithmic scale. Spots marked in red, orange, and grey represent sinpro clones in the A, A-, and S groups, respectively. Top panel: $R^2$ = 0.047, correlation coefficient = 0.228; bottom panel: $R^2$ = 0.014; correlation coefficient = -0.12. (**K**) Scatter plot representing the correlation between the autocorrelation time ($\tau_{1/2}$) [measured via $\tau_{1/2}$-fluo-gated (top) or $\tau_{1/2}$-fluo-total (bottom)] versus time-based transcriptional noise [represented by squared tb-CV-gated (top) or squared tb-CV-total (bottom)] across sinpro clones. The y-axis is in logarithmic scale. Spots marked in red, orange, and grey represent sinpro clones in the A, A-, and S groups, respectively. Top panel: $R^2$ = 0.323, correlation coefficient = 0.568; bottom panel: $R^2$ = 0.188; correlation coefficient = 0.384. (**L**) Scatter plot representing the correlation between the autocorrelation time ($\tau_{1/2}$) [measured via $\tau_{1/2}$-noise-gated (top) or $\tau_{1/2}$-noise-total (bottom)] versus population-based transcriptional noise [represented by squared pb-CV-gated (top) or squared pb-CV-total (bottom)] across sinpro clones. The y-axis is in logarithmic scale. Spots marked in red, orange, and grey represent sinpro clones in the A, A-, and S groups, respectively. Top panel: $R^2$ = 0.056, correlation coefficient = 0.237; bottom panel: $R^2$ = 0.0005; correlation coefficient = -0.0235. (**M**) Scatter plot representing the correlation between mean GFP expression [measured via GFP-fluo-gated (top) or GFP-fluo-total (bottom)] versus the autocorrelation time ($\tau_{1/2}$) [measured by $\tau_{1/2}$-fluo-gated (top) or $\tau_{1/2}$-fluo-total (bottom)] across sinpro clones. The x-axis is in logarithmic scale. Spots marked in red, orange, and grey represent sinpro clones in the A, A-, and S groups, respectively. Top panel: $R^2$ = 0.222, correlation coefficient = 0.522; bottom panel: $R^2$ = 0.807; correlation coefficient = 0.284. (**N**) Scatter plot representing the correlation between mean GFP expression [measured by GFP-fluo-gated (top) or GFP-fluo-total (bottom)] versus the autocorrelation time ($\tau_{1/2}$) [measured by $\tau_{1/2}$-noise-gated (top) or $\tau_{1/2}$-noise-total (bottom)] across sinpro clones. The x-axis is in logarithmic scale. Spots marked in red, orange, and grey represent sinpro clones in the A, A-, and S groups, respectively. Top panel: $R^2$ = 0.448, correlation coefficient = 0.676; bottom panel: $R^2$ = 0.149; correlation coefficient = 0.384. (**O**) Histogram representing the range of noise frequency. Facets separate the frequency calculated based on $F_N$-fluo-gated (top panel on the left-hand side), $F_N$-fluo-total (bottom panel on the left-hand side), $F_N$-noise-gated (top panel on the right-hand side), and $F_N$-noise-total (bottom panel on the right-hand side).

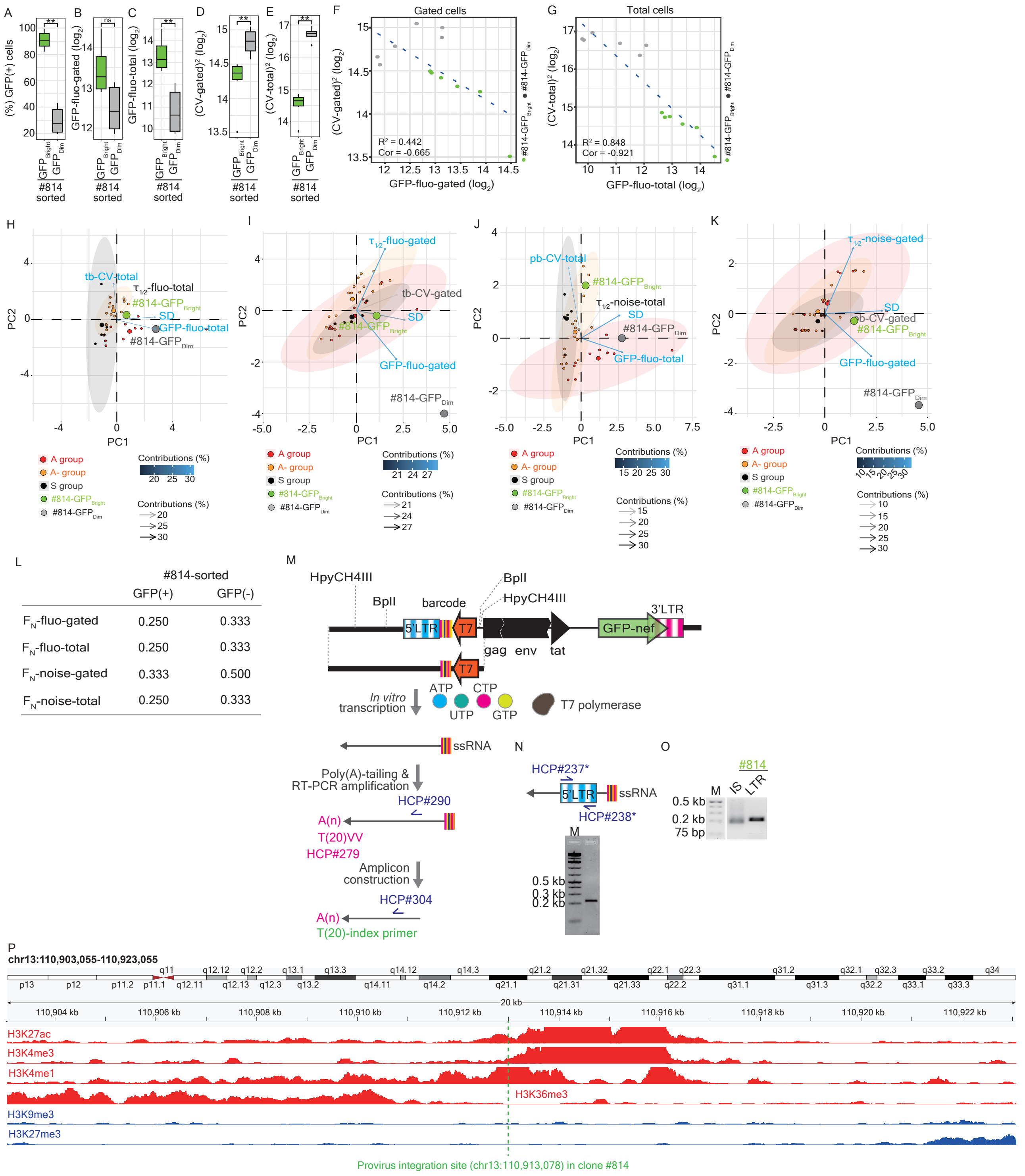


**Fig. S2. Characteristics of transcriptional phenotypes in clone #814 (the F group), related to Fig. 2.** (**A**) Boxplot representing (%) GFP(+) cells between FACS-sorted #814-GFP$_{Bright}$ and GFP$_{Dim}$ subpopulations. Boxes in green and grey represent FACS-sorted #814-GFP$_{Bright}$ and GFP$_{Dim}$ subpopulations, respectively. ***P* < 0.01. (**B**) Boxplot representing mean GFP expression (measured via GFP-fluo-gated in logarithmic scale) between FACS-sorted #814-GFP$_{Bright}$ and GFP$_{Dim}$ subpopulations. Boxes in green and grey represent FACS-sorted #814-GFP$_{Bright}$ and GFP$_{Dim}$ subpopulations, respectively. (**C**) Boxplot representing mean GFP expression (measured via GFP-fluo-total in logarithmic scale) between FACS-sorted #814-GFP$_{Bright}$ and GFP$_{Dim}$ subpopulations. Boxes in green and grey represent FACS-sorted #814-GFP$_{Bright}$ and GFP$_{Dim}$ subpopulations, respectively. ***P* < 0.01. (**D**) Boxplot representing transcriptional noise (measured via squared CV-gated in logarithmic scale) between FACS-sorted #814-GFP$_{Bright}$ and GFP$_{Dim}$ subpopulations. Boxes in green and grey represent FACS-sorted #814-GFP$_{Bright}$ and GFP$_{Dim}$ subpopulations, respectively. ***P* < 0.01. (**E**) Boxplot representing transcriptional noise (measured via squared CV-total in logarithmic scale) between FACS-sorted #814-GFP$_{Bright}$ and GFP$_{Dim}$ subpopulations. Boxes in green and grey represent FACS-sorted #814-GFP$_{Bright}$ and GFP$_{Dim}$ subpopulations, respectively. ***P* < 0.01. (**F**) Scatter plot representing the correlation between mean GFP expression (measured via GFP-fluo-gated) versus transcriptional noise (represented by squared CV-gated) across sinpro clones. Both axes are in logarithmic scale. Spots marked in green and grey represent FACS-sorted #814-$GFP_{Bright}$ and $GFP_{Dim}$ subpopulations, respectively. $R^2$ = 0.442, correlation coefficient = -0.665. (**G**) Scatter plot representing the correlation between mean GFP expression (measured by GFP-fluo-total) versus transcriptional noise (represented by squared CV-total) across sinpro clones. Both axes are in logarithmic scale. Spots marked in green and grey represent FACS-sorted #814-$GFP_{Bright}$ and $GFP_{Dim}$ subpopulations, respectively. $R^2$ = 0.848, correlation coefficient = -0.921. (**H**, **I**, **J**, **K**) Principal component analysis (PCA) biplots illustrating the discrimination of sinpro clones in the A, A-, and S groups and FACS-sorted #814-$GFP_{Bright}$ and $GFP_{Dim}$ subpopulations. Four different sets of parameters (i.e., variables) (detailed in the main text) are used to perform PCA, including (**H**) time-based (tb)-CV-total, GFP-fluo-total, and $\tau_{1/2}$-fluo-total, (**I**) tb-CV-gated, GFP-fluo-gated, and $\tau_{1/2}$-fluo-gated, (**J**) population-based (pb)-CV-total, GFP-fluo-total, and $\tau_{1/2}$-noise-total, and (**K**) pb-CV-gated, GFP-fluo-gated, and $\tau_{1/2}$-noise-gated. Arrows point the direction of variables. Color scale and color code shown in the arrows represent the contribution, a scaled version of the squared correlation between variables and component axes, of each variable. Individuals (sinpro clones) marked in red, orange, dark grey, green, and light grey represent sinpro clones in the A, A-, and S groups, and FACS-sorted #814-$GFP_{Bright}$ and $GFP_{Dim}$ subpopulations, respectively. Areas of separated individuals are enclosed in ellipses, following the same color codes as sinpro clones. (**L**) Table demonstrating the calculation of the noise frequency range ($F_N$) corresponding to four different sets of parameters between FACS-sorted #814-$GFP_{Bright}$ and $GFP_{Dim}$ subpopulations. (**M**) Schematic representation of the experimental workflow of lentiviral integration site sequencing (LIS-seq) (detailed in the **Material and methods**). Genomic DNA isolated from sinpro clones was digested with restriction enzymes, HpyCH4III and BplI, followed by *in vitro* transcription, and RT-PCR amplification with primers embedded with Illumina adaptor sequences. Amplicons are sequenced as 50 bp single reads on a NovaSeq 6000 sequencer (Illumina). (**N**) Verification of production of single-strand RNA (ssRNA) resulting from *in vitro* transcription. ssRNA was used as the template for the performance of RT-PCR amplification, yielding a correct 220 bp product. (**O**) Expected result from the sequencing amplicon using LIS-seq. 2.0% (wt/vol) agarose gel displaying a faint PCR smear corresponding to fragments in different lengths harboring the integration site of the provirus in clone #814. (**P**) Integrative Genomics Viewer (IGV) illustrating epigenetic landscape ranging from the locus at 110,903,055 base pairs (bp) to 110,923,055 bp at the chromosome 13 (chr13). The provirus integration site that mapped at the locus 110,913,078 at chr13 is marked with a green dotted line. ChIP-seq signals (top to bottom) loaded in the browser are H3K27ac (ENCFF701MUZ), H3K4me3 (ENCFF067KWB), H3K4me1 (ENCFF704DNO), H3K36me3 (ENCFF536HDR), H3K9me3 (ENCFF353AHL), and H3K27me3 (ENCFF153LTQ) and are downloaded from the ENCODE Project Consortium. Signal coverages colored in red and blue represent active and repressive histone marks.

A

| Clone | Group | Chr | Position | Gene | Number of mapped reads to the locus |
|---|---|---|---|---|---|
| 760 | A- | chr3 | 17,402,193 | TBC1D5 | 1,621 |
| 761 | A- | chr5 | 78,769,319 | LHFPL2 | 30,206 |
| 762 | A | chr3 | 98,514,271 | CLDND1 | 22,531 |
| 770 | A | chr5 | 123,554,312 | CSNK1G3 | 11,157 |
| 776 | A- | chr1 | 183,515,691 | SMG7 | 8,135 |
| 778 | S | chr14 | 96,325,410 | ATG2B | 73,020 |
| 788 | S | chr1 | 158,447,409 | Intergenic region | 101,159 |
| 800 | S | chr3 | 184,886,958 | VPS8 | 1,107 |
| 802 | A- | chr6 | 14,256,877 | Intergenic region | 87,037 |
| 803 | A- | chr16 | 133,573 | NPRL3 | 47,550 |
| 811 | A | chr11 | 64,231,874 | DNAJC4 | 60,079 |
| 812 | A | chr11 | 65,426,090 | NEAT1 | 45,421 |
| 813 | A- | chr2 | 17,685,226 | SMC6 | 12,559 |
| 814 | F | chr13 | 110,913,078 | ANKRD10 | 221,470 |

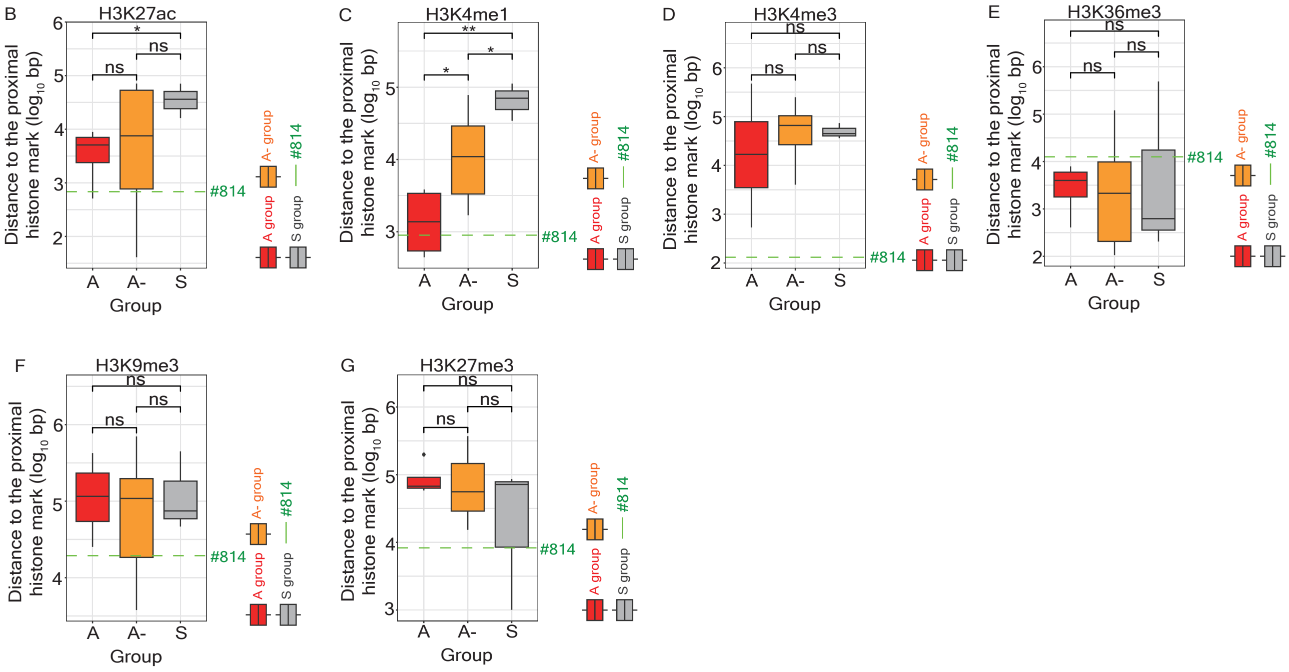


**Fig. S3. Characteristics of provirus integration sites associated with histone modifications in the A, A-, and S groups, related to Fig. 2.**

(**A**) A table summarizing the characteristics of the provirus integration site, including the chromosome, the position of the nucleotide, at which the provirus integrates, the corresponding gene symbol, and the number of the filtered reads mapped to the indicated locus in sinpro clone #760 (the A- group), #761 (the A- group), #762 (the A group), #770 (the A group), #776 (the A- group), #778 (the S group), #788 (the S group), #800 (the S group), #802 (the A- group), #803 (the A- group), #811 (the A group), #812 (the A group), #813 (the A- group), and #814 (the F group). (**B**, **C**, **D**, **E**, **F**, **G**) Boxplots representing the distance to the closest histone marks, including H3K27ac (**B**), H3K4me1 (**C**), H3K4me3 (**D**), H3K36me3 (**E**), H3K9me3 (**F**), and H3K27me3 (**G**) of proviruses retrieved in clones assigned to the A (#762, #770, #811, and #812), A- (#760, #761, #776, #802, #803, and #813), and S (#778, #788, and #800) groups. Distance is plotted in logarithmic scale of base pairs (bp) on the y-axis. Boxes marked in red, orange, and grey represent sinpro clones in the A, A-, and S groups, respectively. The distance to the closest histone marks of the provirus integration site in clone #814 is highlighted with the green dotted line in each boxplot. Significance levels are denoted as follows: *$P < 0.05$; **$P < 0.01$.

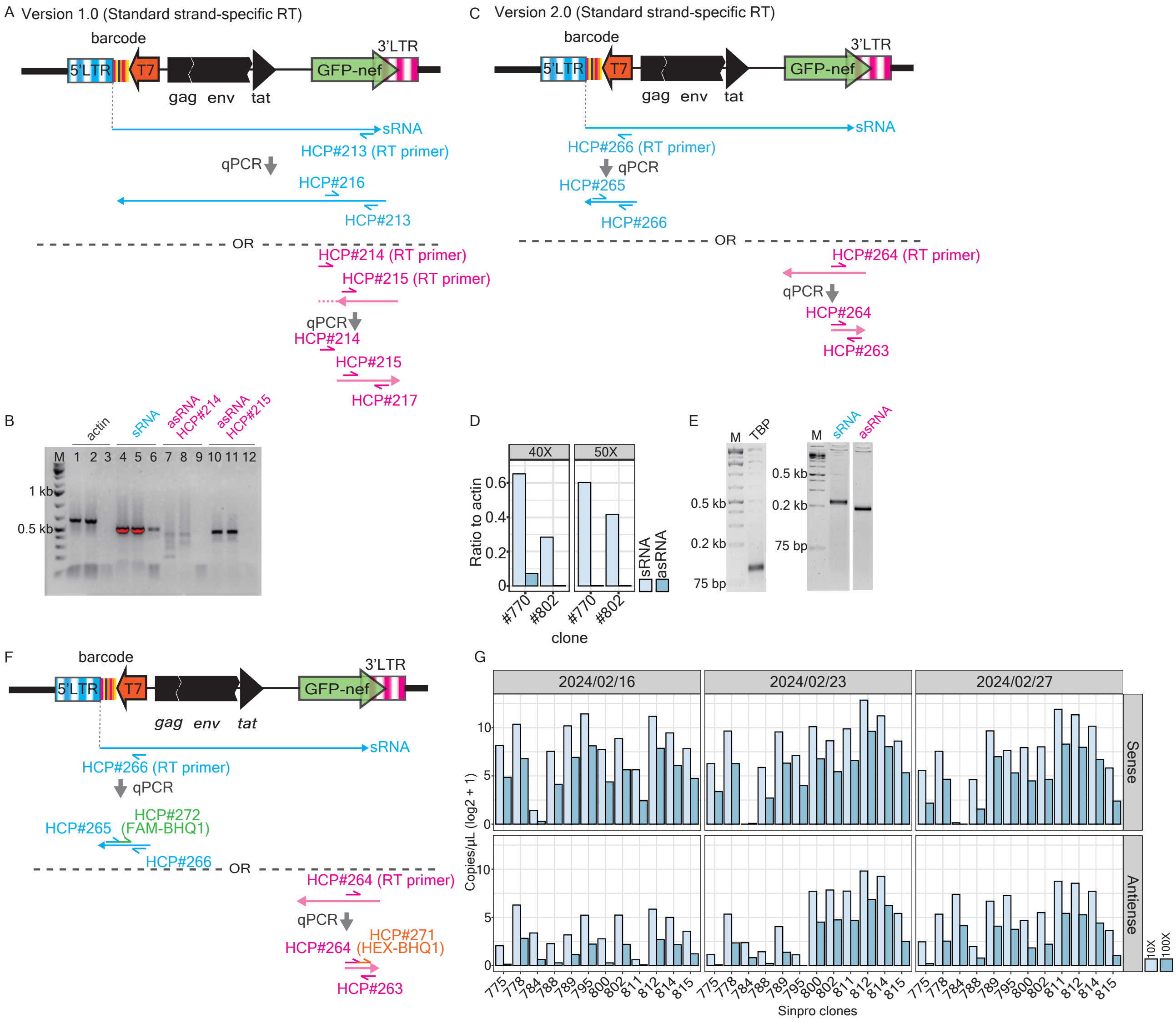


**Fig. S4. Quantitative measurement of HIV sense and antisense transcripts, related to Fig. 3.**
(**A**) Schematic representation of the experimental workflow of strand-specific reverse transcription (RT) followed by quantitative polymerase chain reaction (qPCR) version 1.0. RT primers specific to sense RNAs (sRNAs, HCP#213) and antisense RNAs (asRNAs, HCP#214 or HCP#215) are applied to independent amplification of cDNA reverse-transcribed from sense and antisense transcripts, respectively. Arrows indicate the direction of RT and DNA synthesis driven by primers. Arrows in azure and primers written in azure refer to RT-qPCR amplification based on sRNAs; arrows in magenta and primers written in magenta refer to RT-qPCR amplification based on asRNAs. (**B**) 2.0% (wt/vol) agarose gel displaying a strand-specific RT-qPCR products. A specific 509 bp strand-specific RT-qPCR product was yielded from sRNAs (lanes 4 and 5) and a specific 453 bp strand-specific RT-qPCR product was yielded from asRNAs (lanes 10 and 11). No specific RT-qPCR product was detectable using the RT primer HCP#214 (lanes 7 and 8). The human actin gene is used as a reference (lanes 1 and 2). Lanes 3, 6, 9, and 12: water control. (**C**) Schematic representation of the experimental workflow of strand-specific RT-qPCR version 2.0. RT primers specific to sense RNAs (sRNAs, HCP#266) and antisense RNAs (asRNAs, HCP#264) are applied to independent amplification of cDNA reverse-transcribed from sense and antisense transcripts, respectively. Arrows indicate the direction of RT and DNA synthesis driven by primers. Arrows in azure and primers written in azure refer to RT-qPCR amplification based on sRNAs; arrows in magenta and primers written in magenta refer to RT-qPCR amplification based on asRNAs. (**D**) Bar charts representing the feasibility of strand-specific RT-qPCR version 2.0 mentioned in panel **C**. RNA isolated from clones #770 and #802 were subjected to strand-specific RT-qPCR with the amplification with either 40 (the facet on the left-hand side) or 50 cycles (the facet on the right-hand side), with actin as a reference. Bars in light blue and azure represent the amplification using sRNAs and asRNAs as templates, respectively. (**E**) 2.0% (wt/vol) agarose gel displaying a strand-specific PCR-qPCR product. A specific 230 bp strand-specific RT-qPCR product was yielded from sRNAs (the gel on the right-hand side) and a specific 194 bp strand-specific RT-qPCR product was yielded from asRNAs (the gel on the right-hand side). The human TATA-box binding protein (TBP) gene is used as a

reference (113 bp, the gel on the left-hand side). (**F**) Schematic representation of the experimental workflow of strand-specific RT followed by digital PCR (dPCR). The setting of strand-specific RT and PCR primers is identical to that used in strand-specific RT-qPCR version 2.0 (**C**). Probes, FAM-BHQ1 (HCP#272) and HEX-BHQ1 (HCP#271), are utilized to quantify sense and antisense transcripts, respectively. (**G**) Bar charts representing the quantification of sense (top rows) and antisense (bottom rows) transcripts across sinpro clones at three subsequent time points. Bars in light blue and azure represent the tenfold (10X) and hundredfold (100X) dilution of cDNA for the performance of dPCR.

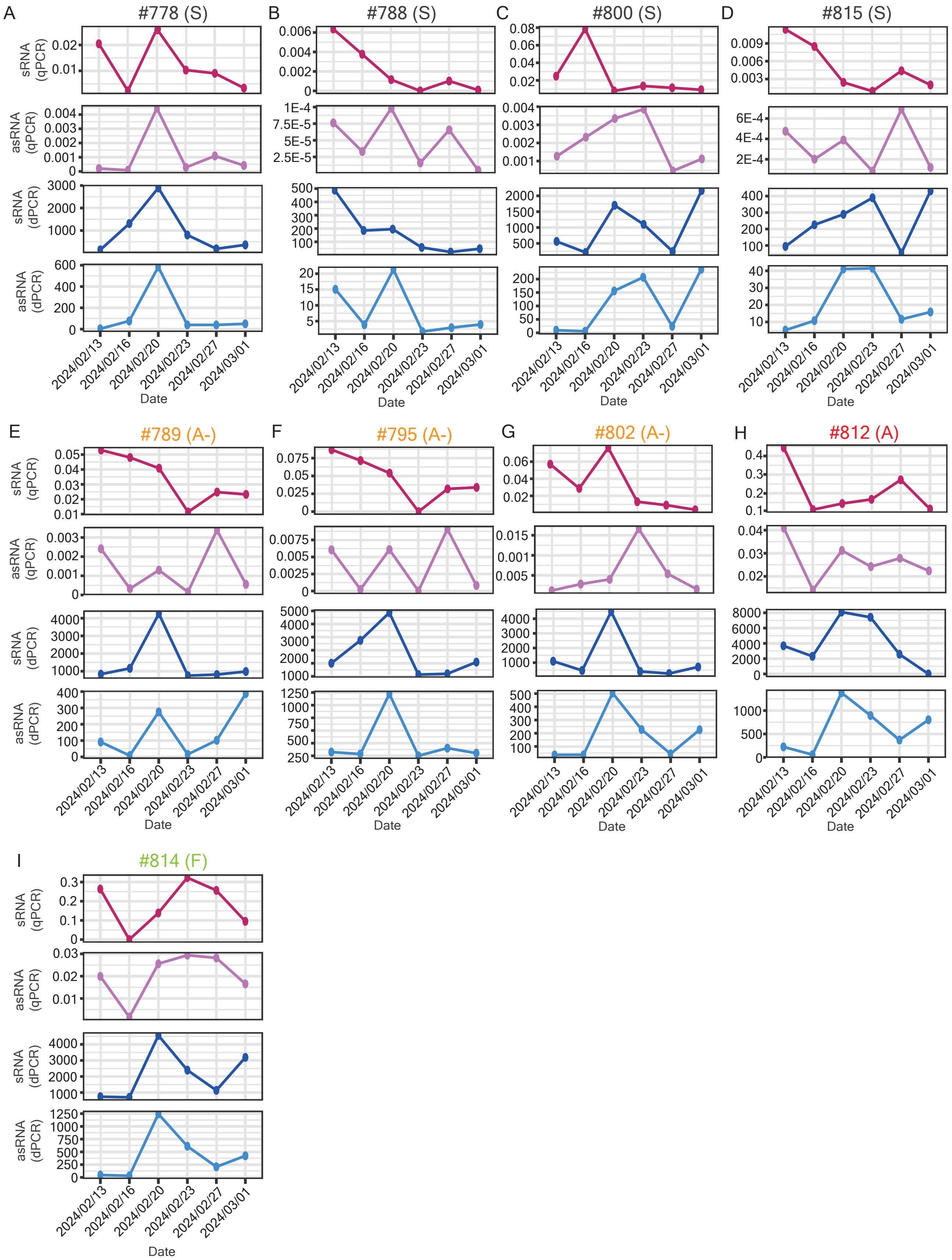


**Fig. S5. Tracking provirus sense and antisense RNA transcription in nine sinpro clones at six time points, related to Fig. 3.**
(**A-I**) Line plots representing the measures of the abundance of provirus sense (the first and third rows from top) and antisense (the second and fourth rows from top) transcripts measured using strand-specific RT qPCR (the first two rows from top) or dPCR (the first two rows from bottom) at six time points in sinpro clones in the S (**A**, #778; **B**, #788; **C**, #800; **D**, #815), A- (**E**, #789; **F**, #795; **G**, #802), A (**H**, #812) and F (**I**, #814) group.

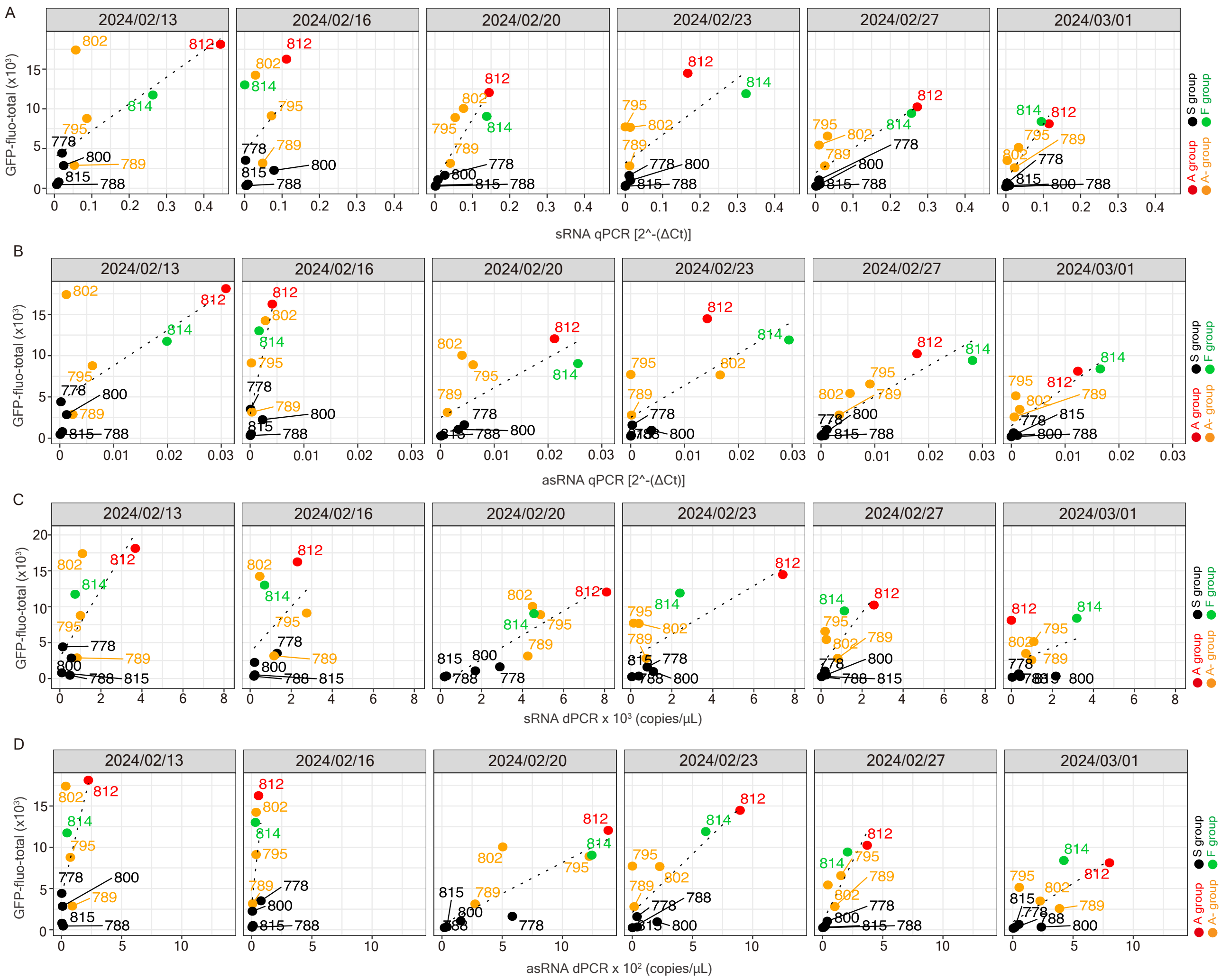

**Fig. S6. Consistency in expression measured at the protein and the RNA level, related to Fig. 3.**
(**A**) Scatter plot representing a positive correlation between the abundance of sRNAs measured by strand-specific qPCR and mean GFP expression (measured via GFP-fluo-total) at six subsequent time points. Spots marked in red, orange, black, and green indicate sinpro clones in the A, A-, S, and F groups, respectively. (**B**) Scatter plot representing a positive correlation between the abundance of asRNAs measured by strand-specific qPCR and mean GFP expression (measured via GFP-fluo-total) at six subsequent time points. Spots marked in red, orange, black, and green indicate sinpro clones in the A, A-, S, and F groups, respectively. (**C**) Scatter plot representing a positive correlation between the abundance of sRNAs measured by strand-specific dPCR and mean GFP expression (measured via GFP-fluo-total) at six subsequent time points. Spots marked in red, orange, black, and green indicate sinpro clones in the A, A-, S, and F groups, respectively. (**D**) Scatter plot representing a positive correlation between the abundance of asRNAs measured by strand-specific dPCR and mean GFP expression (measured via GFP-fluo-total) at six subsequent time points. Spots marked in red, orange, black, and green indicate sinpro clones in the A, A-, S, and F groups, respectively.

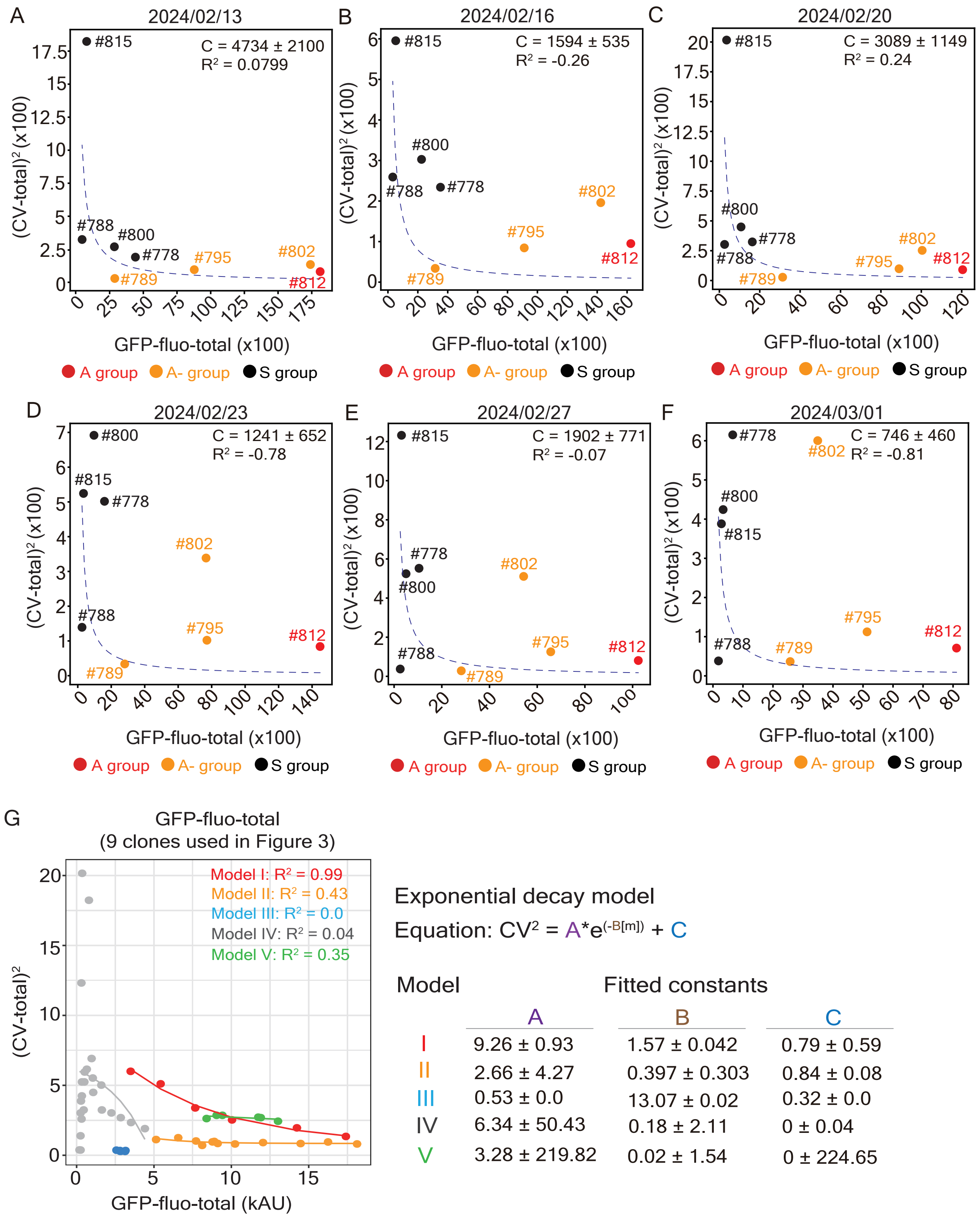


**Fig. S7. The performance of mathematical modeling across sinpro clones, related to Fig. 4.** (**A-F**) Plots of mean GFP expression (measured via GFP-fluo-total) versus transcriptional noise (measured via squared CV-total) across eight sinpro clones in A (#812), A- (#789, #795, and #802), and S (#778, #788, #800, and #815) groups at six subsequent time points based on the reciprocal curve model (Equation 1) mentioned in the main text. Dotted blue lines represent the fitting curves corresponding to each time point: (**A**) 2024/02/13, $R^2$ = 0.0799; (**B**) 2024/02/16, $R^2$ = -0.26; (**C**) 2024/02/20, $R^2$ = 0.24; (**D**) 2024/02/23, $R^2$ = -0.78; (**E**) 2024/02/13, $R^2$ = -0.07; (**F**) 2024/02/13, $R^2$ = -0.81. The value of the proportionality factor ($C$) corresponding to each time point is provided alongside the $R^2$ value in each panel. Spots marked in red, orange, and black represent sinpro clones in the A, A-, and S groups. (**G**) Plots of mean GFP expression (measured via GFP-fluo-total) versus transcriptional noise (measured via squared CV-total) across the same eight sinpro clones as those used in panels **A**-**F** and clone #814 (F group) at six subsequent time points based on the exponential decay model (Equation 2) mentioned in the main text. The $R^2$ values corresponding to individual models are labeled in the panel. The A, B, and C constants in the formula of exponential decay are summarized on the right-hand side of panel **G**. Spots and lines in red, orange, azure, grey, and green represent clones and curve fitting in models **I**, **II**, **III**, IV, and V, respectively.

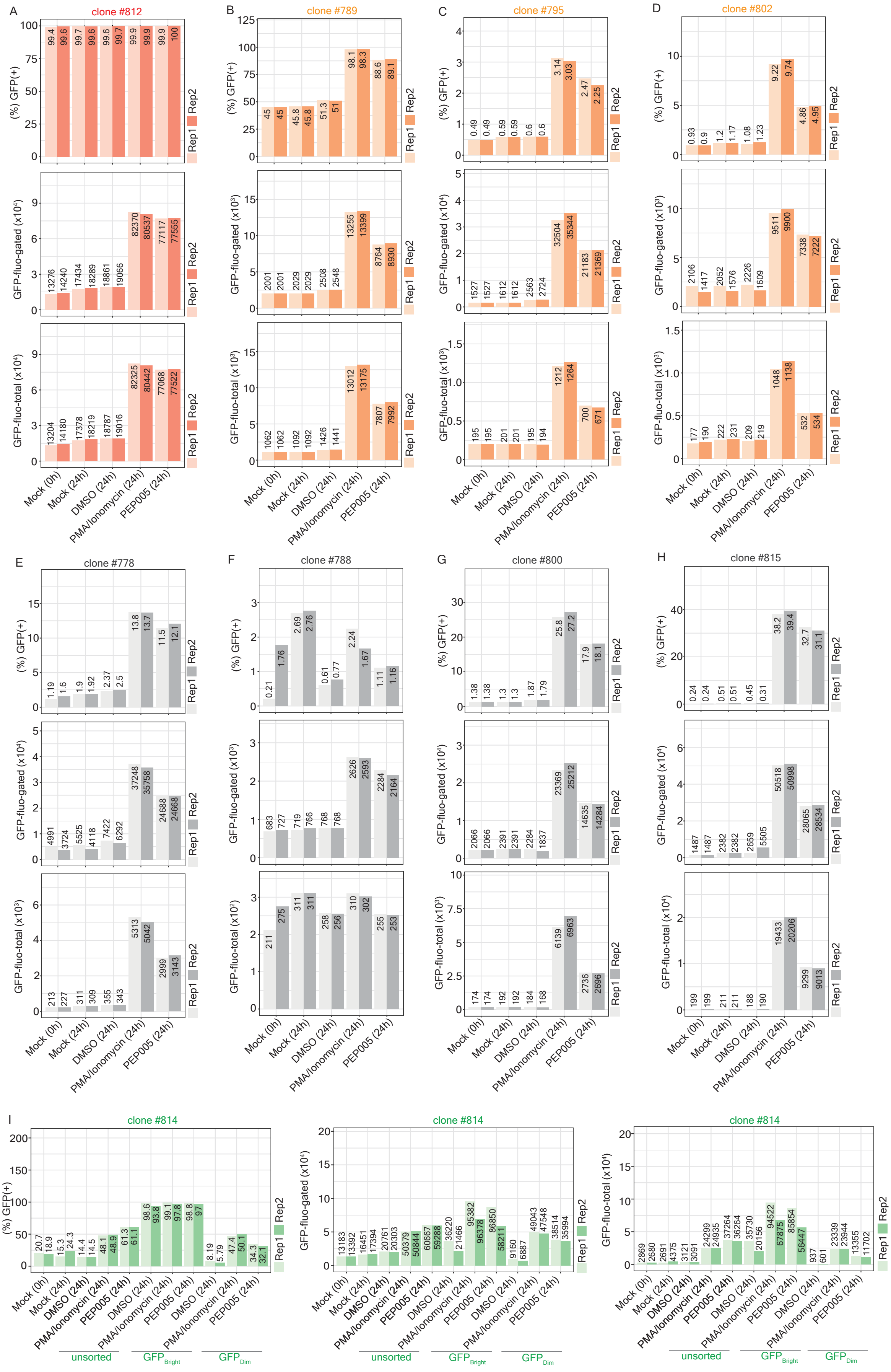

**Fig. S8. FACS measurement of reactivation of proviruses in sinpro clones treated with drugs, related to Fig. 5.**

(**A-H**) Bar charts representing the enrichment of reactivation of proviruses subjected to PMA/Ionomycon or PEP005 across eight sinpro clones, including (**A**) clone #812 (the A group), (**B**) clone #789 (the A- group),

(**C**) clone #795 (the A- group), (**D**) clone #802 (the A- group), (**E**) clone #778 (the S group), (**F**) clone #788 (the S group), (**G**) clone #800 (the S group), and (**H**) clone #815 (the S group). FACS measures of (%) GFP(+) cells (top row), GFP-fluo-gated (middle row), and GFP-fluo-total (bottom row) are shown; measures are labeled aside each bar. Two replicates (Rep1 and Rep2) were carried out in each experiment. Mock (0 h): drug-free control; measurement before the drug treatment. Mock (24 h): drug-free control; measurement conducted after 24 hr incubation time. DMSO (24 h): clones subjected to DMSO; measurement conducted 24 hr post treatment. PMA/Ionomycin (24 h): clones subjected to PMA/Ionomycin; measurement conducted 24 hr post treatment. PEP005 (24 h): clones subjected to PEP005; measurement conducted 24 hr post treatment. (**I**) Bar charts representing the enrichment of reactivation of proviruses subjected to PMA/ Ionomycon or PEP005 in unsorted clone #814 and FACS-sorted clone #814-$GFP_{Bright}$ and $GFP_{Dim}$ subpopulations. FACS measures of (%) GFP(+) cells (left), GFP-fluo-gated (middle), and GFP-fluo-total (right) are shown; measures are labeled aside each bar. Two replicates (Rep1 and Rep2) were carried out in each experiment. Mock (0 h): drug-free control; measurement before the drug treatment. Mock (24 h): drug-free control; measurement conducted after 24 hr incubation time. DMSO (24 h): clones subjected to DMSO; measurement conducted 24 hr post treatment. PMA/Ionomycin (24 h): clones subjected to PMA/Ionomycin; measurement conducted 24 hr post treatment. PEP005 (24 h): clones subjected to PEP005; measurement conducted 24 hr post treatment.

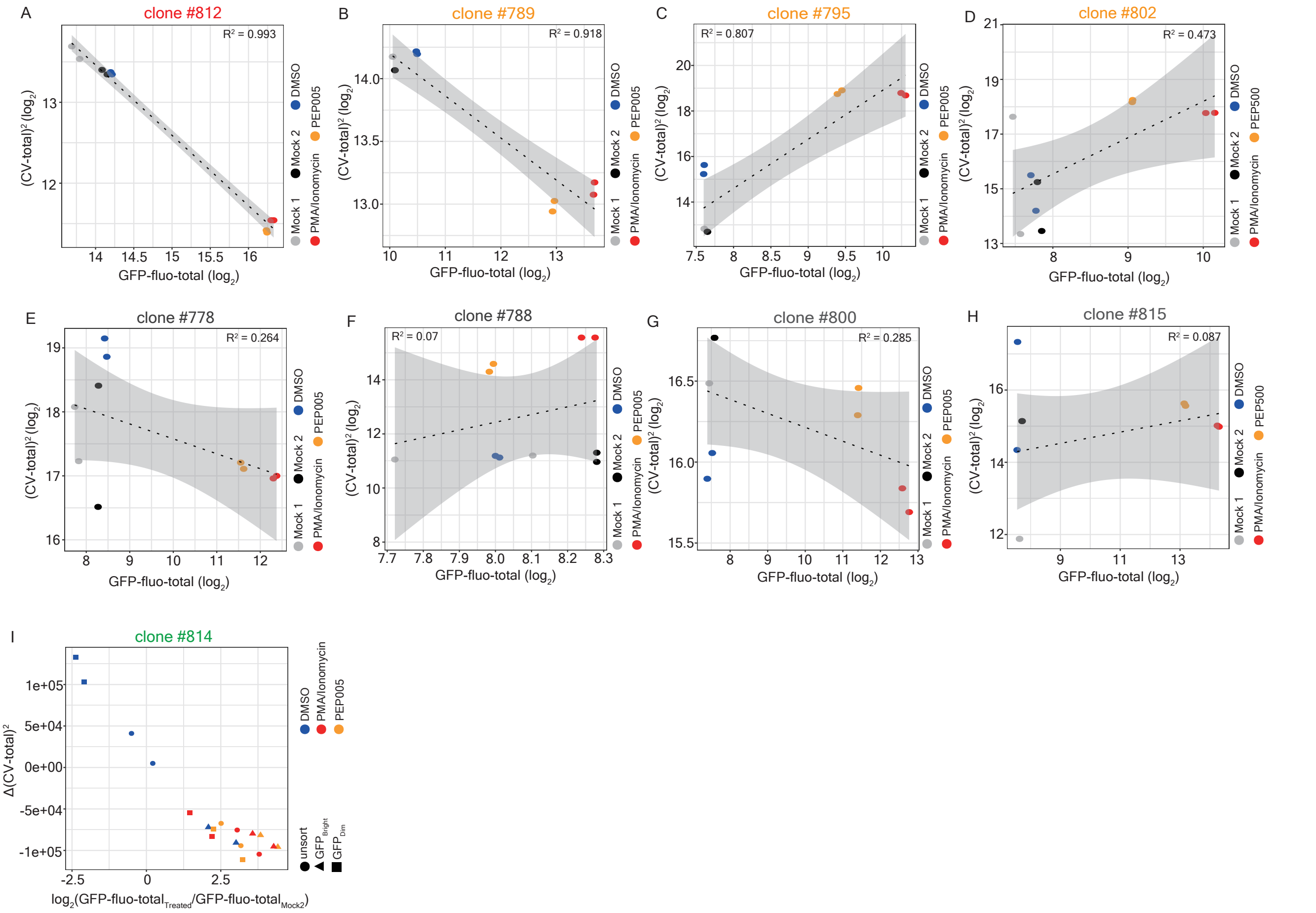


**Fig. S9. Different responses of the proviruses present in distinct groups toward PMA/Ionomycin or PEP005 treatments, related to Fig. 5.**

(**A**-**H**) Scatter plots representing a correlation between mean GFP expression (measured via GFP-fluo-total) and transcriptional noise (measured via squared CV-total) across eight sinpro clones, including (**A**) clone #812 (the A group, $R^2 = 0.993$), (**B**) clone #789 (the A- group, $R^2 = 0.918$), (**C**) clone #795 (the A- group, $R^2 = 0.807$), (**D**) clone #802 (the A- group, $R^2 = 0.473$), (**E**) clone #778 (the S group, $R^2 = 0.264$), (**F**) clone #788 (the S group, $R^2 = 0.07$), (**G**) clone #800 (the S group, $R^2 = 0.285$), and (**H**) clone #815 (the S group, $R^2 = 0.087$) in the presence of different drugs. Both axes are in logarithmic scale. Mock 1 (grey spots): drug-free control; measurement before the drug treatment. Mock 2 (black spots): drug-free control; measurement conducted after 24 hr incubation time. DMSO (blue spots): clones subjected to DMSO; measurement conducted 24 hr post treatment. PMA/Ionomycin (red spots): clones subjected to PMA/Ionomycin; measurement conducted 24 hr post treatment. PEP005 (orange spots): clones subjected to PEP005; measurement conducted 24 hr post treatment. (**I**) Scatter plot representing a correlation between the change in mean GFP expression and in transcriptional noise in unsorted clone #814 (circle) and FACS-sorted #814-$\text{GFP}_{\text{Bright}}$ (triangle) and $\text{GFP}_{\text{Dim}}$ (square) subpopulations in the presence of different drugs. The change in mean GFP expression is calculated with the formula $\text{GFP-fluo-total}_{\text{Treated}}/\text{GFP-fluo-total}_{\text{Mock2}}$; the change in transcriptional noise is calculated with the formula $(\text{CV-total}_{\text{Treated}})^2-(\text{CV-total}_{\text{Mock2}})^2$. Spots marked in blue, red, and orange represent clones subjected to DMSO, PMA/Ionomycin, and PEP005, respectively.

**Table S1.** FACS measurements conducted in 2023 across 33 sinpro clones in 11 subsequent time points, related to Fig. 1 and 4.

Excel file containing additional data too large to fit in a PDF.

**Table S2.** Measures of four sets of parameters used to construct noise space, related to Fig. 1 and 2.

| Clone | GFP-fluo-gated ($\log_2$) | (tb-CV-gated)$^2$ | (pb-CV-gated)$^2$ | $\tau_{1/2}$-noise-gated | $\tau_{1/2}$-fluo-gated | GFP-fluo-total ($\log_2$) | (tb-CV-total)$^2$ | (pb-CV-total)$^2$ | $\tau_{1/2}$-noise-total | $\tau_{1/2}$-fluo-total | Group |
|---|---|---|---|---|---|---|---|---|---|---|---|
| 758 | 12.8805321629162225 | 0.14679690260283432 | 2.248333641975309 | 6.0 | 7.0 | 11.23066262505995 | 0.5187421671944319 | 9.744802777777778 | 3.0 | 6.0 | A- |
| 760 | 12.1414774254105582 | 0.04950844274940569 | 1.3346667438271607 | 5.0 | 6.0 | 11.730806217392418 | 0.17337169917489223 | 2.3091335069444443 | 7.0 | 7.0 | A- |
| 761 | 10.874874648121818 | 0.0697795253759948 | 1.28457037345679 | 3.0 | 6.0 | 9.81431542378781 | 0.28401209395970334 | 2.6900555748456796 | 2.0 | 7.0 | A- |
| 762 | 11.502493434228947 | 0.009225830355324833 | 1.0218963456790124 | 2.0 | 3.0 | 11.352855732531157 | 0.012970850775239618 | 1.2179574845679013 | 3.0 | 3.0 | A |
| 764 | 12.93599629184421 | 0.0821488692863414 | 1.6132527970679014 | 3.0 | 7.0 | 10.327739751926764 | 0.6268993296656873 | 15.015625 | 5.0 | 6.0 | A- |
| 766 | 11.57127203373745 | 0.009925742300087519 | 1.7711173611111117 | 2.0 | 3.0 | 10.235781566151553 | 0.0774569376048029 | 5.363727334104938 | 4.0 | 7.0 | A- |
| 767 | 12.85264058337002 | 0.0043237977762904355 | 0.7005225007716049 | 4.0 | 3.0 | 12.680484675714043 | 0.008804442326708403 | 0.910725602044753 | 6.0 | 4.0 | A |
| 768 | 12.509548225452248 | 0.027871329432927153 | 1.1499583526234567 | 4.0 | 5.0 | 12.375642842222295 | 0.05230688036226133 | 1.4210626736111112 | 4.0 | 6.0 | A |
| 770 | 11.90917951646238 | 0.012290725469093922 | 0.6840668402777776 | 2.0 | 2.0 | 11.887749489771702 | 0.013466693550640427 | 0.7071641992669754 | 2.0 | 2.0 | A |
| 775 | 11.007056475681557 | 0.20328989955423943 | 2.6394063906250005 | 3.0 | 4.0 | 8.054604317960672 | 0.06086840597050186 | 8.119333641975306 | 3.0 | 3.0 | S |
| 776 | 11.021124571436076 | 0.09411379177938939 | 1.0491734184027777 | 2.0 | 7.0 | 10.420501233277868 | 0.42230334816120857 | 2.0504648919753086 | 6.0 | 7.0 | A- |
| 777 | 10.551387840733 | 0.005626640816304781 | 0.918216444637346 | 3.0 | 4.0 | 9.891340702522346 | 0.035418420934798155 | 1.5642365933641977 | 2.0 | 6.0 | A- |

| | | | | | | | | | | | |
|---|---|---|---|---|---|---|---|---|---|---|---|
| 778 | 11.671534924016066 | 0.1404926511893542 | 1.6784641975308643 | 3.0 | 3.0 | 9.980992404272712 | 2.0262429244249764 | 8.018336111111111 | 3.0 | 3.0 | S |
| 779 | 12.910939358044425 | 0.09751385435096911 | 1.498924093364197 | 6.0 | 7.0 | 12.796662122463275 | 0.12197971491512723 | 1.7626250192901238 | 6.0 | 7.0 | A |
| 780 | 11.778139887778057 | 0.028597884209453025 | 1.1748753402777776 | 3.0 | 6.0 | 10.737693332891233 | 0.1753431937361176 | 3.4266123456790125 | 4.0 | 7.0 | A- |
| 783 | 11.136724158904979 | 0.007480507640776237 | 1.042639537229938 | 2.0 | 3.0 | 10.527137578750946 | 0.02567892159044366 | 1.9348037229938273 | 2.0 | 5.0 | A- |
| 784 | 11.718069027139666 | 0.1005255094264893 | 1.5635418402777779 | 4.0 | 6.0 | 10.990882002331722 | 0.412896305519022 | 3.5983982253086424 | 5.0 | 6.0 | A- |
| 788 | 10.957152436635626 | 0.025530249484409382 | 0.7033617777777782 | 2.0 | 3.0 | 8.107653000973423 | 0.01215951974568636 | 2.2360217777777778 | 3.0 | 2.0 | S |
| 789 | 10.850251963254221 | 0.007653375044104708 | 0.3463649452160494 | 3.0 | 3.0 | 10.595568306623326 | 0.03278240914499391 | 0.5553975624999998 | 6.0 | 6.0 | A- |
| 790 | 10.441964323610517 | 0.004848292431665883 | 0.7524598641975309 | 2.0 | 2.0 | 8.757251095010854 | 0.12174641698023841 | 2.09525625 | 2.0 | 7.0 | A- |
| 791 | 11.400027000849253 | 0.007348811781992533 | 0.7538821807484569 | 3.0 | 3.0 | 11.15320894446625 | 0.017141887454939192 | 1.043717640625 | 4.0 | 6.0 | A |
| 795 | 11.530365692650026 | 0.06027976769617885 | 1.0638776419753084 | 2.0 | 7.0 | 10.857905420940265 | 0.33244537067240787 | 2.5422530864197532 | 5.0 | 7.0 | A- |
| 799 | 11.077427722012272 | 0.014583419599542922 | 0.6380237501929011 | 2.0 | 3.0 | 10.97405602262563 | 0.020478849678476064 | 0.7360210069444444 | 2.0 | 4.0 | A |
| 800 | 11.53282938580094 | 0.07451017998587067 | 2.217824194637345 | 2.0 | 3.0 | 8.797256217509785 | 0.054944629934005444 | 12.49720696373457 | 3.0 | 3.0 | S |
| 802 | 12.391318187018987 | 0.08826514157374299 | 2.358784027777778 | 6.0 | 6.0 | 10.907527061079618 | 0.5678244931480655 | 10.33533196373457 | 7.0 | 7.0 | A- |

| | | | | | | | | | | | |
|---|---|---|---|---|---|---|---|---|---|---|---|
| 803 | 11.610922300412643 | 0.16400067387392098 | 1.4734587970679012 | 4.0 | 7.0 | 10.376728346212643 | 0.8572855883189454 | 5.056876562499999 | 5.0 | 6.0 | A- |
| 807 | 11.563593522829398 | 0.02435113410358018 | 1.2296345679012348 | 2.0 | 6.0 | 10.386521179185038 | 0.19998536603193495 | 3.9833506944444435 | 5.0 | 7.0 | A- |
| 809 | 13.502210247449966 | 0.02656459222343663 | 0.5374722656250002 | 6.0 | 6.0 | 13.403450368273157 | 0.04096155668701065 | 0.6630709184027778 | 6.0 | 6.0 | A |
| 811 | 14.686549160905042 | 0.04981550696576057 | 0.7715426406249999 | 4.0 | 7.0 | 14.651872331972573 | 0.05992676522693302 | 0.8986250017361112 | 4.0 | 7.0 | A |
| 812 | 13.11226980310909 | 0.030356880209856402 | 0.708169000771605 | 6.0 | 6.0 | 13.102884726504435 | 0.031913510276925613 | 0.729529515625 | 6.0 | 6.0 | A |
| 813 | 11.005205417529607 | 0.014189948208289923 | 1.5510972687114188 | 3.0 | 4.0 | 10.21254704784705 | 0.11133146730246866 | 3.5568912229938268 | 2.0 | 7.0 | A- |
| 814 | 13.422159692570391 | 0.050577388466589546 | 2.0864197530864197 | 3.0 | 6.0 | 12.76856629518445 | 0.07257567312139838 | 4.248751562499997 | 3.0 | 3.0 | F |
| 815 | 11.120399895988982 | 0.04745073269305944 | 2.328421673611111 | 3.0 | 3.0 | 8.024985715833052 | 0.04196976568702571 | 6.886323588155867 | 5.0 | 5.0 | S |
| 814_Dim | 12.564701065177957 | 0.12289112203424248 | 2.920501108033242 | 3.0 | 4.0 | 11.073014026890975 | 0.3637614882873661 | 10.730796675900278 | 4.0 | 4.0 | F_neg |
| 814_Bright | 13.53764628108355 | 0.16450081760843802 | 1.9999914127423832 | 2.0 | 3.0 | 13.459795590047815 | 0.23043439761477277 | 2.400053254847646 | 3.0 | 3.0 | F_pos |

**Table S3.** FACS measurements conducted in 2024 in sorted clone #814 $GFP_{Bright}$- and $GFP_{Dim}$ subpopulations, related to Fig. 2.

| Clone (#814 subpopulations) | Live count | (%) GFP(+) | (%) GFP(-) | GFP-fluo-gated | GFP-fluo-total | CV-gated | CV-total | Date |
|---|---|---|---|---|---|---|---|---|
| #814-$GFP_{Dim}$ | 3179 | 43.1 | 56.7 | 9010 | 4305 | 181 | 319 | 2024-04-19 |
| #814-$GFP_{Bright}$ | 3381 | 99.4 | 0.27 | 22755 | 23330 | 108 | 115 | 2024-04-19 |
| #814-$GFP_{Dim}$ | 2974 | 39.6 | 60.3 | 8958 | 3679 | 174 | 291 | 2024-04-23 |
| #814-$GFP_{Bright}$ | 3483 | 96.6 | 3.16 | 15126 | 14960 | 140 | 150 | 2024-04-23 |
| #814-$GFP_{Dim}$ | 3918 | 33.3 | 66.5 | 6333 | 2251 | 184 | 325 | 2024-04-26 |
| #814-$GFP_{Bright}$ | 4509 | 93.7 | 6.23 | 11083 | 10535 | 143 | 155 | 2024-04-26 |
| #814-$GFP_{Dim}$ | 5471 | 20.8 | 79 | 3706 | 918 | 161 | 335 | 2024-04-30 |
| #814-$GFP_{Bright}$ | 5873 | 87 | 12.9 | 7779 | 6808 | 151 | 165 | 2024-04-30 |
| #814-$GFP_{Dim}$ | 5393 | 21.2 | 78.7 | 4716 | 1139 | 168 | 358 | 2024-05-03 |
| #814-$GFP_{Bright}$ | 6201 | 86 | 13.9 | 8861 | 7687 | 148 | 166 | 2024-05-03 |
| #814-$GFP_{Dim}$ | 6211 | 19.4 | 80.5 | 3835 | 886 | 156 | 338 | 2024-05-07 |
| #814-$GFP_{Bright}$ | 6272 | 82.1 | 17.8 | 7632 | 6312 | 152 | 172 | 2024-05-07 |

**Table S4.** FACS measurements coupled with sRNAs and asRNAs measures conducted in 2024 across 9 sinpro clones in 6 subsequent time points, related to Fig. 3 and 4.

Excel file containing additional data too large to fit in a PDF.

**Table S5**. List of primers used in this study, related to Fig. 2, 3 and 5.

| Primer ID | Sequence | FW or RV | Description |
|---|---|---|---|
| HCP#213 | 5'-CTGATCTAGAGGTACCGGATCCTGAATTAGCCCTTCCAGTCC-3' | RV | RT-primer (plus Tag) sRNAs and sRNAs PCR amplification; strand-specific RT-qPCR version 1 |
| HCP#214 | 5'-CTGATCTAGAGGTACCGGATCCACCATGGTGAGCAAGGGCGA-3' | FW | RT-primer (plus Tag) asRNAs and asRNAs PCR amplification; strand-specific RT-qPCR version 1 |
| HCP#215 | 5'-CTGATCTAGAGGTACCGGATCCCGCCACAACATCGAGGACGG-3' | FW | RT-primer (plus Tag) asRNAs and asRNAs PCR amplification; strand-specific RT-qPCR version 1 |
| HCP#216 | 5'-AGCTGGAGTACAACTACAAC-3' | FW | sRNAs PCR amplification; strand-specific RT-qPCR version 1 |
| HCP#217 | 5'-TCAAGGATATCTTGTCTTCG-3' | RV | asRNAs PCR amplification; strand-specific RT-qPCR version 1 |
| HCP#237* | 5'-TGACAGCCGCCTAGCATTTC-3' | FW | PCR amplification on the HIV 5'LTR to verify the *in vitro* transcription product |
| HCP#238* | 5'-CCAGGCTCAGATCTGGTCTAAC-3' | RV | PCR amplification on the HIV 5'LTR to verify the *in vitro* transcription product |
| HCP#263 | 5'-CAGCTGCCTTGTAAGTCATTGG-3' | RV | asRNAs PCR amplification; strand-specific RT-qPCR version 2 & dPCR |
| HCP#264 | 5'-GGGATCACTCTCGGCATGG-3' | FW | RT-primer asRNAs and asRNAs PCR amplification; strand-specific RT-qPCR version 2 and dPCR |
| HCP#265 | 5'-CCTACACGACGCTCTTCCG-3' | FW | sRNAs PCR amplification; strand-specific RT-qPCR version 2 & dPCR |
| HCP#266 | 5'-CTCCCCCGCTTAATACTGACG-3' | RV | RT-primer sRNAs and sRNAs PCR amplification; strand-specific RT-qPCR version 2 and dPCR |
| HCP#267 | 5'-CAGTGAATCTTGGTTGTAAACTTGA-3' | FW | Homo sapiens TATA-box binding protein (TBP), transcript variant 1 and 2, mRNA targeting |
| HCP#268 | 5'-TCGTGGCTCTCTTATCCTCAT-3' | RV | Homo sapiens TATA-box binding protein (TBP), transcript variant 1 and 2, mRNA targeting |
| HCP#269 | 5'-AACATGCCATCCAGACTGAG-3' | FW | Homo sapiens ribosomal protein L27a (RPL27A), mRNA targeting |

| | | | |
|---|---|---|---|
| HCP#270 | 5'-GGTATTTGTCGAAGTTGATCCG-3' | RV | Homo sapiens ribosomal protein L27a (RPL27A), mRNA targeting |
| HCP#271 | 5'-[HEX]ATTGTGCCTGGCTAGAAGCACAAGAGGAGG[BHQ1]-3' | FW | Probe for asRNAs dPCR |
| HCP#272 | 5'-[FAM]AAAGCGAAAGGGAAACCAGAGGAGCTCTCT[BHQ1]-3' | FW | Probe for sRNAs dPCR |
| HCP#273 | 5'-TTTTTTTTTTTTTTTTTTTTTTTTTTTTTTTTTVV-3' | - | Oligo for poly(A) tailing |
| HCP#279 | 5'-[Biotin]TTTTTTTTTTTTTTTTTTTTTTTTTTTTTTTVV-3' | FW | PCR amplification on the *in vitro* transcription product (LIS-seq) |
| HCP#290 | 5'-CTTGTAGCACCATCCAAAGG-3' | RV | PCR amplification on the *in vitro* transcription product (LIS-seq) |
| HCP#304 | 5'-AATGATACGGCGACCACCGAGATCTACACTCTTTCCCTACACGACGCTCTTCCGATCT GGAGTGAATTAGCCCT-3' | FW | PCR amplification to prepare Illumina sequencing library (LIS-seq); PE1.0 sequence |
| HCP#305 | 5'-CAAGCAGAAGACGGCATACGAGAT[TCAAGT]GTGACTGGAGTTCAGACGTGTGCTCTTCCGATCTTTTTTTTTTTTTTTTTTTTTTT-3' | RV | PCR amplification to prepare Illumina sequencing library (LIS-seq); PE2.0 sequence; 6-nt index is written in the square bracket |
| HCP#364 | GACCAAAGAGCCCTATGATT | FW | PCR verification of the integration site of the provirus in clone #814; paired with the primer HCP#238* |
| HCP#365 | TCGTTTAGTCCTCTGGCAAC | FW | PCR verification of the integration site of the provirus in clone #814; paired with the primer HCP#238* |
